# A mosaic renal myeloid subtype with T-cell inhibitory and protumoral features is linked to immune escape and survival in clear cell renal cell cancer

**DOI:** 10.1101/2020.01.20.912865

**Authors:** Dorothee Brech, Tobias Straub, Evangelos Kokolakis, Martin Irmler, Johannes Beckers, Florian Buettner, Elke Schaeffeler, Stefan Winter, Matthias Schwab, Peter J. Nelson, Elfriede Noessner

## Abstract

Mononuclear phagocytes moderate tissue repair, immune activation and tolerance. In the renal tubulo-interstitium specialized dendritic cells help maintain homeostasis and protect tubuli from immune injury. Human renal cell carcinoma (RCC) is immunogenic; yet immunotherapies that target T-cell dysfunction show limited clinical efficacy suggesting additional mechanisms of immunoinhibiton. We previously described “enriched-in-renal cell carcinoma” (erc)DCs that are often found in tight contact with T cells which are dysfunctional. Here we describe that ercDCs exhibit a distinct polarization state imparted by tissue-specific signals characteristic for RCC and renal tissue homeostasis. The resulting mosaic transcript signature includes features associated with host defense activity, angiogenesis/invasion and T-cell inhibition. An ercDC-specific profile was predictive for patient survival and suggests potential therapeutic targets for improved immunotherapy.

**Significance:** Immunotherapies, which re-invigorate T-cell activity, achieve clinical responses in subsets of patients only revealing additional layers of T-cell inhibition. Mononuclear phagocytes can be immunoinhibitory. But, they are highly plastic and repolarization may be possible if key programming molecules can be identified, potentially enabling antitumor responses in tumors refractory to checkpoint blockade. We describe a myeloid cell type with mosaic feature including tumor-promotion and immunoinhibition in human clear cell renal cell carcinoma. Observed tight contacts with T cells may translate into T-cell dysfunction. A high ercDC score in tumor tissue correlates with poor patient survival suggesting ercDCs as targets for therapeutic intervention. Targeting molecules that are identified in the ercDC profile may expand the range of patients effectively treated by immunotherapy.

**Highlights:** Bullet points:

- Renal cell carcinoma (ccRCC) harbors polarized mosaic myeloid cells (ercDCs)
- ercDCs are found in contact with dysfunctional T cells in ccRCC
- ercDCs express novel immunoinhibitory proteins
- High ercDC z-score in ccRCC tissue correlates with poor patient survival

## Introduction

Dendritic cells (DCs) and macrophages of the mononuclear phagocyte system (MPS) are central players in the control of tissue homeostasis, wound healing and damage prevention. These processes are mediated in part through their capacity to remove cellular debris and induce angiogenesis, as well as by their ability to activate or induce tolerization of relevant immunocytes. The effector activities of MPS cells are strongly dependent on the cell’s polarization state which is imprinted by cues from the local environment (Das et al., 2015; Davies et al., 2013; Gordon et al., 2014).

In the murine kidney, the tubulointerstitial region is densely covered by myeloid cells expressing the kidney-specific homing receptor CX3CR1 (Engel et al., 2015; Hochheiser and Kurts, 2015; Soos et al., 2006). Due to their stellate morphology, these cells were designated DCs despite their dual expression of macrophage (F4/80) and DC markers (CD11c for murine DCs). The cells are thought to act as sentinels maintaining homeostasis and protecting the renal tubuli from immune-induced injury through tolerogenic mechanisms (Scholz et al., 2008). Clear cell renal cell carcinoma (ccRCC) arises from epithelial cells of the renal tubulointerstitium. This tumor type is generally richly infiltrated by immune cells, including T, NK and myeloid cells. However, despite abundance of CD8+ T cells that can recognize and destroy tumor cells when taken out of the ccRCC environment (Jantzer and Schendel, 1998; Leisegang et al., 2010), control of tumor progression fails, suggesting local suppression of T-cell effector activity (Giraldo et al., 2015; Giraldo et al., 2017; Prinz et al., 2012). Invigorating the T-cell response through blockade of the immune checkpoint molecule PD-1 has been approved as 2nd line treatment for metastatic RCC (Carlo et al., 2016; Motzer et al., 2015), but only a fraction of patients have been shown to respond. This suggests that mechanisms beyond those directly targeting the T cells help control the antitumor response, thus underscoring the critical need to better understand the tumor environment and to identify additional therapeutic targets beyond PD-1 to improve response rates and expand the range of patients that can be effectively treated.

We have previously shown enrichment of an unusual myeloid cells in patient ccRCC tissue that coexpress macrophage markers (CD14) and DC markers, i.e. CD209, which identifies them as a subset of interstitial DCs with crosspresentation ability (van Kooyk et al., 2013), as well as the costimulatory molecules HLA-DR and CD40. We designated the myeloid subpopulation “enriched-in-renal-carcinoma DCs” (ercDCs), due to its strong enrichment in the tumor center, lower abundance in the tumor periphery and even less presence in the non tumor-inflicted kidney area (Figel et al., 2011). We speculated that, analogous to the cell type described in murine kidney, ercDCs may also convey tolerogenic, tissue protective functions that shelter emerging tumor cells from immune attack. In ccRCC tissues, ercDCs were often found tightly engaged with T cells suggesting intercellular communication *in situ* (Figel et al., 2011). Through these observed T-cell contacts, we hypothesized that ercDCs could play a role in the suppressed T-cell function previously described (Prinz et al., 2012).

To help clarify the functional attributes of ercDCs and to position them within the MPS continuum, an analysis of the native cell population isolated from ccRCC tissue was performed. Transcriptomic profiling has been useful in defining subsets and polarization states within the MPS continuum (Hume and Freeman, 2014) and it has helped assign functional characteristics as demonstrated e.g. by Houser et al. who described two distinct subsets of CD14+ decidual marcophages, CD11cHI and CD11cLO, with distinct functions in tissue remodeling, growth and development (Houser et al., 2011).

In the present study we provide a molecular characterization of ercDCs isolated from ccRCC tissues. A definition of functional characteristics as well as their relationship to myeloid subtypes from other human tissues was established. The transcriptomic profiling identified ercDCs as a unique myeloid subset within the macrophage spectrum, and placed them in close relationship to inflammatory macrophages from the ascites of human ovarian cancer. The identified ercDC-specific gene expression profile was predictive for patient survival and suggests potential targets for therapeutic intervention that may help improve clinical efficacy of immunotherapy.

## Results

### The ercDC transcriptional profile identifies them as a unique myeloid subset within the macrophage spectrum

We recently reported that human ccRCC tissue harbor an unusual myeloid type, called ercDCs. ErcDCs resemble classical DCs by expression of costimulatory molecules (CD80, CD86, CD40), HLA-DR (Figure S1), and by their capacity to cross-present antigen. However, they also express both DC (CD209) and macrophage markers (CD14, CD163) (Figel et al., 2011). Here we report expression of additional surface markers that further highlight the unusual differentiation of this resident cell type (Figure S1). ErcDCs express CD141, a marker for crosspresenting myeloid cells, and CX3CR1, a homing receptor thought to regulate the abundance of DCs in the kidney (Hochheiser and Kurts, 2015, Hochheiser et al., 2013). Lack of CD83 and marginal levels of CCR7, but high levels of CCR1, CCR2 and CCR5 indicate an immature phenotype. Surface expression of other common myeloid markers, including CD68 and CD11b, is seen. In previous work we reported that ercDCs are tightly engaged with T cells in ccRCC tissue (Figel et al., 2011) and that the tumor-infiltrating T cells (TILs) are dysfunctional with features of cell cycle arrest and suppressed AKT/ERK pathway activation (Prinz et al., 2012). ErcDCs were also found to secret the matrix metalloproteinases, MMP2 and MMP9 as well as TNF (Figel et al., 2011), suggesting tumor-promoting characteristics in addition to direct effects on immune cells. To obtain a deeper functional understanding of ercDCs, in particular as it relates to a potential involvement in shaping the antitumor immune response, we performed an unbaised analysis.

CD14+CD209+ ercDCs and CD209-CD14+ macrophages were sorted from ccRCC tissue cell suspensions (Figure S2A, tables S1 and S2A) using flow cytometry and subjected to genome-wide gene expression analysis. Reference transcriptomes were generated from sorted blood monocytes (CD14+), slanDC and CD1c+ DC (all from PBMCs of healthy donors (HD)), and from *in vitro*-polarized M1- and M2-macrophages as described (Murray et al., 2014; Martinez et al., 2006) (Figure S2B, table S2A). CD1c+ DC and slanDCs were used as reference cell types for DC characteristics (Hansel et al., 2011; Nizzoli et al., 2013) to substitute for interstitial DCs, i.e. CD209+CD14-cells, which could not be sorted from ccRCC tissue cell suspensions due to low cell frequency.

Previously, using FACS analysis, it was difficult to assign ercDCs to either macrophages or DC subgroups as they co-expressed markers of both cell types, i.e. CD209, CD14, and co-stimulatory molecules CD80, CD86 and CD40. Analysis of their transcriptome with respect to core macrophage and DC genes (Gautier et al., 2012; Miller et al., 2012; Xue et al., 2014) now reveals that ercDC_ccRCC strongly express most of the human macrophage-associated core genes and show some expression of the established DC-associated core genes (Figure 1A). Protein expression analysis of selected macrophage (CD64A, CD14, CD32A, MerTK) and DC markers (ANPEP/CD13, FLT3) was confirmed by flow cytometry (Figure 1B, C). FLT3 or ANPEP/CD13, which are suggested DC-associated proteins (Xue et al., 2014), were not reliable for DC/macrophage distinction in our system (Figure 1C). For example, FLT3 discriminated between DCs and macrophages on the transcript level, but strong surface protein expression was seen on all cell types, CD1c+ DCs, M1- and M2-macrophages, and also ercDCs.

**Figure 1:**
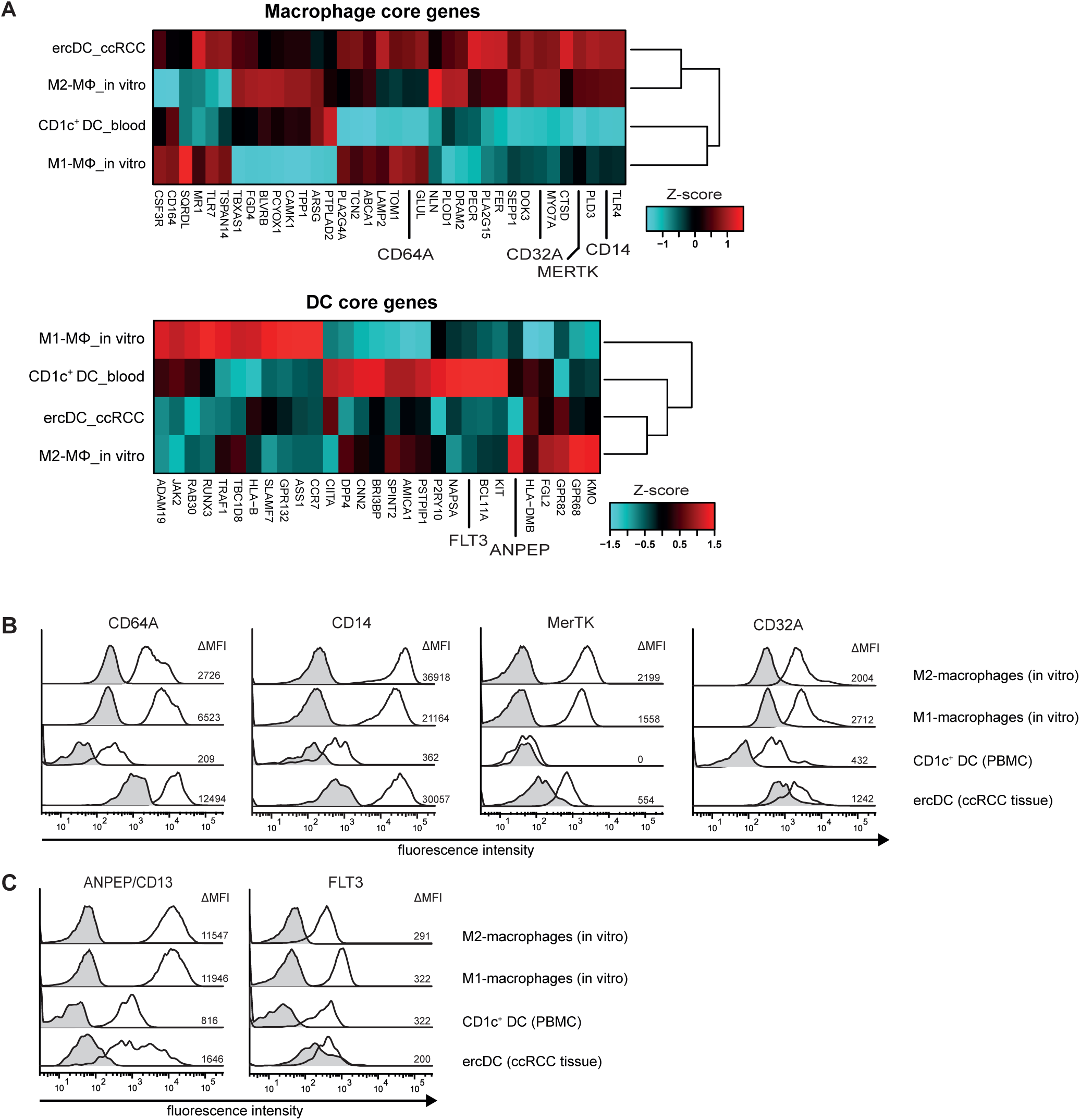
Human macrophage and DC core genes in ercDC_ccRCC. (A) Clustered heatmaps depicting relative gene expression levels of human macrophage and DC core genes comparing ercDC, *in vitro-*generated M1- and M2-macrophages, and CD1c+ DC from blood. Genes whose expression was validated on protein level (B, C) are set apart. (B), (C) Protein expression by flow cytometry. *In vitro*-generated M1- and M2-macrophages, PBMCs and ccRCC tissue cell suspensions were stained with marker combinations and gated on ercDCs (CD209+CD14+ cells among CD45+ live single CD3-CD11c+ cells of ccRCC cell suspension), CD1c+DCs (CD1c+cells among live single CD3-CD11c+ of PBMCs) and on M1- and M2-macrophage population among live single cells. Depicted are representative histograms of human macrophage markers CD64A, CD14, MerTK, CD32A (B) as well as DC markers ANPEP/CD13 and FLT3 (C) from at least 6 different patient tissues or PBMCs. Black line histogram: specific staining, gray filled histogram: control staining. Numbers indicate the control-corrected median fluorescence intensity.

A second MPS classification scheme based on transcription factors and growth factor receptors (Guilliams et al., 2014) showed robust expression of the macrophage-associated factors *MAF*, *MAFB*, *CREG1* and *CSF1R* by ercDC_RCC. CSF1R protein expression was validated by flow cytometry (Figure 2A). In contrast, ercDCs weakly expressed the DC-associated transcription factors *IKZF1*, *BCL6*, *IRF4* and the growth factor receptor *FLT3* (Figure 2B). Thus, using two different means of classification, ercDCs appeared to more closely represent a macrophage rather than a DC subtype. Macrophages represent a contiuum of different subtypes, wherein M1-(classical) and M2-(alternatively activated) macrophages are positioned at the opposing ends of the polarization spectrum (Mantovani et al., 2004; Sica and Mantovani, 2012). However, tissue macrophages are extremely heterogeneous and may adopt specialized functions as they respond to a variety of signals that change during homeostasis and inflammation (Gordon et al., 2014; Gordon and Taylor, 2005). A classification system based on the fundamental homeostatic macrophage activities – host defense, wound healing/tissue modulation and immunoregulation - has been established to help address this complexity (Mosser and Edwards, 2008). To position the ercDC_ccRCC within these classification schemes, we analyzed the transcriptome with regards to gene lists associated with biologic function (Gustafsson et al., 2008; Mosser and Edwards, 2008; Wang et al., 2012) (tables S3, S4, S5) including also an invasive signature gene list (Wang et al., 2012) which we supplemented with key angiogenic genes taken from the GSEA MSigDB database (table S6). The ercDC_ccRCC shared expression patterns with both M1- and M2-macrophages (Figure 2C, table S3, S4). Notable was the strong expression of the prototypic M1 gene *CD64A* (Mantovani and Sica, 2010; Tarique et al., 2015). However, ercDC_ccRCC also shared many markers with M2-macrophages, including *MerTK*, *CD204, CD206* and *CD36*. Validation of surface expression was performed for the M1-marker CD64A (Figure 1B) and M2-markers (CD204, CD206, CD36) (Figure 2C). Single cell flow cytometry using ccRCC tissue suspensions confirmed on protein level that the majority of CD14+CD209+ cells co-expressed CD64A (M1-marker) with MerTK and MSR1/CD204 (both M2-markers) (Figure 2D) supporting the suggestion that ercDCs might represent mosaic cells on the single cell level.

**Figure 2:**
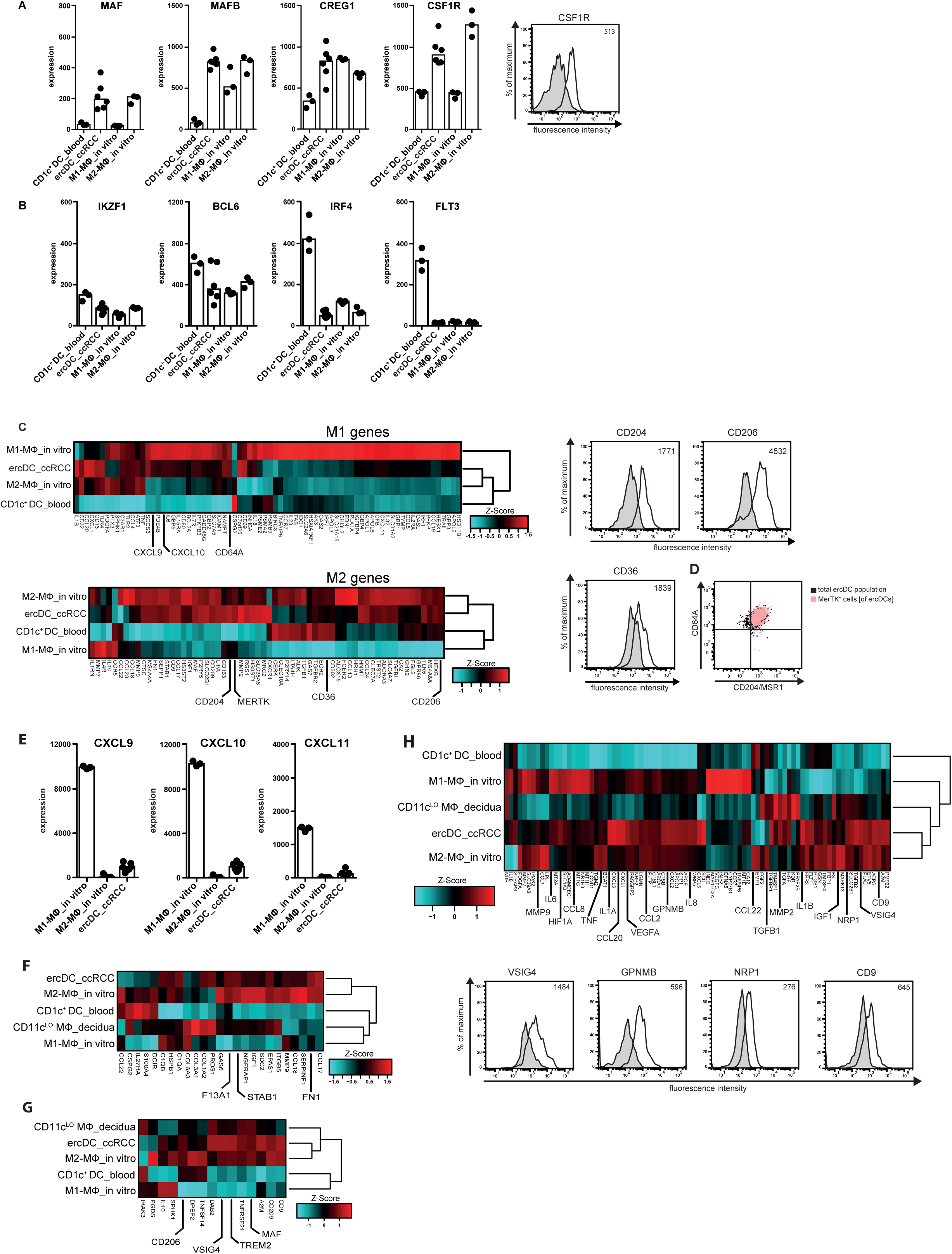
DC- and macrophage-associated genes and genes associated with angiogenesis and invasion in ercDCs. (A) Gene expression of macrophage-associated transcription factors and growth factor receptor CSF1R. Bars: median of each group, symbols correspond to individual array replicates of a cell type. Histogram depicts surface expression of CSF1R on ercDCs of ccRCC tissue cell suspensions by flow cytometry (B) Linear expression of genes for DC-associated transcription factors and growth factor receptor FLT3. (C) Clustered heatmaps depicts relative expression levels of genes associated with M1- and M2-macrophage polarization. Surface protein levels of indicated markers validated by flow cytometry. See table S3 and S4 for M1- and M2-genes. (D) Dot plot of polychromatic flow cytometry demonstrates co-expression of M1-(CD64A) and M2-macrophage markers (CD204/MSR1 and MerTK) on ercDCs in ccRCC tissue cell suspensions (n=4). Overlay of MerTK+ cells on gated CD64A+CD204+ double-positive CD209+CD14+ ercDCs (upper right quadrant) depicted in pink. (E) Linear expression of chemokines associated with host defense activity of macrophages. M1-macrophages are the positive reference. Bars: median of each group; symbols correspond to individual array replicates of a cell type. (F), (G) Clustered heatmap of relative expression levels of genes associated with wound healing and tissue remodeling (F) and immunoregulation (G). References are M2-macrophages and CD11cLO macrophages from decidua (CD11cLO MΦ_decidua). (H) Clustered heatmap depicting relative expression levels of genes associated with angiogenesis and invasion. Positive reference: *in vitro*-generated M2-macrophages; negative control: CD1c+ DCs from blood. Surface expression of indicated proteins on ercDCs by flow cytometry of ccRCC tissue-cell suspensions and gating on ercDCs (CD209+CD14+ cells among CD45+ live single CD3-CD11c+ cells). Histograms or dot plots are representative of at least n= 6) different tissue suspensions. Black line histogram: specific stainings, filled grey histogram: control stainings. Numbers indicate the control-corrected median fluorescence intensity.

ErcDC_ccRCC were found to express a number of chemokines (*CXCL9*, *CXCL10,* and *CXCL11*) that are characteristic of macrophage subtypes associated with host defense activity and help in the recruitment of Th1-polarized immune cells (Figure 2E). ErcDC_ccRCC were found to strongly express genes linked to wound healing and tissue remodeling (i.e. *STAB1*, *FN1, F13A*) (Figure 2F), thus resembling M2-MΦ_in vitro and CD11cLO MΦ_decidua, which have been linked to wound healing and tissue remodeling (Martinez et al., 2009; Martinez et al., 2008; Zhang and Mosser, 2008; Houser et al., 2011). The ercDC transcriptome appeared more similar to M2-macrophages with regards to immunoregulatory genes (i.e. *MAF*, *VSIG4*, *TREM2*, *CD206*) (Figure 2G) (Mosser, 2003; Pollard, 2009; Morris et al., 2011). In addition, ercDCs showed a strong signature of angiogenesis and invasion-associated genes. These included genes associated with the recruitment of proinflammatory monocytes, e.g. *CCL2*, *CCL8* and *NRP1,* as well as proinflammatory factors such as *TNF*, *IL6*, or *IL1B*. Genes involved in degradation of the extracellular matrix (e.g. *MMP2*, *MMP9*), hypoxia regulated genes (*HIF1A*) and proangiogenic genes (*GPNMB*, *VEGFA*, *IGF1*) were also part of the ercDC profile. Of note was the robust expression of *VSIG4* in ercDC_ccRCC. Surface protein expression of VSIG4, GPNMB, NRP1 and CD9 was validated by FACS (Figure 2H).

Collectively, these results indicated that ercDC_ccRCC combine features of various macrophage polarization states. High mRNA expression of *MMP2* and *MMP9* confirmed our previously described protein data and the angiogenic signature identified supports the hypothesis that ercDCs may help promote tumor growth (Figel et al., 2011). The strong expression of factors linked to the induction of T cell tolerance, *MAF* (Cao et al., 2005) and *VSIG4* (Vogt et al., 2006), support a potential immunoregulatory role for ercDCs in ccRCC tissue. VSIG4 was strongly expressed on the protein level, while other well described markers of immunoinhibition (PD-L1/B7-H1 and PD-L2/B7-DC) (Latchman et al., 2001; Saunders et al., 2005) were only marginally expressed. TIM-3 showed weak expression and B7-H3 was strongly expressed (Figure S1E).

### ErcDCs have a gene expression signature similar to inflammatory macrophages from ascites of ovarian cancer with characteristics of immune tolerance

Renal tubulo-interstitial DCs are described to act as sentinels maintaining homeostasis and protecting the renal tubuli from immune-induced injury through tolerogenic mechanisms (Scholz et al., 2008). To investigate whether ercDCs arising in the tubulointerstitial milieu of ccRCC similarly exhibit tolerizing features that might confer tumor immune protection we conducted a global analysis across published transcriptomes. The reference data comprised human myeloid cell types from blood and various non-lymphoid tissues, including myeloid subsets originating from tissues with described tolerogenic milieus (detailed information in supplemental procedures and table S2B). In brief, the subsets tested were CD11cHI and CD11cLO decidual macrophages (Houser et al., 2011), three DC subtypes from the lamina propria distinguished by their expression of CD103 and Sirpα (CD103+Sirpα+ DCs, CD103-Sirpα+ DCs, CD103+Sirpα-DCs) (Watchmaker et al., 2014), alveolar macrophages from non-smokers, smokers and COPD or asthma patients (Shaykhiev et al., 2009; Woodruff et al., 2005), human TAMs from gastrointestinal stromal tumors (GIST) (Cavnar et al., 2013), two myeloid cell types from the ascites of ovarian cancer patients (inflammatory macrophages (infMΦ_ascOvCa) and inflammatory DCs (infDC_ascOvCa)) (Segura et al., 2013), and CD141+ DCs from peripheral blood (Lindstedt et al., 2005). We supplemented the published data sets with in-house expression profiles of PBMC-derived monocytes, slanDCs and CD1c+ DCs.

Hierarchical clustering revealed similiarity of ercDC_ccRCC with MΦ_ccRCC, infMΦ_ascOvCa and infDC_ascOvCa (Figure 3A). CD11cLO MΦ_decidua, TAM_GIST, CD103+Sirpα+DC_gut and MΦ_asthma_avlung formed a separate subgroup. Blood-derived cell types, together with CD103+Sirpα-DC_gut and CD103-Sirpα+ DC_gut clustered distinct from all other cell types. CD141+ DC_blood more closely resembled CD103+Sirpα-DC_gut, confirming the similarity in characteristics previously described, including expression of markers associated with crosspresentation (Watchmaker et al., 2014). Principle component analysis (PCA) (Figure 3B) provided further evidence that ercDC_ccRCC are most similar to MΦ_ccRCC and infMΦ_ascOvCa, and are clearly different from blood-derived cells. To assess if expression states of specific genes can distinguish ercDC_ccRCC from other myeloid subtypes, we used the nearest shrunken centroids method (NSCM) (Tibshirani et al., 2002), a supervised machine learning approach suited to define subsets of genes that best characterize specific cellular states. Feature selection on the classifier that we trained to predict ercDC_ccRCC revealed 61 marker genes as predictive for ercDC (Figure S2C). Hierarchical clustering of all myeloid cell types based on expression of the 61 marker genes (Figure 4A) showed that ercDC_ccRCC clearly separated from the other human cell types analyzed with the exceptions of infMΦ_ascOvCa and CD11cLO MΦ_decidua. Despite originating from the same tissue, MΦ_ccRCC were positioned in a different cluster together with infDC_ascOvCa, and were thus not classified as ercDC_ccRCC. The tumor-associated macrophages from GIST (TAM_GIST), which are described as an antitumoral M1-like TAM subtype, did not show similarities with the ercDC_ccRCC profile. Blood-derived myeloid cells, slanDC_blood, CD1c+DC_blood and Mono_blood, exhibited the strongest differences to the ercDC_ccRCC profile, with an almost inverse expression of many of the marker genes.

**Figure 3:**
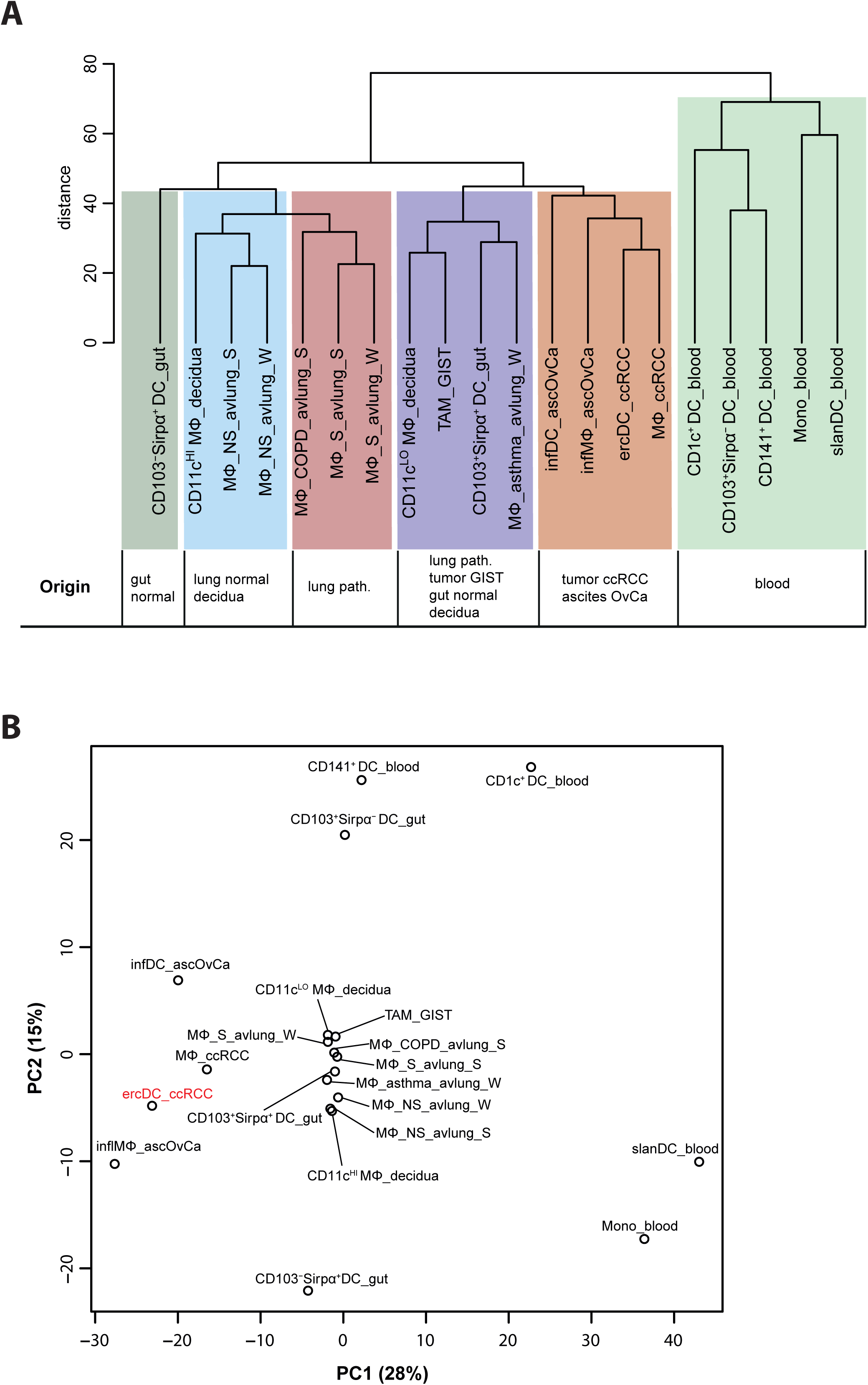
Transcriptome relationship of ercDC_ccRCC and myeloid cell types from blood and non-lymphatic healthy and pathological tissues. (A) Hierachical clustering of indicated cell types based on median expression values of replicate samples considering only the top 50% of genes with highest variation across cell types (6107 “informative” genes). (B) PCA on same data set as described in (A). Shown are the first two PC that describe 28% and 15% of the variance, respectively.

**Figure 4:**
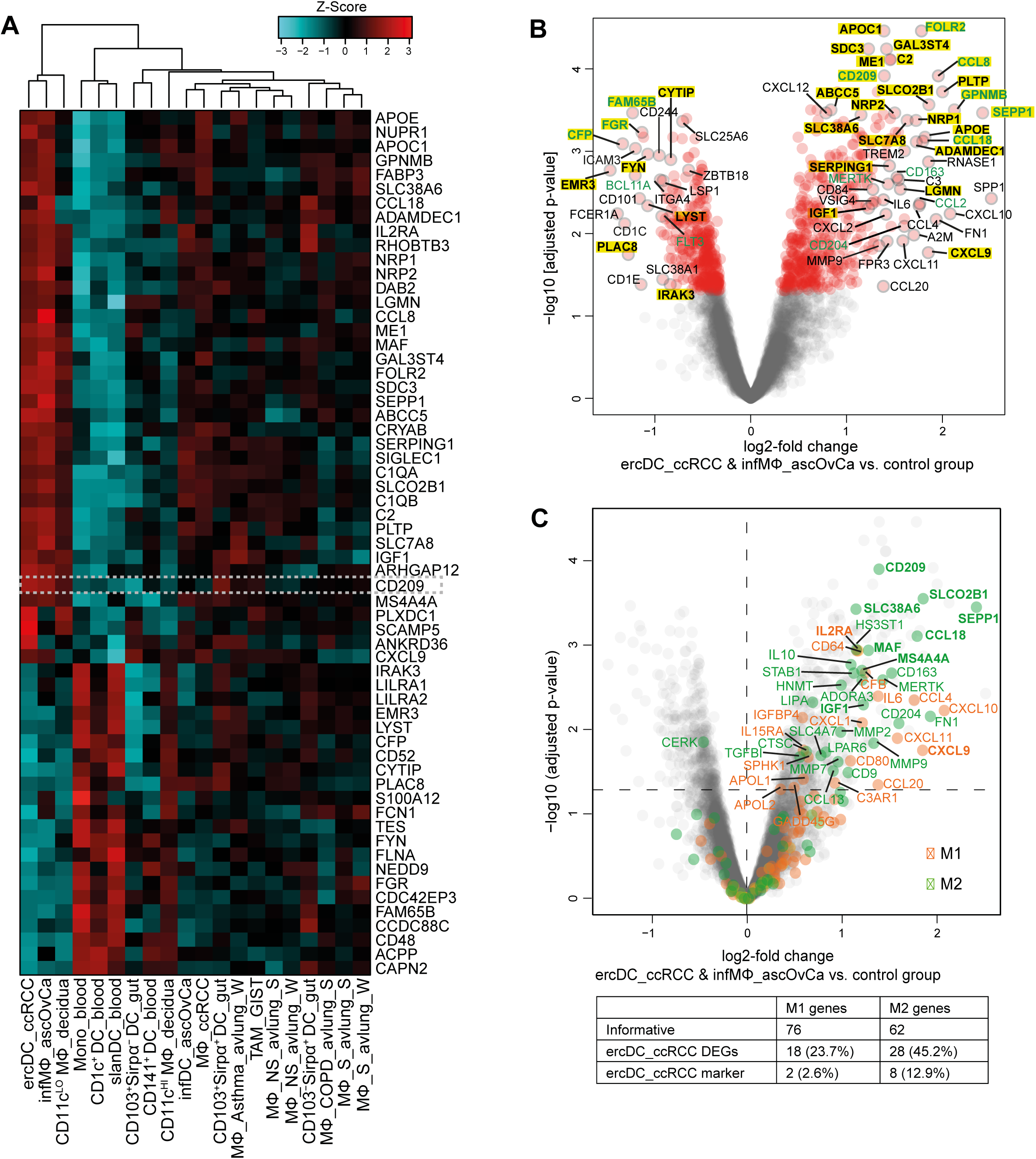
Marker gene profile and DEGs of ercDC_ccRCC. (A) Clustered heatmap of relative expression levels of 61 marker genes calculated with NSCM (see also Figure S2) comparing expression levels of ercDCs_ccRCC with those of all other cell types listed in tables S2 (control group). Dashed box highlights the gene CD209/DC-SIGN, which was used to distinguish ercDC_ccRCC from MΦ_ccRCC in FACS. (B) Volcano plot depicting differences in gene expression across informative genes of ercDC_ccRCC&infMΦ_ascOvCa (n=11) and control group (n=177). DEGs are red symbols (adjusted p < 0.05). Symbols with assigned gene name have a grey edge. ErcDC_ccRCC marker genes are in bold and highlighted in yellow; green: genes discussed in the text. (C) Volcano plot depicting M1- and M2-macrophage-associated genes (orange and green); (grey): informative genes resulting from the comparison of ercDC_ccRCC&infMΦ_ascOvca and the control group. DEGs among informative genes are located above the dotted horizontal line (p < 0.05). The table shows the number and percentage of ercDC_ccRCC&infMΦ_ascOvCa DEGs and marker genes (bold) within the informative M1- and M2-macrophage-associated genes.

Thirty-nine of the 61 marker genes were upregulated while 22 were downregulated in ercDC_ccRCC as compared to the other cell types (Figure 4A, table S7). As expected, CD209/DC-SIGN, which was used to distinguish ercDC_ccRCC from MΦ_ccRCC in FACS, was present in the marker gene list and showed increased expression (Figure 4A). The macrophage core gene *SEPP1*, the M2-associated gene *MAF* and the M1-associated gene *CXCL9* were present in the ercDC_ccRCC marker gene list, as well as genes associated with immunoinhibitory and proangiogenic functions, such as *GPNMB* and *NRP1*.

Given the strong similarity of ercDCs and infMΦ_ascOvCa we definded a list of differentially expressed genes (DEGs) between these two cell types and the other myeloid cell types. The genes with significantly different expression (FDR <0.05) in ercDC&infMΦ_ascOvCa as compared to the other samples (control group) are shown in Figure 4B and were designated ercDC_ccRCC DEGs. They include 788 genes, with 431 showing upregulation and 357 downregulation (tables S8, S9). The DEG list includes 54 of the 61 marker genes (89%), which are shown in bold letters with yellow background in the volcano plot (Figure 4B). Most of the marker genes showing the strongest and most significant expression differences were upregulated DEGs, e.g. *CD209/DC-SIGN*, *FOLR2*, *GPNMB* and *SEPP1*. The upregulated DEGs included genes associated with antiinflammatory function and macrophage recruitment. These include *CD204/MSR1*, *MERTK*, *CD163*, *CCL2*, *CCL8* and *CCL18* (Figure 4B). Downregulated DEGs included ercDC_ccRCC marker genes (*FAM65B*, *FGR, CFP*) and DC-associated genes, including *BCL11A* and *FLT3*.

Superposition of M1- and M2-associated genes (table S3, S4) on the list of informative ercDC_ccRCC genes again illustrated the mosaic expression of M2- and M1-associated genes by this myeloid subtype (Figure 4C). Among the most significantly upregulated genes were the M2-associated genes, *CD209*/*DC-SIGN, SLCO2B1, SLC38A6, SEPP1, CCL18, MAF*, *MS4A4A* and *IGF1*, but also the M1-associated genes *IL2RA* and *CXCL9*. These M1/M2-associated genes also belonged to the ercDC_ccRCC marker genes. Overall, 2.6% of the M1-associated genes and 12.9% of the M2-associated genes were among the ercDC_ccRCC marker genes. Moreover, 23.7% of M1-associated genes and 45.2% of M2-associated genes were part of the ercDC_ccRCC DEGs. GSEA analysis provided evidence of significance for the difference in expression of the M2-gene set (p=0.02) and enrichment of the M1-gene set (p=0.09) in the ercDC_ccRCC&infMΦ_ascOvCa. This suggests that ercDC_ccRCC may represent a hybrid myeloid subtype with mosaic features of M2- and M1-polarization.

To identify enriched biological processes associated with the DEGs, GO term analysis was employed. The results showed “response to wounding” and “inflammatory response” as the most significantly scored categories (Figure 5A). These effector processes correspond with the described inflammatory milieu in RCC (Dvorak, 1986; Fox et al., 2013) and the well recognized role for renal macrophages in normal tissue homeostasis and wound healing (Nelson et al., 2012; Kawakami et al., 2013). The third most enriched category was “defense response”, underscoring a potential bactericidal activity of ercDC_ccRCC, which agrees with the general expression of M1-associated genes.

**Figure 5:**
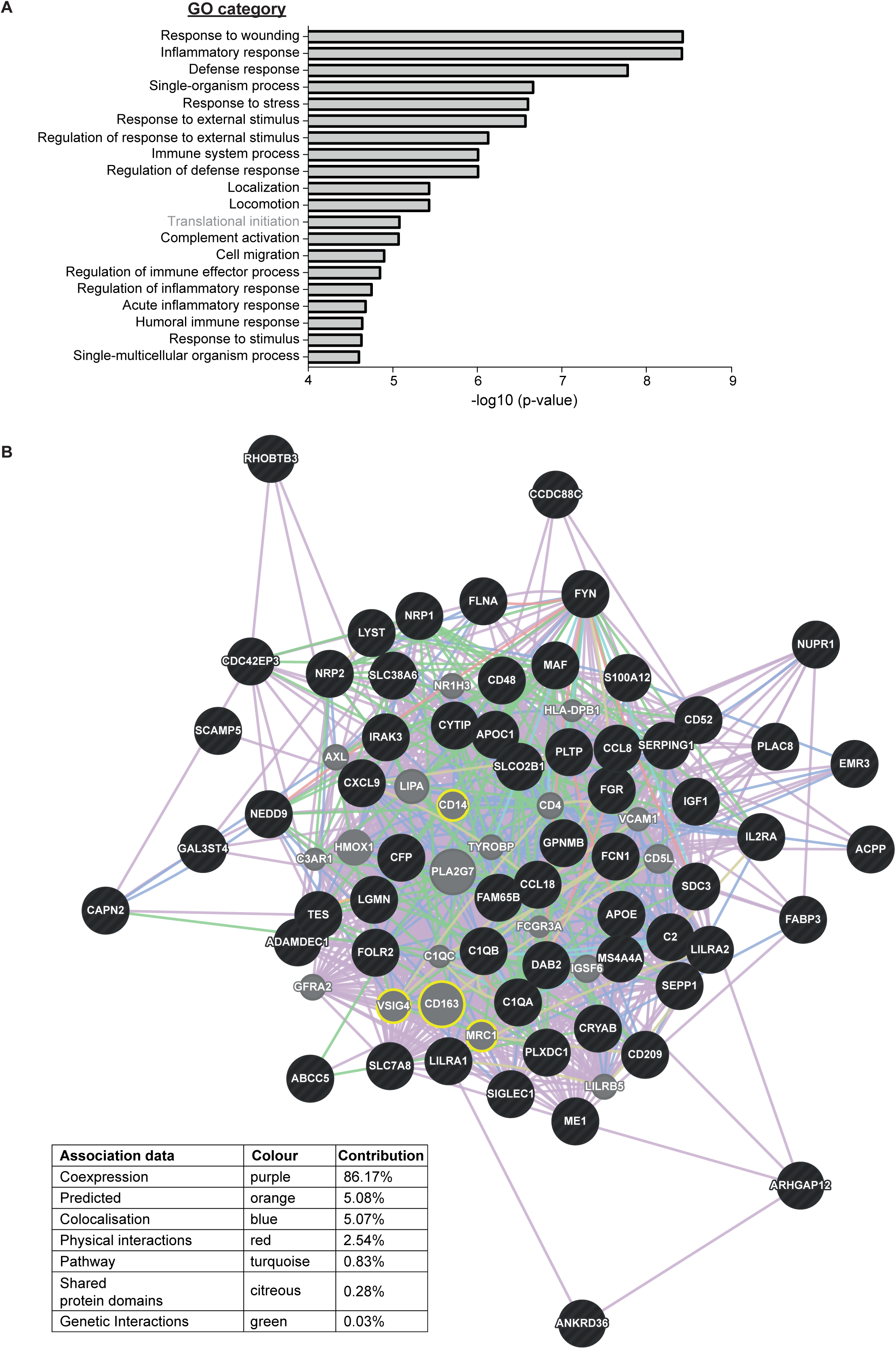
GO categories and functional network of ercDC_ccRCC. (A) Upregulated DEGs were enriched in 19 of the top 20 categories, downregulated DEGs were only enriched in one category (“Translational initiation”) of the top 20 significantly enriched terms. Significance of GO category enrichment is depicted as -log10 (p-value). (B) Functional associations of ercDC_ccRCC marker genes (black) and their computed related genes (grey) by GeneMANIA network analysis. Yellow circles highlight related genes mentioned in the text. Associations were defined based on different criteria (coloured lines, see Figure).

The results of InnateDB and GSEA (based on the data bases “KEGG”, Reactome” and “Biocarta”) analyses overlapped with GO term analysis (table S10). Most of the enriched pathways identified are associated with the complement system, lipid metabolism and modulation of the extracellular matrix. Deregulated lipid metabolism has been described to promote the development and progression of RCC (Drabkin and Gemmill, 2010).

GeneMANIA network analysis was conducted to better characterize the functional network of ercDC_ccRCC marker genes, and to identifiy functionally related genes (Figure 5B). The related genes identified (grey) included many macrophage-associated genes, particularly those of M2-macrophages and their immunoinhibitory function, *CD163*, *CD206*/*MRC1*, *CD14* and *VSIG4* (yellow circles in Figure 5B).

### ErcDCs are distinct from blood-derived monocytes from RCC patients

Recently it was reported that blood-derived monocytes from RCC patients display a tumor-promoting transcription profile (Chittezhath et al., 2014). Inflammatory blood monocytes can act as precursors for TAMs (Caso et al., 2010; Movahedi et al., 2010; Qian et al., 2011). Hierarchical clustering and PCA evaluation including the RCC-monocyte transcriptome (designated as Mono_RCC_blood) clearly separated the Mono_RCC_blood from the ercDC_ccRCC and clustered the Mono_RCC_blood with CD11cHI MΦ_decidua and CD103-Sirpα*+* DC_gut (Figure 6A, B).

**Figure 6:**
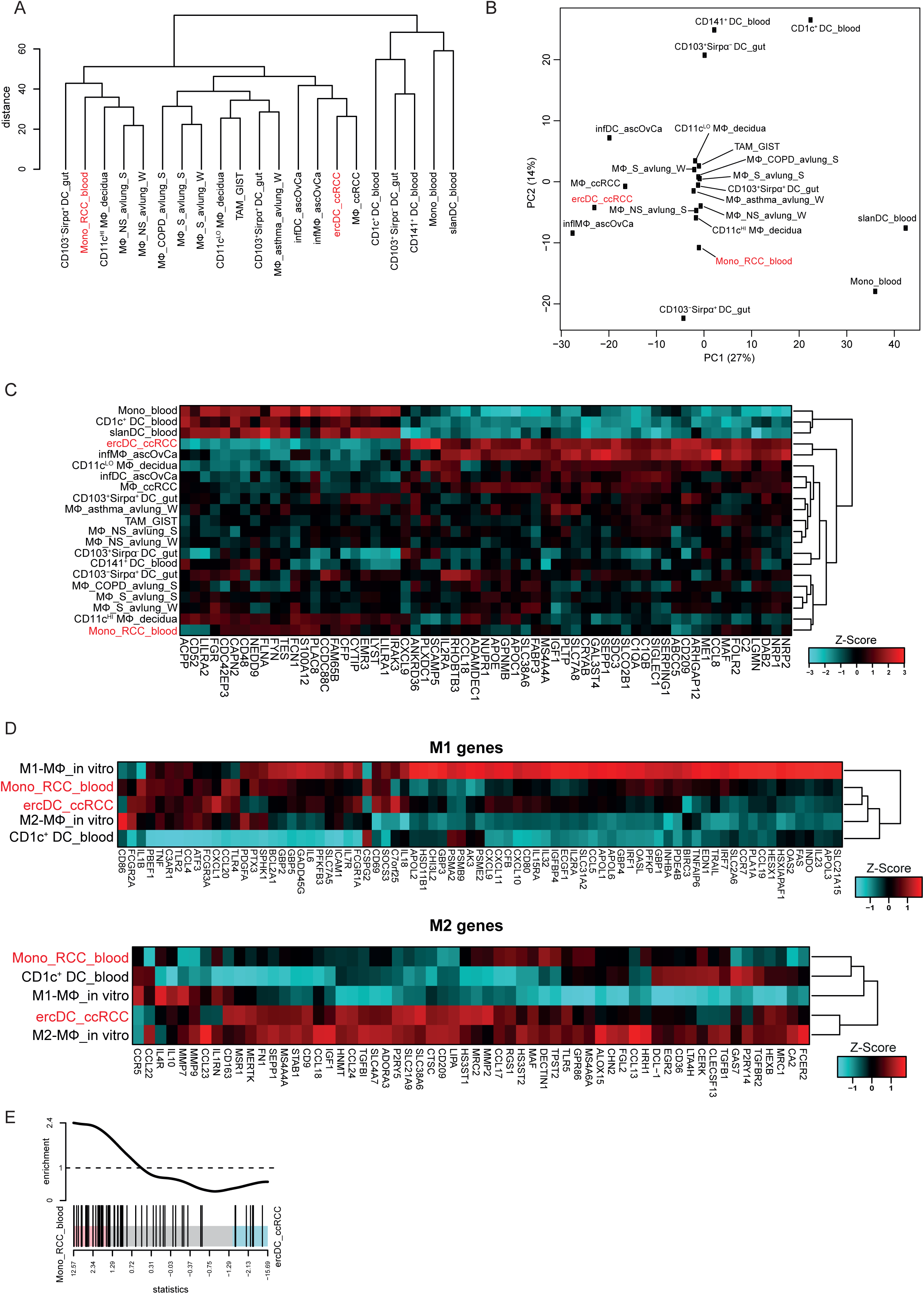
Transcriptome comparison of ercDC_ccRCC with published blood monocytes from RCC patients. All figures are based on expression levels with blood-specific genes not excluded. See Figure S3 for results after exclusion of blood-specific genes. (A) Hierachical clustering and (B) PCA of gene expression profiles. Analyses are based on informative genes for which the median expression value of replicates for each cell type was calculated. (C) Clustered heatmap of ercDC_ccRCC marker genes with blood monocytes of RCC patients (Mono_RCC_blood). (D) Clustered heatmaps showing relative expression levels of genes associated with M1- and M2-macrophages in Mono_RCC_blood and ercDC_ccRCC. (E) Enrichment plot for 73 IL-1R pathway genes defined by merging MSIGDB sets “BIOCARTA_IL1R_PATHWAY”, “REACTOME_IL1_SIGNALING”, “PID_IL1_PATHWAY” and “INTERLEUKIN_1_SECRETION”. Genes indicated by vertical bars are spread along the x-axis based on t-statistic comparing ercDC_ccRCC and Mono_RCC_blood cells. Enrichment worm on top shows the relative enrichment of genes in each part of the plot.

Tissue-specific gene expression can obscure cell type-specific profiles. However, even after exclusion of blood-specific genes (“tissue preferential expressed gene list (blood genes)” (Mele et al., 2015)) Mono_RCC_blood remained clearly distinct from ercDC_ccRCC and retained their similarity with the CD11cHI_MΦ_decidua and CD103-Sirpα*+* DC_gut (Figure S3A, B). Clustering based on the ercDC_ccRCC marker genes positioned the Mono_RCC_blood distant to ercDC_ccRCC in a subcluster together with CD11cHI MΦ_decidua (Figure 6C), similar to the hierarchical cluster analysis on all informative genes (Figure 6A). Identical clustering was observed after exclusion of blood-specific genes (Figure S3C).

A comparison of the expression of M1- and M2-associated genes between RCC-monocytes and ercDCs revealed that Mono_RCC_blood resembled ercDC_ccRCC in their expression of M1-associated genes, but were more similar to CD1c+_DC_blood in their expression of M2-associated genes, while ercDC_ccRCC clustered with M2-macrophages (Figure 6D). Removal of the blood-specific genes again did not change this profile (Figure S3D). Mono_RCC_blood are thought to derive their protumoral activity through an interleukin-1 receptor (IL-1R)-dependent mechanism. Comparing the IL-1R pathway activation by GSEA showed that the corresponding genes were less expressed (p=0.0001) in ercDC_ccRCC as compared to Mono_RCC_blood (Figure 6E).

### Expression of ercDC marker genes in ccRCC tissue is predictive of patient survival

We have previously shown that CD209+ cell numbers were higher in advanced ccRCC tumors with poor prognostic tumor stage (Figel et al., 2011). The transcriptome analysis described here further supports a role of ercDCs in tumor promotion and immunoinhibiton. As shown in Figure 7, an ercDC score based on the expression of the ercDC marker genes was found to be associated with cancer-specific survival (CSS). The association was significant for the large Cancer Genome Atlas (TCGA) cohort of 442 ccRCC samples (HR=1.8, p=3.0E-02) and an independent cohort of 28 ccRCCs (Rostock cohort) (table S12) (HR=4.8, p=8.8E-03). In both cohorts, patients with high ercDC score showed decreased CSS as compared to patients with low ercDC score. In addition, the ercDC score correlated with tumor grade (Figure 7B, D) in both ccRCC cohorts.

**Figure 7:**
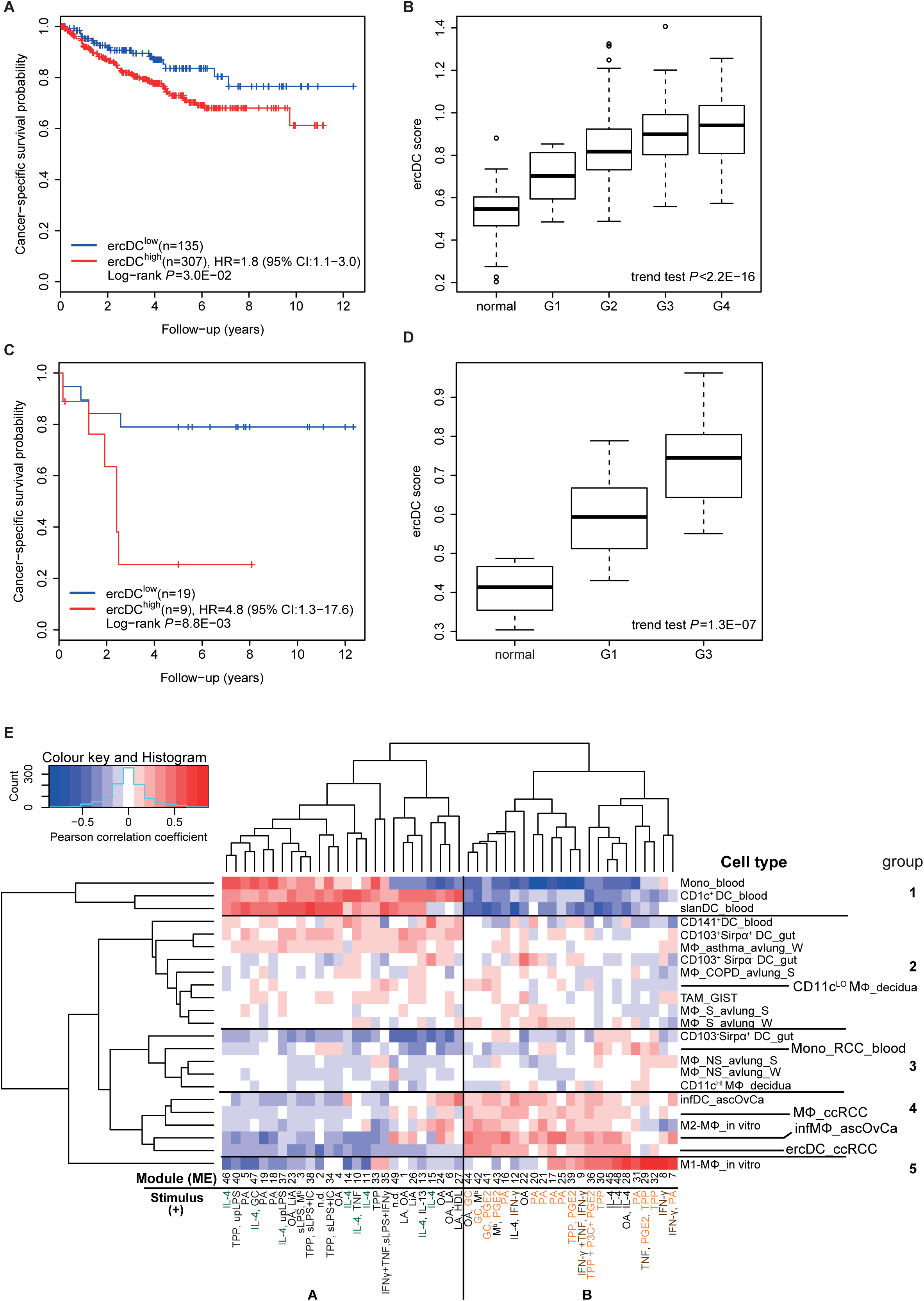
Association of ercDC_ccRCC marker gene profile with cancer-specific survival and specific polarization stimuli. (A) (C) Kaplan-Meier estimates of cancer-specific survival (CSS) of ccRCC tumors from the TCGA cohort (n = 442) and the Rostock cohort (n = 28) predicted by the ercDC score. Partitioning into ercDClow and ercDChigh using conditional inference trees with endpoint CSS. (B) (D) Box plots of ercDC score values of TCGA ccRCC tumors (n = 439, no grading was available for 3 samples of TCGA cohort, see table 12) and normal kidney tissues (n = 65) or the Rostock tumors (n = 14 for grade G1, G3, and normal kidney tissues) (p-value by linear trend test). (E) Heatmap of Pearson correlation coefficients calculated between module eigengenes (ME) and myeloid cell types. Cell types were clustered based on correlations with stimuli-associated eigengen modules. Depicted are the numbered modules with up to two significantly positive correlated stimuli for each module. Stimuli dominating in module group A are highlighted green, in module group B in orange and brown. Stimuli negatively correlating with the modules as well as the correlation rules for stimuli and modules are listed in table S11.

### The ercDC_ccRCC polarization profile reveales distinct tissue imprints

Recent reports have suggested that the phenotype and function of tissue-resident macrophages is robustly influenced by factors present in the tissue micromilieus (Gordon et al., 2014; Davies et al., 2013; Das et al., 2015). To investigate the potential influence of the ccRCC milieu on the gene expression profile of ercDC_ccRCC, predefined stimulus-specific human gene sets were used that are known to be induced by distinct macrophage activation signals (Xue et al., 2014). We correlated the module eigengenes (ME) of the stimulus-specific gene sets (so-called modules) with the gene expression profile of ercDC_ccRCC, various human myeloid cell types from non-lymphoid tissues, including Mono_RCC_blood, and *in vitro*-generated M1- and M2-macrophages (Figure 7E, table S11). The ercDC_ccRCC clustered closest with infMΦ_ascOvCa, confirming the strong relationship seen previously using the NSCM. Macrophages from the ccRCC tissues (MΦ_ccRCC), M2-MΦ_in vitro and infDC_ascOvCA clustered within the same group (no. 4). All cell types from this group also showed largely negative correlations with modules in cluster A, linked to signals associated with the cytokine IL-4. On the other hand, they showed predominantly strong positive associations with modules in cluster B. The modules of cluster B were generally linked to signals from glucocorticoids (GC), palmitic acid (PA), prostaglandin E2 (PGE2) or a combination of TNF, PGE2 and P3C (Pam3CysSerLys4, TLR2-ligand, TPP) (in orange letters). Other modules in cluster B are linked to M1-polarizing stimuli (IFN-γ, TNF) (in brown letters); accordingly, a very strong positive correlation was found with *in vitro*-polarized M1-macrophages (M1_MΦ_in vitro) and a negative correlation with M2_MΦ_in vitro. ErcDC_ccRCC also correlated negatively or showed no correlation, while infMΦ_ascOvCA, infDC_ascOvCA and MΦ_ccRCC showed some positive correlation. Overall, Mono_RCC_blood showed much weaker polarization than ercDC_ccRCC, illustrating that the two cell types have different tissue origins.

In summary, the data suggest that the characteristic transcriptional profile of ercDCs may be induced in part by GC, PGE2, PA and TPP, and absence of IL-4. This is consistent with described characteristics of the RCC tissue milieu, featuring GC (Huynh et al., 2015), PGE2 (Wu et al., 2011) and TNF (Gogusev et al., 1993; Chuang et al., 2008) and lacking IL-4 (Hosse, 2009). PA has been described as playing a role in kidney fibrosis (Kang et al., 2015). The emerging imprinting stimuli are also consistent with the marker profile determined using NSCM and the upregulated DEGs. PGE2, for example, is described to induce the transcription of *MSR1*/*CD204* (Domingo-Gonzalez et al., 2013), a gene also strongly expressed in ercDC_ccRCC. Moreover, PGE2 and GC are known inducers of *CD163* (Schaer et al., 2002; Zhang et al., 2014), another marker strongly upregulated in ercDC_ccRCC. In addition, GC regulates the expression of *MERTK* as well as *C1QB*, *CCL8*, *VSIG4* and *FCN1*, all of which belong to the ercDC_ccRCC marker genes or DEGs.

### VSIG4 on ercDCs and contacts with FOXO1-positive T cells *in situ*

Based on the observation that a high ercDC score is associated with poor patient survival, we then examined the transcriptional profile of ercDCs for genes linked to immune evasion and tumor progression in more detail. In particular, *VSIG4* appeared among the immunoregulatory and proangiogenic/invasive features and emerged as a gene functionally related to the marker genes in the GeneMANIA network analysis. ErcDC_ccRCC strongly expressed *VSIG4* at the RNA (Figure 8A) and protein level at the cell surface (Figure 8B, C). Multiparameter immunofluorescence histology of ccRCC tissues showed that the majority (85-98%) of CD209+ cells coexpressed VSIG4. Tumors from advanced tumor stages harbored more CD209+VSIG4+ cells than those of earlier tumor stages (abs. median number 45.7, range 21-75; vs 35, range 31-58). We have previously reported that ercDCs are in tight contact with T cells in ccRCC tissue (Figel et al., 2011). We had additionally observed that the T cells from ccRCC tissue are functionally inactive with a molecular signature of anergy, including suppression of ERK and AKT pathways and high expression of cell cycle inhibitor p27kip (Prinz et al., 2012). We therefore asked whether the cell cycle arrest in T cells could be caused through T-cell contact with ercDCs expressing VSIG4, thereby inducing p27kip. As p27kip is a downstream target of FOXO1, which is a transcription factor that is negatively regulated by AKT, we evaluated the expression of VSIG4 and FOXO1 in ercDC-T cell contacts *in situ* using immunohistology. We observed that many T cells (12-41%) in ccRCC tissue stained positive for FOXO1. FOXO1 was localized to the nucleus indicating an active protein (Figure 8D), which is consistent with suppressed AKT pathway in ccRCC-resident T cells. FOXO1 was preferentially expressed in T cells that interacted with VSIG4+ ercDCs compared to T cells not in contact with VSIG4+ ercDCs (Figure 8E). In late stage tumors (T105, T114, T118), the majority of the FOXO1+ T cells were found in contact with VSIG4+ cells contrasting early stage tumor (T90).

**Figure 8:**
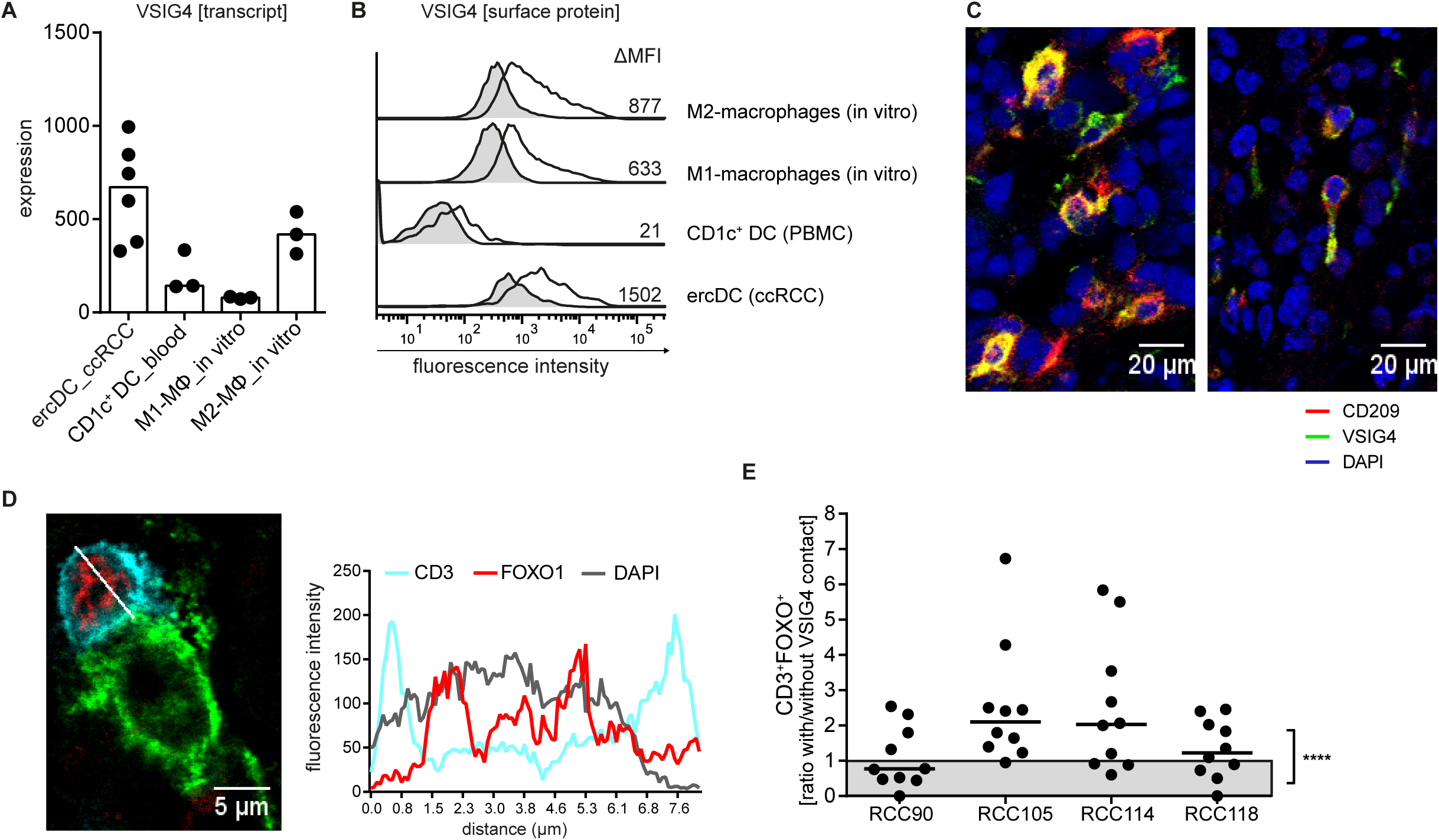
VSIG4 on ercDCs and contacts with FOXO1-positive T cells. (A) Linear expression of *VSIG4*. Bars are the median of each group; symbols correspond to individual array replicates of a cell type. CD1c+DC_blood is the negative reference, M2-MΦ_in vitro the positive reference. (B) Validation of VSIG4 protein on ercDCs in ccRCC tissue cell suspension by flow cytometry. ercDCs are gated as CD209+CD14+ cells among CD45+ live single CD3-CD11c+ cells in ccRCC cell suspension, CD1c+DCs are gated within PBMCs as CD1c+ cells among live single CD3-CD11c+ cells. Control staining: grey filled histogram. Numbers depict the difference in median fluorescence intensity (MFI) between specific antibody and control. Shown is a representative example of n = 4 different ccRCC tissue cell suspensions. (C) Confocal images of ccRCC tissues stained with CD209 (red), VSIG4 (green) and DAPI (blue). Original magnification, x400. Cells coexpressing CD209 and VSIG4 are yellow. Images are merged fluorescent channels and maximal projection of 6-9 z-planes (z-step size = 0.7 µm). Left image: late stage ccRCC (RCC114, T3cG2), right image: early stage ccRCC (RCC90, T1aG2), (D). Confocal image of an ccRCC tissue area (RCC118) depicting a T cell (CD3+, cyan) expressing FOXO1 (red) in contact with VSIG4+ cell (original magnification, x400, zoom 5). Histograms document fluorescence intensities of a single z-plane along indicated line. (E) Scatter plot showing the ratio of the % FOXO+ T cells among T cells that are in contact with VSIG4+ cells to the % of FOXO+ T cells among T cells that are not in contact with VSIG4+ cells. Values above >1 indicate that the majority of CD3+FOXO+ cells is in contact with VSIG4+ cells. **** = p < 0.001, Mann Whitney test comparing across all 4 ccRCC tissues.

## Discussion

Renal cell carcinoma (RCC) has long been recognized as an immune-responsive tumor with a well documented sensitivity to T-cell attack dating from the 1970s (Kim et al., 2003; Motzer, 2003; Vogelzang and Stadler, 1998). Yet, clinical studies using to immune checkpoint blockade therapy that releases T cells from tumor-induced inhibition by targeting the co-inhibitory protein CTLA-4 or the PD-1/L1 axis (Sharma and Allison, 2015) document that RCC patients do not benefit substantially more compared to patients with tumors previously considered to be non-immunogenic (Sunshine and Taube, 2015). This observation indicates the existence of additional layers of tumor-mediated immunosuppression beyond T cell checkpoints that hamper antitumor immune responses and need to be addressed to improve efficacy of cancer immunotherapy.

We have previously shown that T cells infiltrating human ccRCC display a signature of cell cycle inhibition with suppressed ERK and AKT pathway activation (Prinz et al., 2012). Many of the T cells seen in ccRCC tissue are tightly engaged with CD209/CD14 double-positive myeloid cells (Figel et al., 2011). Here we provide evidence for tumor-promoting qualities as well as immunoinhibitory characteristics of ercDCs that may translate to T-cell dysfunction. A high ercDC score was found to be strongly associated with poor patient survival, showing prognostic value and suggesting that ercDCs may be targets for therapeutic intervention. Targeting ercDCs, particularly in situations, where tumors have high ercDC content, may expand the range of patients that can be effectively treated by immunotherapy.

The ercDC transcriptome revealed details regarding their functional polarization and positioning within the MPS. The transcriptomic data suggest that ercDCs belong to the macrophage lineage as they strongly expressed human macrophage-associated core genes, transcription factors and growth factor receptors. Notably, they show combined features of M1-macrophages and M2-macrophages as well as gene signatures associated with wound healing and tissue remodeling, immunoregulation, and bactericidal effector activities. These observations were supported by GO term, InnateDB and GSEA analyses, showing “response to wounding”, “inflammatory response” and “defense response” as the most significantly enriched categories. Furthermore, pathways associated with the complement system, lipid metabolism and modulation of the extracellular matrix were highly enriched. These features are strongly associated with the inflammatory milieu in RCC tissue (Dvorak, 1986; Fox et al., 2013) and the general role of macrophages in tissue homeostasis and wound healing in kidney biology (Kawakami et al., 2013; Nelson et al., 2012). An altered lipid metabolism is a characteristic of ccRCC (Pinthus et al., 2011; Rezende et al., 1999) as is the accumulation of immune complexes with accompanying complement activation, which is also seen in many inflammatory renal diseases (Bagavant and Fu, 2009; Nikolic-Paterson and Atkins, 2001). The identification of these complex features for a myeloid cell subtype enriched in RCC confirms and extends previous studies that have reported diametrically polarized macrophages in RCC tissue (Kovaleva et al., 2016).

A comparative analysis across published transcriptome databases of human myeloid cell types from blood and non-lymphoid tissues using the NSCM method showed an association with an inflammatory macrophage subtype from the ascites of ovarian cancer (infMΦ_ascOvCa) as closest relative to ercDC_ccRCC. Described features for ovarian TAMs include the expression of M2-markers (CD14, CD206, CD11b, CD204), select M1-markers (CD86, TNF) (Allavena et al., 2010; Bellora et al., 2014; Colvin, 2014; Kawamura et al., 2009; Merogi et al., 1997; Segura et al., 2013), and expression of the immunosuppressive and antiinflammatory chemokine CCL18. These expression patterns parallel those seen in ercDC_ccRCC. Interestingly, cytokines shown to induce the ercDC phenotype *in vitro*, IL-6, CXCL8/IL-8 and VEGF (Figel et al., 2011), are also present in ovarian ascites (Giuntoli et al., 2009; Lane et al., 2011; Reinartz et al., 2014). Thus, the relatedness of ercDC_ccRCC and infMΦ_ascOvCa may be explained in part by their exposure to a related tissue milieu. In addition, PGE2, GC, TNF, palmitic acid (PA) and TLR2-ligands were identified as ercDC_ccRCC polarizing factors. PGE2 is an important factor in RCC biology (Wu et al., 2011). The enrichment in PGE2 and GC is linked to the expression of CD163 (Schaer et al., 2002; Zhang et al., 2014) in ercDCs. In addition, GC has been described to induce MerTK (Zizzo et al., 2012) and VSIG4 (Gorgani et al., 2011), and as such, may be an underlying mechanism for the expression of those markers in ercDCs.

By hierarchical clustering of the transcriptome data of ercDC_ccRCC and other myeloid cell types, ercDCs clearly separated from all blood-derived myeloid cells, including blood-derived monocytes from RCC patients, which were described to potentially represent precursors of RCC-TAMs (Caso et al., 2010; Chittezhath et al., 2014; Movahedi et al., 2010; Qian et al., 2011). Hierarchical clustering and classification based on the ercDC_ccRCC marker genes similarly identified the Mono_blood_RCC as blood-derived myeloid cells with a transcriptome distinct from tissue-derived cell types including ercDCs_ccRCC, even after elimination of blood-specific genes. Based on the results we conclude that ercDCs represent a cell type that is resident to non-lymphoid tissues displaying a distinct polarization state imparted by tissue-specific signals. Indeed, we were not able to detect CD209+CD14+CD163+ myeloid cells among peripheral blood mononuclear cells by FACS (data not shown) and the ercDCs score was strongly decreased in samples of acute myeloid leukemia of the TCGA collection compared to ccRCCs (not shown).

Several genes within the ercDC marker profile suggest T-cell inhibitory effector activity. While PD-L1, PD-L2 and TIM-3 were only marginally expressed by ercDCs, prominent cell surface expression was found for the immunoinhibitory proteins VSIG4 (He et al., 2008; Ikarashi et al., 2013; Tanaka et al., 2012; Vogt et al., 2006) and GPNMB (Glycoprotein NMB, DC-associated transmembrane protein (DC-HIL), osteoactivin) (Chung et al., 2007; Ripoll et al., 2007). These markers represent novel targets to potentially reverse ercDC-mediated T cell inhibition. VSIG4 may be involved in the T-cell dysfunction associated with overexpressed p27kip, previously described by our group (Prinz et al., 2012). T cells in contact with VSIG4+ ercDCs preferentially expressed FOXO1, a positive regulator of the tolerance-inducing p27kip. NRP1 (Miyauchi et al., 2016) and CSF-1R are additional potential targets expressed by ercDCs where therapeutic reagents are currently in phase I studies (Link: https://clinicaltrials.gov/ct2/show/NCT02452424).

The results presented identify new potential mechanisms for immunoregulation in ccRCC malignancy involving direct communication between ercDCs and T cells. The next generation of studies will help determine whether the functional attributes suggested by the ercDC transcriptome are found in one single cell, or if they are represented by the assembly of multiple cell types with different functions. Single cell transcriptomics and multiplex mass cytometry may help clarify this issue. Our analysis using polychromatic flow cytometry indicates that the cells co-expressing the ercDC index markers, CD209 and CD14, largely also co-express the M1-marker CD64 and two M2-markers, CD204 and MerTK. This provides a first indication that the ercDCs may be a unique cell type on the single cell level with mixed macrophage polarization. Recently, macrophages from RCC tissues were analyzed by highly multiplex mass cytometry (Chevrier et al., 2017). Unfortunately, the analysis did not include the CD209 marker nor other relevant ercDC signature genes identified here, thus precluding a comparative analysis of our ercDCs to 17 TAM phenotypes identified in the mass cytometry study. Mass cytometry is a powerful approach, but as a front-line analysis tool it is disadvantaged by its bias towards previously characterized markers. By contrast, the transcriptomic and bioinformatics-based analysis described here allowed the identification of a novel series of proteins, including VSIG4 and GPNMB, or NRP1 and CSF1R, that likely contribute to T-cell suppression and tumor immune escape in ccRCC. In addition to providing new markers for mass cytometry studies, these proteins also represent potential targets for immunotherapy, and combined with checkpoint blockade, or other therapeutic strategies, may improve treatment outcome, especially for tumors with a high ercDC score.

## Experimental procedures

### Tissues, cells, cell culture

Tissue and blood collection were approved by the local ethics commission of the LMU München, and patients/donors consented to the donation. Tissue samples of histologically diagnosed clear cell renal cell carcinoma (ccRCC) (n=15) were obtained from untreated patients who underwent surgery at the Urologische Klinik Dr. Castringius Planegg (Munich, Germany). Patient characteristics are shown in table S1. Fresh postoperative material was used to prepare cell suspensions and cryosections (Figel et al., 2011). Peripheral blood mononuclear cells (PBMCs) from healthy donors (HD) were used to isolate monocytes (using CD14+ microbeads, Miltenyi Biotec) and to sort CD1c+ DC and slanDCs using FACS (Aria IIIu, BD). Before cell sorting, PBMCs were depleted of B and NK cells via CD19+ and CD56+ microbeads according to the manufacturer’s protocol (Miltenyi Biotec). M1- and M2- macrophages were generated from blood monocytes *in vitro*, as described (Martinez et al., 2006). Briefly, monocytes were cultured in 6-well plates (Nunc) in serum-free medium (5 x 106/4 ml of AIM-V) supplemented with M-CSF (50 ng/ml; R&D Systems) over 7 days followed by 18 h treatment with IFN-γ (20 ng/ml; R&D Systems) plus LPS (100 ng/ml; Sigma-Aldrich) for M1-polarization, or IL-4 (20 ng/ml; PromoKine) for M2-polarization.

### Cell sorting

ErcDCs and macrophages were sorted from ccRCC tissue-cell-suspensions stained with CD45-PeCy7, CD11c-APC, CD3-PB, CD209-PE (all BD Biosciences), CD14-PerCPCy5.5 (eBioscience) and LIVE/DEAD® Fixable Near-IR Dead Cell Stain Kit (Thermo Fisher Scientific). Sorting gates were set on CD209+CD14+ cells (ercDCs) and CD209-CD14+ cells (macrophages), among pre-gated CD45+, live, single CD11c+ CD3-cells. CD1c+ DC and slanDCs were sorted from B- and NK-depleted PBMCs of healthy donors (HD) using anti-CD11c-PE, anti-CD3-PB (all BD Biosciences), anti-CD56-APC (Beckman Coulter), anti-CD19-PB (Dako), anti-CD1c-PeCy7 (Biolegend), anti-slan-FITC (Miltenyi Biotec) and LIVE/DEAD® Fixable Near-IR Dead Cell Stain Kit. Gating strategy and instrument parameters are in supplemental Figure S2. Gates were set very strictly, not covering the whole population, to avoid contamination with other cell populations. Cell population purity varied between 98%-100%. Cells were directly sorted into 250 µl of RLT lysis buffer with ß-mercaptoethanol (RNeasy Micro Kit, Qiagen) using FACSAria IIIu (BD Biosciences), then homogenized (QIAshredder, Qiagen) and stored at -80°C. Details about sorted cell types, number of biological replicates are listed in table S2).

### Polychromatic flow cytometry

1-5 x 105 cells were incubated with antibodies for 30 min/4°C and LIVE/DEAD® Fixable Near-IR or Blue fluorescent Dead Cell Stain Kit, washed, optionally incubated with secondary antibodies and acquired at LSRII (BD Biosciences). In tissue-cell-suspensions, CD45 was used to identify leukocytes after exclusion of dead cells and doublets. Myeloid cells were selected based on CD11c (pan-myeloid marker in human) with exclusion of CD3+ T cells. Within CD11c+ cells, ercDCs and macrophages were distinguished as CD209/CD14 double-positive cells (ercDCs) and CD14 single-positive cells (macrophages). For M1/M2-macrophages, live cells where selected and doublets excluded before marker analysis. Antibodies are listed in table S13.

### Immunofluorescence histology and confocal microscopy

Cryosections were fixed with 4% paraformaldehyde (PFA, Merck) and stained with primary and fluorescent-labeled secondary antibodies as described (Figel et al., 2011). human CD209/DC-SIGN (mouse IgG2a, SantaCruz Biotechnology) followed by secondary antibodies donkey anti-goat-A488 (ThermoScientific) and rat anti-mouse-IgG-Cy5 or donkey anti-mouse-IgG-A647 (both Jackson ImmuneResearch); goat-anti-human VSIG4, rabbit-anti-human FOXO1 (Cell Signaling) and mouse-anti-human CD3 (UCHT1, BD) followed by secondary antibodies donkey anti-goat-A488, donkey anti-rabbit-RRX (Jackson ImmuneResearch) and donkey anti-mouse-A647 or rat anti-mouse IgG-Cy5. Slides were mounted with ProLong® Gold Antifade (ThermoFischer Scientific). Fluorescence images were captured with a laser scanning microscope TCS SP5 Leica Microsystems, Wetzlar, Germany) with settings as described (Figel et al., 2011).

### RNA isolation, microarray hybridization

RNA was prepared using RNeasy micro kit (Qiagen). RNA quantity and quality was assessed using a Nanodrop 1000 Spectrometer (Peqlab) and Agilent 2100 Bioanalyzer. From *in vitro*-generated cells (M1-, M2-macrophages) and CD14+ monocytes, three replica pools (each consisting of RNA from 5 different HD) were generated. RNA from flow-sorted cells was not pooled (table S2). RNA (30 ng of each replica pool; 0.5–15 ng of sorted cells) was subsequently amplified and converted into cDNA by a linear amplification method using WT-Ovation PicoSL System in combination with the Encore® Biotin Module (both from NuGen, San Carlos, US). cDNA was hybridized to Affymetrix GeneChip® Human Gene 1.0 ST Arrays.

### Microarray data preprocessing and probe set filtering

Raw intensity data were processed with R/Bioconductor (Bioconductor.org). If not stated otherwise, functions were called with default parameters. We calculated normalized expression values for each study group (in-house generated and external data sets, see table S2 and supplemental experimental procedure) independently using Robust Multichip Average (RMA, library ’oligo’) preprocessing including background correction and quantile normalization. Technical control probe sets as well as probe sets whose values did not vary between arrays (variance = 0) were excluded from all further analyses. Many-probe-sets-to-one-gene relationships were resolved by keeping only one probe set with highest variance for each gene. Further analyses included only informative genes, which we defined as group with the 50% most variable expression within the respective study group.

### Combining microarray studies

For comparing transcript levels across studies, gene expression values were merged based on annotated gene entrez ids and study batch effects were removed using the COMBAT method (Johnson et al., 2007) (library “inSilicoMerging”). We included all samples of a study in the merge and selected samples of interest afterwards.

### Hierarchical clustering and heatmaps

Gene and samplewise hierarchical clustering of expression profiles used euclidean distances and complete agglomeration method. Heatmaps represent color-coded genewise standardized expression levels (mean = 0, standard deviation = 1; z-score).

### Marker genes

Nearest shrunken centroid classifiers (Tibshirani et al., 2002) were constructed with the function “pamr.train” and cross-validated with the function “pamr.cv” (library “pamr”) (Figure S2C) on **i**nformative genes of compared data sets. The classification threshold (1.88) was set such that the false positive classification rate was smaller than 20% and a preferably small number of genes was obtained.

GeneMANIA network analysis (Warde-Farley et al., 2010) was conducted based on the ercDC_ccRCC marker genes with default parameters (data as of May 2014). Differentially expressed genes (DEGs) between ercDC_ccRCC&infMΦ_ascOvCa and control group (all other samples/groups listed in table S2) were identified by a linear model using empirical Bayes moderated t-tests (R package “limma”) and Benjamini-Hochberg correction for multiple testing. DEGs were defined by an adjusted p-value < 0.05.

For Gene ontology enrichment (GO) analysis, hypergeometric p-values for enrichment or depletion of differentially expressed genes in GO categories of the group “biological process” were calculated with the function “hyperGTest”, Bioconductor package “GOstats”. Informative genes served as background and p-value threshold was set to 0.001.

Enrichment of differentially expressed genes within InnateDB (Breuer et al., 2013, Lynn et al., 2010, Lynn et al., 2008) signaling pathway gene sets (from “KEGG“, “BioCarta“, “Reactome“, “NetPath“, “INOH“ und “PID“) was defined with default settings at InnateDB.com (data as of May 2014). Informative genes served as background. GSEA analysis was performed with GSEA 1.0 R-script of the Broad institute with default parameters (Subramanian et al., 2005) (http://www.broad.mit.edu/gsea/). For describing gene-expression-module-to-cell-type relationships, we first calculated median expression values per gene for each cell type. Eigengene values for gene expression modules as described (Xue et al., 2014) were calculated with the function “moduleEigengens” from the R-package “WGCNA” (Langfelder and Horvath, 2008). Eigengenes were correlated to cell types using Pearson’s method.

### ercDC score in the Cancer Genome Atlas (TCGA) and the validation (Rostock) cohort

Transcriptome profiling data (“HTSeq - FPKM-UQ”) of the TCGA ccRCC cohorts (KIRC, LAML) were downloaded from the Genomic Data Commons Portal (https://gdc-portal.nci.nih.gov/) on December 9, 2016 and December 15, 2016, respectively. Clinical data were obtained from the same platform on October 10, 2016 and November 3, 2016, respectively. To be used in this study, TCGA samples had to meet the following criteria: Patients with neoadjuvant therapies (“history_of_neoadjuvant_treatment”) were excluded. Moreover, only subjects with available survival data were considered (overall survival, OS, for the LAML cohort; cancer-specific survival (CSS) as defined in (Buttner et al., 2015) for the ccRCC cohort). Follow-up time was required to be greater than 0. Missclassified patients (Chen et al., 2016; Buttner et al., 2015) revealed by cluster analysis and/or by re-evaluation of tissue histology were also discarded from the TCGA ccRCC cohort. Characteristics of the final TCGA ccRCC cohort (n = 442 patients) are summarized in table S12.

The validation cohort included a collection of selected G1 and G3 ccRCC tissues (n = 14 each) and corresponding normal kidney tissue (n = 14) microarray data (“Rostock cohort”, Array GeneChip HG U133 Plus 2.0, Affymetrix) (Maruschke et al., 2011; Maruschke et al., 2014). Clinical data and recent follow up are found in table S12.

An ercDC score was established by gene expression deconvolution. Expression values of the 61 marker genes in the ercDC arrays were first collapsed by taking the median. FPKM-UQ expression values in the TCGA ccRCC cohort were log-transformed (log2(x+1)). Subsequently, using the marker gene set, a simple linear regression model was fit with “expression in ercDC cell type” as predictor and “expression in sample” as regressor. Finally, the slope of the linear model constituted the ercDC score of a sample. Conditional inference trees from R-package partykit_1.1-1 (Hothorn et al., 2006b, Hothorn and Zeileis, 2015)http://jmlr.org/papers/v16/hothorn15a.html were used to identify groups with significantly varying CSS curves in both the TCGA ccRCC cohort and the validation cohort. The p-value criterion of the conditional inference tree method was weakened (0.1) and the minimum group size was set to 10. Kaplan-Meier curves and corresponding log-rank tests using R-package survival_2.40-1 (Therneau and Grambsch, 2000) were applied for survival analysis. Further, R-package coin_1.1-3 (Hothorn et al., 2006a) was used to perform a linear trend test between ercDC score and tumor grade.

### Public access to raw data of data sets analyzed in this paper

Our data sets of human ercDCs and macrophages from ccRCC tissue, myeloid cells from PBMCs and *in vitro*-generated M1- and M2-macrophages are accessible via super series GSE108312.

## Supporting information

Brech et al._supplementary text tables figures

Brech et al_supplementary Table S8

Brech et al_supplementary Table S9

## Author contributions

Conception and design: D.B., E.N., P.J.N.; acquistion of data: D.B., E.K.; Microarray hybridization: M.I., J.B.; Bioinformatics: T.S., F.B., E.S. S.W., M.S., D.B.; data interpretation and manuscript writing: D.B., E.N., P.J.N., T.S., F.B., E.S., S.W., M.S.; Study supervision: E.N.. All authors approved the manuscript.

## Acknowledgements

We acknowledge A. Buchner and M. Maruschke for providing the Rostock microarray information, M. Schmitz, R. Wehner, R. Oberneder for providing tissues/tissue suspensions. We thank J. Ellwart, J. Mysliwietz, A. Brandl, A. Bettenbrock, C. Jäckel and B. Mosetter for technical help. We thank The Cancer Genome Atlas initiative (http://cancergenome.nih.gov/), all tissue donors and investigators who contributed to the acquisition and analyses of the samples used in this study. This work was supported by Robert Bosch Foundation (Stuttgart, Germany), ICEPHA Graduate School Tuebingen-Stuttgart, Deutsche Krebshilfe, the Helmholtz Alliance ‘Aging and Metabolic Programming, AMPro’ and DGF SFB-TR36. The authors declare no conflicts of interests.

## Supplemental information

Supplemental information includes 3 figures and 13 tables and can be found with this article online at super series GSE108312.

