## Supplementary material for "A mosaic renal myeloid subtype with T-cell inhibitory and protumoral features is linked to immune escape and survival in clear cell renal cell cancer": Brech et al._supplementary text tables figures

**Supplemental figures and legends**

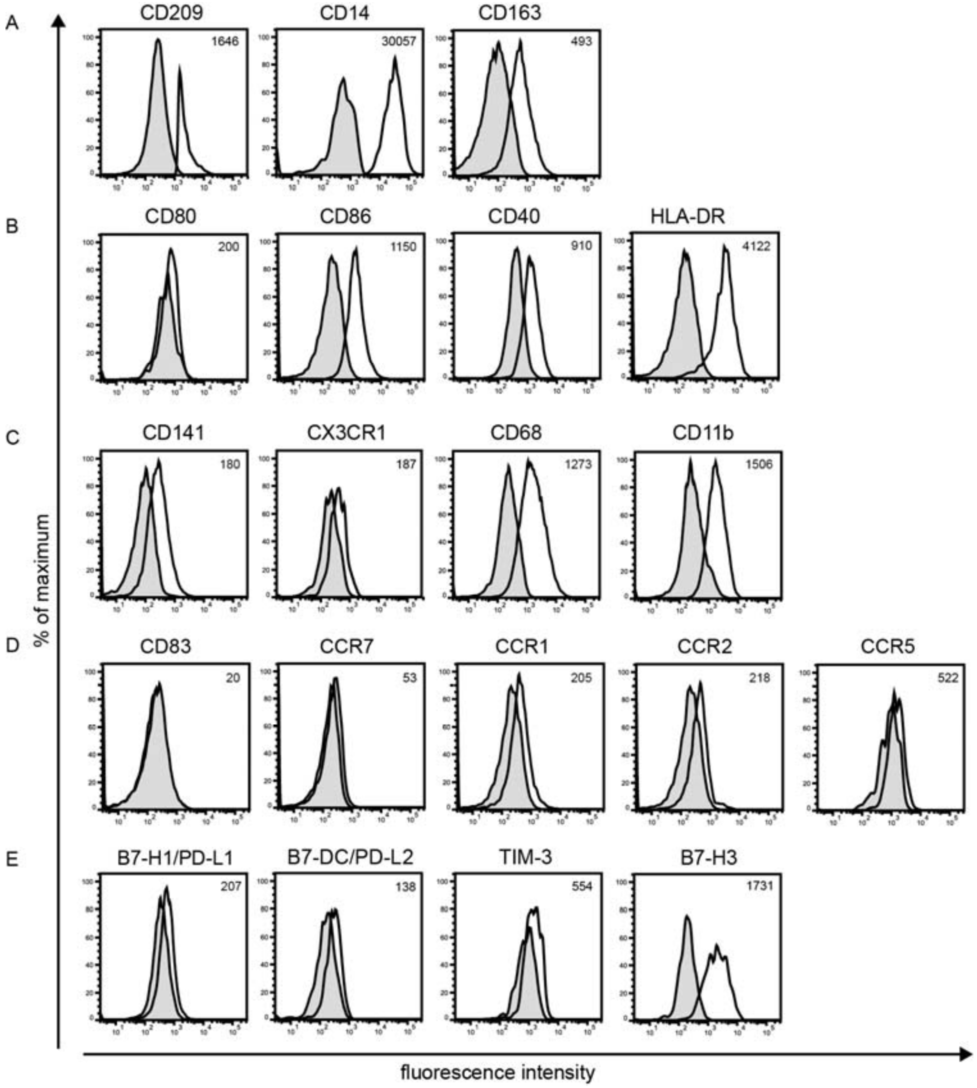


**Figure S1: Validation of selected DC- and macrophage markers as well as markers associated with immunoinhibition by flow cytometry.** Flow cytometric analysis of ccRCC tissue-cell-suspension to determine protein expression levels of (A) human DC (CD209) and macrophage (CD14, CD163) markers; (B) costimulatory molecules and HLA-DR, (C) markers associated with DC maturation state; (D) kidney-specific homing receptor $CX_3CR1$ and various myeloid markers; (E) markers associated with immunoinhibition. Exemplatory histograms of ercDCs ($CD209^+CD14^+$ cells among gated $CD45^+CD11c^+CD3$-live single cells) from ccRCC tissue cell suspensions (n=6) is shown. Grey filled histogram is the isotype staining or FMO (fluorescence-minus-one), black line is the specific antibody. Numbers indicate the median of control-corrected median fluorescence intensity (delta MFI).

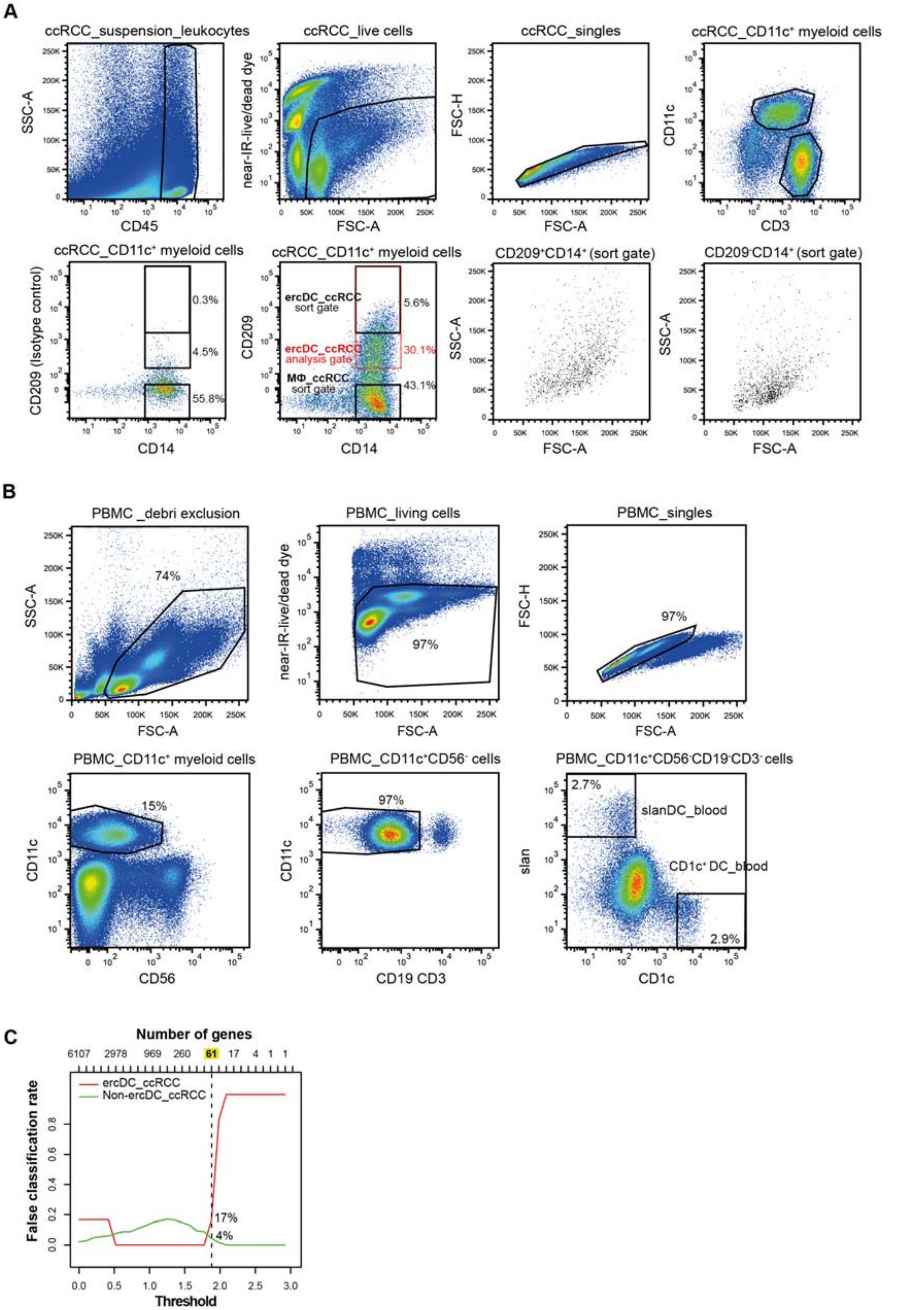


**Figure S2: Gating strategy for the sorting of ercDCs and macrophages from ccRCC tissue cell suspension, CD1c⁺ DCs and slanDCs from PBMCs (blood of healthy donors) and cross-validation of ercDC_ccRCC marker genes. Cross-validation related to figure 4A.** (A) ErcDCs and macrophages were sorted from ccRCC tissue-cell-suspensions stained with antibody combination CD45-PeCy7, CD11c-APC, CD3-PB, CD209-PE (all BD Biosciences), CD14-PerCPCy5.5 (eBioscience), LIVE/DEAD® Fixable Near-IR Dead Cell Stain Kit (Thermo

Fisher Scientific). (B) Sorting strategy for CD1c$^+$ DC and slanDCs from PBMCs of blood of healthy human donors. CD19/CD56-depleted PBMCs were stained with antibody combination anti-CD11c-PE, anti-CD3-PB (all BD Biosciences), anti-CD56-APC (Beckman Coulter), anti-CD19-PB (Dako), anti-CD1c-PeCy7 (Biolegend), anti-slan-FITC (Miltenyi Biotec) and LIVE/DEAD® Fixable Near-IR Dead Cell Stain Kit (Thermo Fisher Scientific). CD11c+ myeloid cells were selected after gating on CD45$^+$ leukocytes, live and single cells. CD1c$^+$ DCs and slanDCs were sorted from gated CD11c$^+$CD56$^-$CD19$^-$CD3$^-$ cells on the basis of the marker CD1c and slan, respectively. Positivity of both markers was identified with isotype control (not shown). Cell types were selected as following: CD45$^+$ leukocytes were selected in dot plots against SSC, dead cells and doublets were excluded and myeloid cells were selected by CD11c and exclusion of CD3. ErcDCs were selected as CD209 and CD14 co-expressing cells among gated live CD11c$^+$ CD3$^-$ cells. Macrophages within the tissue cell suspension were selected as CD209$^-$CD14$^+$ cells among gated CD45+ live single CD11c$^+$ CD3$^-$ cells. ErcDCs had a larger size (FSC-A) and granularity (SSC-A) than corresponding macrophages. CD209 positivity was judged with the help of an isotype control, CD14 positivity through CD14$^+$ monocytes within PBMC (not shown). Sorts used BD Aria IIIu flow cytometer with 100 µm nozzle and instrument settings modi “purity” or “single cell”. (C) Cross-validated misclassification error curves of the nearest shrunken centroid classifier. Depicted are the cross-validation errors for the training set at 30 different threshold levels of shrinkage. The training set comprised 6 ercDC_ccRCC and 182 non-ercDC_ccRCC samples (green line, contains all cell types used in Figure 4A). The number of discriminative genes as a function of threshold level in the classifier is indicated on top of the panel. The threshold for defining the set of 61 ercDC_ccRCC marker genes (dashed line) classifier was set in a way that a preferably small group of genes was choosen and the false negative classification rate was below 20 % while the false positive one was below 5%.

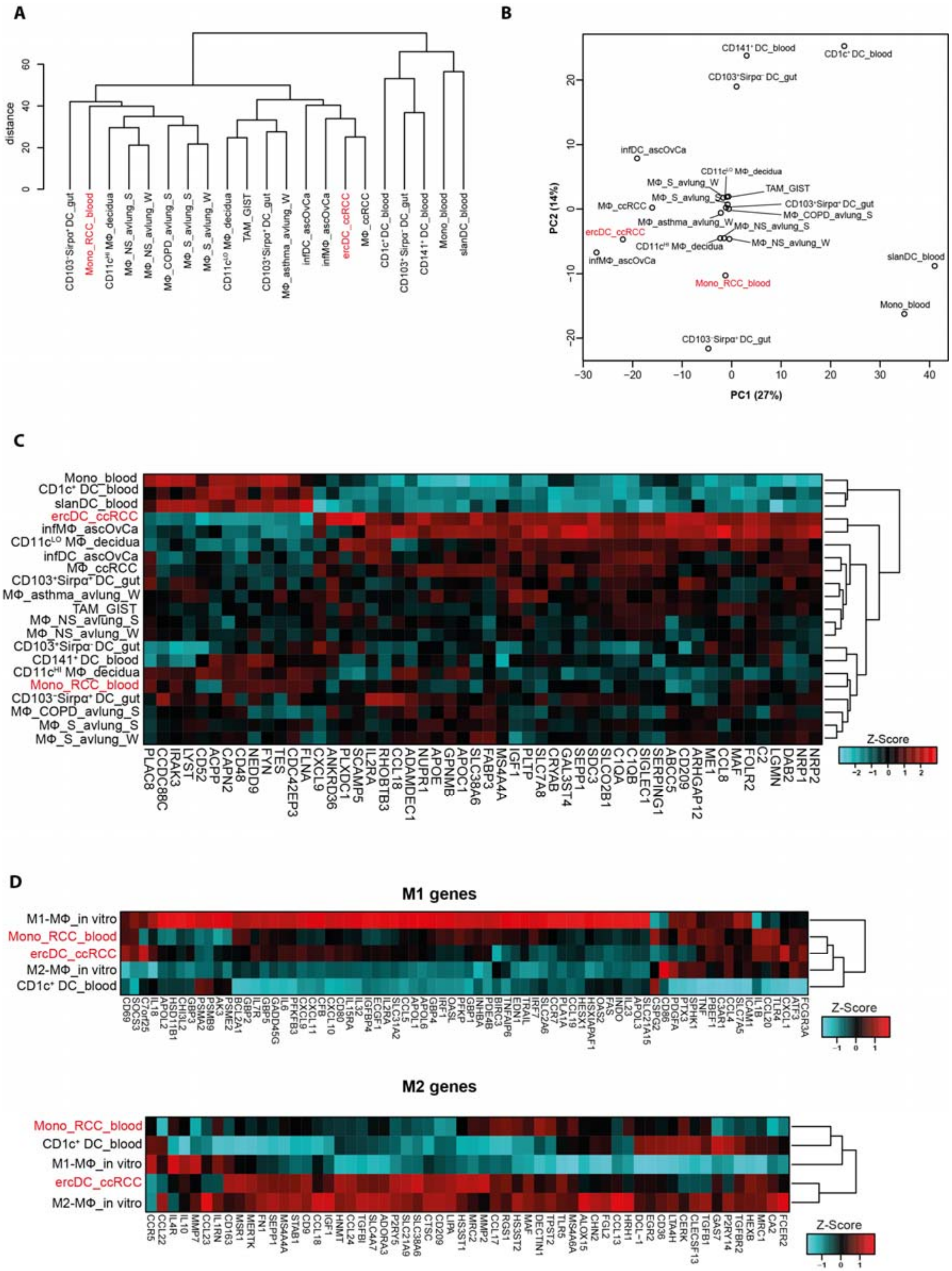


**Figure S3: Transcriptome comparison of monocytes from blood of RCC patients with ercDCs from ccRCC tissues and myeloid cells from non-lymphoid tissues after depletion of blood-specific genes. Related to figure 6.** Figures are calculated based on expression levels with blood-specific genes excluded. The list of blood-specific genes was from the tissue preferential expressed gene list (blood genes) (Mele et al., 2015). (A) Hierachical clustering of indicated cell types based on median expression values of replicate samples considering only "informative" genes (B) PCA on data set shown in (A) depicting the first two PCs that describe 27% and 14% of the variance. (C) Heatmap comparing ercDC_ccRCC marker genes (generated with NSCM (Tibshirani et al., 2002)) with blood monocytes of RCC patients (Mono_RCC_blood) (Chittezhath et al., 2014). Relative expression levels (Z-score) of the 61 marker genes are shown. (D) Heatmap showing relative expression levels of genes associated with M1- and M2-macrophages in Mono_RCC_blood and ercDC_ccRCC.

**Supplemental tables**

**Table S1: ccRCC tissues.** Samples were obtained anonymously with pathologic tumor description but without specification of personal data.

| **Patient-ID** | **TNM classification[a]** | **Gender** |
|---|---|---|
| DD1* | n.a. | n.a. |
| DD2* | n.a. | n.a. |
| RCC42 | pT3b pN0 G3 | Female |
| RCC66* | pT3a pN0 M1 G3 | Male |
| RCC69 | pT3a pN0 (0/4) L0 V0 G2 R0 | Female |
| RCC71 | pT3a pN0 G2 | Female |
| RCC74* | pT1a pN0 G1 | Male |
| RCC89* | pT3a pNX G2 | Female |
| RCC90 | pT1a pNX G2 | Male |
| RCC91* | pT3a pNX G3 | Male |
| RCC94* | pT1b pN0 G2 | Female |
| RCC97* | pT2b pN0 G2 | Female |
| RCC105 | pT3a pNX G3 | Male |
| RCC114 | pT3c pN0 G2 | Male |
| RCC118 | at least pT3b pN0 G3 | Male |

[a] Classification according to Union International Contre le Cancer (UICC), T: primary tumor size, N: lymph node involvement, M: distant metastasis; * ccRCC tissues used for gene expression analysis

**Table S2A: Myeloid cell types used for Affymetrix arrays (in house generated data sets).**

| Cell type | Origin | Name | Marker profile | Replicates | RNA RIN | (Sorted) total cell number |
|---|---|---|---|---|---|---|
| **ercDC** | ccRCC tissue | ercDC_ccRCC | CD11c$^+$ CD209$^+$ CD14$^+$ | RCC66 | 8.6 | $2.1 \times 10^4$ |
| | | | | RCC74 | 6.1 | $1.7 \times 10^4$ |
| | | | | RCC89 | 8.0 | $9.5 \times 10^4$ |
| | | | | RCC91 | 8.0 | $1.1 \times 10^4$ |
| | | | | RCC94 | 6.7 | $1.1 \times 10^4$ |
| | | | | DD2 | 5.6 | $0.7 \times 10^4$ |
| **MΦ** | ccRCC tissue | MΦ_ccRCC | CD11c$^+$ CD209$^-$ CD14$^+$ | RCC66 | 8.8 | $6.0 \times 10^4$ |
| | | | | RCC89 | 8.0 | $2.2 \times 10^5$ |
| | | | | RCC91 | 7.3 | $4.2 \times 10^4$ |
| | | | | RCC94 | 6.1 | $2.5 \times 10^4$ |
| | | | | DD1 | 7.8 | $0.5 \times 10^4$ |
| | | | | DD2 | 7.8 | $6.1 \times 10^4$ |
| **CD1c$^+$ DC** | Blood healthy | CD1c$^+$ DC_blood | CD11c$^+$ CD1c$^+$CD19$^-$ | Donor 1 | 9.0 | $2.0 \times 10^4$ |
| | | | | Donor 2 | 7.4 | $1.2 \times 10^4$ |
| | | | | Donor 3 | 6.2 | $1.0 \times 10^4$ |
| **slanDC** | Blood healthy | slanDC_blood | CD11c$^+$slan$^+$ CD19$^-$ | Donor 1 | 7.6 | $1.5 \times 10^4$ |
| | | | | Donor 2 | 8.8 | $5.0 \times 10^4$ |
| | | | | Donor 3 | 8.6 | $2.7 \times 10^4$ |
| **Monocytes**[a] | Blood healthy | Mono_blood | CD14$^+$ | Donor Pool 1[+] | 8.0 | $5.0 \times 10^5$ |
| | | | | Donor Pool 2[+] | 8.6 | $5.0 \times 10^5$ |
| | | | | Donor Pool 3[+] | 7.9 | $5.0 \times 10^5$ |
| **M1-MΦ**[b] | In vitro healthy | M1-MΦ_in vitro | CD14$^+$ | Donor Pool 1[+] | 8.2 | $5.0 \times 10^5$ |
| | | | | Donor Pool 2[+] | 8.5 | $5.0 \times 10^5$ |
| | | | | Donor Pool 3[+] | 8.7 | $5.0 \times 10^5$ |
| **M2-MΦ**[#] | In vitro healthy | M2-MΦ_in vitro | CD14$^+$ | Donor Pool 1[+] | 8.7 | $5.0 \times 10^5$ |
| | | | | Donor Pool 2[+] | 8.8 | $5.0 \times 10^5$ |
| | | | | Donor Pool 3[+] | 8.8 | $5.0 \times 10^5$ |

[a] Monocytes isolated from PBMC using CD14$^+$ microbeads (Miltenyi). [b] M1- and M2-macrophages (MΦ) generated as described (Martinez et al., 2006). [+] Each pool consists of cells from 5 different donors.

**Table S2B: human myeloid cell subtypes of external data sets used for comparison to ercDC_ccRCC.**

| Name | Origin | No. of replicates | Accession code |
|---|---|---|---|
| CD141$^+$ DC_blood | Blood healthy | 3 | E-TABM-34 (Lindstedt et al., 2005) |
| CD11c$^{HI}$ MΦ_decidua | decidua first trimester | 8 | GSE22342 (Houser et al., 2011) |
| CD11c$^{LO}$ MΦ_decidua | | 8 | |
| CD103$^+$Sirpα$^-$ DC_gut | gut | 3 | GSE50380 (Watchmaker et al., 2014) |
| CD103$^+$Sirpα$^+$ DC_gut | | 5 | |
| CD103$^-$Sirpα$^+$ DC_gut | | 3 | |
| MΦ_NS_avlung_S | avlung | 24 | GSE13896 (Shaykhiev et al., 2009) (S) |
| MΦ_S_avlung_S | | 34 | |
| MΦ_COPD_avlung_S | | 12 | |
| MΦ_NS_avlung_W | avlung | 15 | GSE2125 (Woodruff et al., 2005) (W) |
| MΦ_S_avlung_W | | 15 | |
| MΦ_asthma_avlung_W | | 15 | |
| TAM_GIST | GIST | 12 | GSE51697 (Cavnar et al., 2013) |
| infDC_ascOvCa | ascOvCa | 5 | GSE40484 (Segura et al., 2013) |
| Mono_RCC_blood | Blood of RCC | 4 | GSE38424 (Chittezhath et al., 2014) |

_S_ = smoker, _NS_ = non-smoker, _S = Shaykhiev data; _W = Woodruff data; GIST = gastrointestinal stromal tumor, COPD = chronic obstructive pulmonary disease, av = alveolar; ascOvCa = ascites of ovarian cancer

**Table S3: M1-associated genes (Houser et al., 2011, Martinez et al., 2006, Shaykhiev et al., 2009).** Bold: ercDC_ccRCC marker gene, purple: ercDC_ccRCC DEGs

| Gene name | Gene symbol/alternative name | Informative |
|---|---|---|
| **Membrane receptors** | | |
| Complement component 3a receptor 1 | C3AR1 | yes |
| CCR7 | CCR7 | yes |
| CD69 | CD69 | yes |
| CD80 | CD80 | yes |
| CD86 | CD86 | no |
| Fc fragment of IgG, high affinity Ia, receptor (CD64) | CD64/FCGR1A | yes |
| Fc fragment of IgG, high affinity IIa, receptor (CD32) | CD32/FCGR2A | yes |
| Fc fragment of IgG, high affinity IIIa, receptor (CD16) | CD16/FCGR3A | yes |
| ICAM1 | ICAM1 | yes |
| Interleukin 15 receptor α chain | IL15RA | yes |
| Interleukin 2 receptor α chain | **IL2RA** | yes |
| Interleukin 7 receptor | IL7R | yes |
| Toll-like receptor 2 | TLR2 | yes |
| Toll-like receptor 4 | TLR4 | yes |
| **Cytokines and chemokines** | | |
| CXCL11 | CXCL11 | yes |
| CCL14 | CCL14 | no |
| CCL15 | CCL15 | no |
| CCL19 | CCL19 | yes |
| CCL20 | CCL20 | yes |
| CCL4 | CCL4 | yes |
| CCL5 | CCL5 | yes |
| CXCL1 | CXCL1 | yes |
| CXCL10 | CXCL10 | yes |
| CXCL9 | **CXCL9** | yes |
| Thymidine phosphorylase | TYMP | yes |
| Interleukin 12B | IL12B | no |
| Interleukin 15 | IL15 | no |
| Interleukin 18 | IL18 | yes |
| Interleukin 1B | IL1B | yes |
| Interleukin 23 | IL23 | yes |
| Interleukin 32 | IL32 | yes |
| Interleukin 6 | IL6 | yes |
| Nicotinamide phosphoribosyltransferase | NAMPT/PBEF1 | yes |
| Tumor necrosis factor ligand superfamily, member 2 | TNF | yes |
| Tumor necrosis factor, α-induced protein 6 | TNFAIP6 | yes |
| Tumor necrosis factor (ligand) superfamily, member 10 | TRAIL | yes |
| **Apoptosis associated genes** | | |
| BCL2-related protein A1 | BCL2A1 | yes |
| Baculoviral 1AP repeat-containing 3 | BIRC3 | yes |
| Tumor necrosis factor receptor superfamily, member 6 | FAS | yes |
| Growth arrest and DNA-damage-inducible, γ | GADD45G | yes |
| XIAP associated factor-1 | HSXIAPAF1 | yes |
| **Transport proteins** | | |
| Solute carrier family 21, member 15 | SLC21A15 | yes |
| Solute carrier family 2, member 6 | SLC2A6 | yes |
| Solute carrier family 31, member 2 | SLC31A2 | yes |
| Solute carrier family 7, member 5 | SLC7A5 | yes |
| **Enzymes and other proteins** | | |
| Adenylate kinase 3 | AK3 | yes |
| B factor, properdin (complement factor B) | CFB | yes |
| Chitinase 3-like 2 | CHI3L2 | yes |
| Hydroxysteroid (11-β) dehydrogenase 1 | HSD11B1 | yes |
| Indoleamine 2,3 dioxygenase 1 | IDO1 | yes |
| Nitric oxide synthase 2A (inducible) | NOS2A | no |
| 2‘-5‘-oligoadenylate synthetase 2 | OAS2 | yes |
| 2’-5’-oligoadenylate synthetase-like | OASL | yes |
| Phosphodiesterase 4B, cAMP-specific | PDE4B | yes |

| | | |
|---|---|---|
| 6-phosphofructo-2-kinase/fructo-2,6-biphosphatase 3 | PFKFB3 | yes |
| Phosphofructokinase | PFKP | yes |
| Phospholipase A1 member A | PLA1A | yes |
| Proteasome subunit α type 2 | PSMA2 | yes |
| Proteasome subunit β type 9 | PSMB9 | yes |
| Proteasome activator subunit 2 | PSME2 | yes |
| Sphingosine kinase 1 | SPHK1 | yes |
| **Extracellular mediators** | | |
| Apolipoprotein L1 | APOL1 | yes |
| Apolipoprotein L2 | APOL2 | yes |
| Apolipoprotein L3 | APOL3 | yes |
| Apolipoprotein L6 | APOL6 | yes |
| Chondroitin sulfate proteoglycan 2 | CSPG2 | yes |
| Endothelin 1 | EDN1 | yes |
| Insulin-like growth factor binding protein 4 | IGFBP4 | yes |
| Inhibin β A | INHBA | yes |
| Platelet-derived growth factor α | PDGFA | yes |
| Pentraxin 3 | PTX3 | yes |
| **DNA binding factors** | | |
| Activating transcription factor 3 | ATF3 | yes |
| Homeobox expressed in ES cells 1 | HESX1 | yes |
| Interferon regulatory factor 1 | IRF1 | yes |
| Interferon regulatory factor 7 | IRF7 | yes |
| Storkhead box 1 | STOX1 | no |
| **Signaling-associated genes** | | |
| Guanylate-binding protein 1, IFN-inducible | GBP1 | yes |
| Guanylate-binding protein 2, IFN-inducible | GBP2 | yes |
| Guanylate-binding protein 3 | GBP3 | yes |
| Guanylate-binding protein 4 | GBP4 | yes |
| Guanylate-binding protein 5 | GBP5 | yes |
| Immunoresponsive 1 homolog | IRG1 | no |
| Suppressor of cytokine signaling 3 | SOCS3 | yes |
| **Function not known** | | |
| Chromosome 7 open reading frame 25 | C7orf25 | yes |

**Table S4: M2-associated genes (Houser et al., 2011, Martinez et al., 2006, Shaykhiev et al., 2009).** Bold: ercDC_ccRCC marker gene, purple: ercDC_ccRCC signature genes

| Gene name | Gene symbol/alternative name | Informative |
|---|---|---|
| **Membrane receptors** | | |
| Adenosine A3 receptor | ADORA3 | yes |
| CCR5 | CCR5 | yes |
| CD163 | CD163 | yes |
| CD36 | CD36 | yes |
| CD9 | CD9 | yes |
| C-type lectin family 7A | CLEC7A/DECTIN1 | yes |
| C-type lectin domain family 10, member A | CLEC10A/CLECSF13 | yes |
| CXCR4 | CXCR4 | yes |
| CD302 | CD302/DCL-1 | yes |
| CD209 | **CD209/DC-SIGN** | yes |
| Fc fragment IgE, low affinity II, receptor for (CD23) | FCER2/CD23 | yes |
| G protein-coupled receptor 86 | GPR86 | yes |
| Histamine receptor H1 | HRH1 | yes |
| IL4R | IL4R | yes |
| c-mer protooncogene tyrosine kinase | MERTK | yes |
| Mannose receptor, C type 1 | CD206/MRC1 | yes |
| Mannose receptor, C type 2 | MRC2 | yes |
| Membrane-spanning 4-domains, subfamily A, member 4A | **MS4A4A** | yes |
| Membrane-spanning 4-domains, subfamily A, member 6A | MS4A6A | yes |
| Macrophage scavenger receptor 1 | CD204/MSR1 | yes |
| Purinergic receptor P2Y, G-protein coupled, 14 | P2RY14 | yes |
| Purinergic receptor P2Y, G protein-coupled, 5 | P2RY5/LPAR6 | yes |
| Stabilin 1 | STAB1 | yes |
| Transforming growth factor β receptor II | TGFBR2 | yes |
| Toll-like receptor 5 | TLR5 | yes |
| **Cytokines and chemokines** | | |
| CCL13 | CCL13 | yes |
| CCL17 | CCL17 | yes |
| CCL18 | **CCL18** | yes |
| CCL22 | CCL22 | yes |
| CCL23 | CCL23 | yes |
| CCL24 | CCL24 | yes |
| Insulin-like growth factor 1 | **IGF1** | yes |
| Interleukin 10 | IL10 | yes |
| IL1 receptor antagonist | IL1RN | yes |
| TGFB1 | TGFB1 | yes |
| **Transport proteins** | | |
| Solute carrier family 21, member 15 | **SLCO2B1/SLC21A9** | yes |
| Solute carrier family 38, member 6 | **SLC38A6** | yes |
| Solute carrier family 4, member 7 | SLC4A7 | yes |
| **Enzymes and other proteins** | | |
| Adenosine kinase | ADK | yes |
| Arachidonate 15-lipoxygenase | ALOX15 | yes |
| Arginase 1 | ARG1 | no |
| Carbonic anhydrase II | CA2 | yes |
| Ceramide kinase | CERK | yes |
| Collagen, type VI, α 2 | COL6A2 | no |
| Cathepsin C | CTSC | yes |
| Hexosaminidase B | HEXB | yes |
| Histamine *N*-methyltransferase | HNMT | yes |
| Heparan sulfate (glucosamine) 3-*O*-sulfotransferase 1 | HS3ST1 | yes |
| Heparan sulfate (glucosamine) 3-*O*-sulfotransferase 2 | HS3ST2 | yes |
| Lipase A cholesterol esterase | LIPA | yes |
| Leukotriene A4 hydrolase | LTA4H | yes |
| Matrixmetallopeptidase 2 (gelatinase A, 72-kDa gelatinase) | MMP2 | yes |
| MMP7 (matrilysin, uterine) | MMP7 | yes |

| | | |
|---|---|---|
| MMP9 (gelatinase B, 92-kDa gelatinase) | MMP9 | yes |
| Tyrosylprotein sulfotransferase 2 | TPST2 | yes |
| **Extracellular mediators** | | |
| Chimerin 2 | CHN2 | yes |
| Fibrinogen-like 2 | FGL2 | yes |
| Fibrionectin 1 | FN1 | yes |
| Selenoprotein P, plasma, 1 | **SEPP1** | yes |
| Transforming growth factor, beta-induced, 68 kDa | TGFBI | yes |
| **DNA binding factors** | | |
| Early growth response 2 | EGR2 | yes |
| Growth arrest-specific 7 | GAS7 | yes |
| *v-maf* musculoaponeurotic fibrosarcoma oncogene homolog | **MAF** | yes |
| **Signaling associated genes** | | |
| Regulator of G protein signaling 1 | RGS1 | yes |

**Table S5: Markers of 3 tissue macrophage subtypes.** Adopted from (Mosser and Edwards, 2008), supplemented with genes from (Gustafsson et al., 2008) (green).

| **Marker** | **Gene symbol/ alternative name** |
|---|---|
| **Classical activated, bactericidal macrophages** | |
| CCL15 | CCL15[a] |
| CCL20 | CCL20[b] |
| CXCL10 | CXCL10 |
| CXCL11 | CXCL11 |
| CXCL9 | CXCL9 |
| IL-12 | IL12A/B[b] |
| iNOS | NOS2[b] |
| **Wound healing associated, tissue modulating macrophages** | |
| CCL17 | CCL17 |
| CCL18 | CCL18 |
| CCL22 | CCL22 |
| DCIR | DCIR/CLEC4A |
| Factor XIII-A | F13A1 |
| IGF-1 | IGF1 |
| IL-27Rα | IL27RA |
| RELMα | FIZZ1 |
| Stabilin 1 | STAB1 |
| YM1 | YM1 |
| BEX3 | NGFRAP1 |
| C1q, subunit A | C1QA |
| C1q, subunit B | C1QB |
| COL1A2 | COL1A2 |
| COL3A1 | COL3A1 |
| COL4A2 | COL4A2 |
| COL6A3 | COL6A3 |
| FN1 | FN1 |
| Gas6 | GAS6 |
| HIF-2α | EPAS1 |
| Hsp27 | HSPB1 |
| ITGB5 | ITGB5 |
| MMP-9 | MMP9 |
| PADI4 | PADI4 |
| Protein S | PROS1 |
| S100-A4 | S100A4 |
| SDC2 | SDC2 |
| Serpin F1 | SERPINF1 |
| VCAN | VCAN |
| **Immunoregulatory macrophages** | |
| CCL1 | CCL1 |
| IL-10 | IL10 |
| LIGHT | LIGHT/TNFSF14 |
| SPHK1 | SPHK1 |
| α2M | A2M |
| CD9 | CD9 |
| c-Maf | MAF |
| DAB2 | DAB2 |
| CD209 | CD209/DC-SIGN |
| DPEP2 | DPEP2 |
| IRAK3 | IRAK3 |
| CD206 | CD206/MRC1 |
| PGDS | PGDS |
| TNFRSF21 | TNFRSF21 |
| TREM2 | TREM2 |
| VSIG4 | VSIG4 |

[a] Gene not present in study group and could not be analyzed; [b] Gene not shown in **figure 2E** because it was only marginally expressed by all cell types or not suitable because it was not strongly expressed by positive controls compared to negative controls

**Table S6: Genes associated with invasion and angiogenesis** (Wang et al., 2012), gene set "angiogenesis" from GSEA MSigDB database (Subramanian et al., 2005) and factors listed at http://www.sabiosciences.com/rt_pcr_product/HTML/PAHS-024A.html. (as of May 2014)

| Gene | Entrez ID |
|---|---:|
| EGF | 1950 |
| IL18 | 3606 |
| IL8 | 3576 |
| TGFB2 | 7042 |
| VEGFA | 7422 |
| CCL2 | 6347 |
| CXCL2 | 2920 |
| CXCL3 | 2921 |
| CXCL5 | 6374 |
| FGF1 | 2246 |
| FGF2 | 2247 |
| HGF | 3082 |
| HIF1A | 3091 |
| IGF1 | 3479 |
| IL15 | 3600 |
| IL1B | 3553 |
| IL6 | 3569 |
| KDR | 3791 |
| MMP2 | 4313 |
| MMP9 | 4318 |
| NRP1 | 8829 |
| NRP2 | 8828 |
| PDGFA | 5154 |
| PGF | 5228 |
| PLAU | 5328 |
| PTGS1 | 5742 |
| TEK | 7010 |
| TGFA | 7039 |
| TGFB1 | 7040 |
| TGFBR1 | 7046 |
| TNF | 7124 |
| VEGFC | 7424 |
| CTSB | 1508 |
| ANGPT1 | 284 |
| FLT1 | 2321 |
| IL17A | 3605 |
| SPP1 | 6696 |
| GPNMB | 10457 |
| MT1G | 4495 |
| MMP7 | 4316 |
| CCL8 | 6355 |
| CCL22 | 6367 |
| CCL7 | 6354 |
| CCL4L2 | 388372 |
| APOE | 348 |
| APOC1 | 341 |
| TNFAIP6 | 7130 |
| IL1A | 3552 |
| A2M | 2 |
| NR1H3 | 10062 |
| CTSL1 | 1514 |
| MT1E | 4493 |
| PLTP | 5360 |

| | |
|---|---|
| CCL4L1 | 9560 |
| ADAMDEC1 | 27299 |
| DFNA5 | 1687 |
| MT1H | 4496 |
| SLC39A8 | 64116 |
| FPR3 | 2359 |
| CCL20 | 6364 |
| CXCL1 | 2919 |
| MT2A | 4502 |
| SLCO2B1 | 11309 |
| ACP5 | 54 |
| TM4SF1 | 4071 |
| TGM2 | 7052 |
| NDP | 4693 |
| PMP22 | 5376 |
| ABCA1 | 19 |
| TM7SF4 | 81501 |
| LPL | 4023 |
| INDO | 3620 |
| CCL3 | 6348 |
| GJB2 | 2706 |
| CYP27B1 | 1594 |
| SLC1A3 | 6507 |
| PTGES | 9536 |
| CCL3L1 | 6349 |
| MAPK13 | 5603 |
| CCR7 | 1236 |
| IL23A | 51561 |
| VSIG4 | 11326 |
| STEAP3 | 55240 |
| SCG5 | 6447 |
| CCND1 | 595 |
| CA12 | 771 |
| WBP5 | 51186 |
| LGMN | 5641 |
| BCAT1 | 586 |
| EMP1 | 2012 |
| MAP1LC3A | 84557 |
| FBP1 | 2203 |
| F3 | 2152 |
| RASGRP3 | 25780 |
| GSN | 2934 |
| CRABP2 | 1382 |
| ETV5 | 2119 |
| PPAP2B | 8613 |
| IRAK2 | 3656 |
| CD9 | 928 |
| SCD | 6319 |

**Table S7: List of 61 marker genes of ercDC_ccRCC.** Genes were computed with the nearest shrunken centroids method (NSCM) (Tibshirani et al., 2002). 39 genes were upregulated, 22 genes downregulated compared to the other cells listed in heatmap **figure 4A**. Depicted are the shrunken centroid values (positive for upregulated, negative for downregulated genes) that indicate the degree of contribution of the gene to the profile.

| Gene symbol | Entrez ID | Shrunken Centroid value |
|---|---|---|
| CD209 | 30835 | 0.292 |
| IGF1 | 3479 | 0.259 |
| PLXDC1 | 57125 | 0.200 |
| SEPP1 | 6414 | 0.194 |
| FOLR2 | 2350 | 0.185 |
| PLTP | 5360 | 0.168 |
| SCAMP5 | 192683 | 0.160 |
| NUPR1 | 26471 | 0.142 |
| C2 | 717 | 0.141 |
| ME1 | 4199 | 0.140 |
| GPNMB | 10457 | 0.127 |
| SDC3 | 9672 | 0.121 |
| IL2RA | 3559 | 0.100 |
| SLCO2B1 | 11309 | 0.099 |
| NRP1 | 8829 | 0.088 |
| CCL8 | 6355 | 0.086 |
| APOC1 | 341 | 0.083 |
| CCL18 | 6362 | 0.080 |
| GAL3ST4 | 79690 | 0.073 |
| SIGLEC1 | 6614 | 0.073 |
| SLC38A6 | 145389 | 0.071 |
| C1QA | 712 | 0.051 |
| CRYAB | 1410 | 0.050 |
| SERPING1 | 710 | 0.048 |
| ARHGAP12 | 94134 | 0.042 |
| APOE | 348 | 0.040 |
| ANKRD36 | 375248 | 0.038 |
| DAB2 | 1601 | 0.031 |
| RHOBTB3 | 22836 | 0.030 |
| ADAMDEC1 | 27299 | 0.026 |
| MS4A4A | 51338 | 0.024 |
| NRP2 | 8828 | 0.023 |
| ABCC5 | 10057 | 0.020 |
| C1QB | 713 | 0.018 |
| LGMN | 5641 | 0.016 |
| FABP3 | 2170 | 0.014 |
| CXCL9 | 4283 | 0.011 |
| SLC7A8 | 23428 | 0.008 |
| MAF | 4094 | 0.003 |
| TES | 26136 | -0.006 |
| LYST | 1130 | -0.013 |
| NEDD9 | 4739 | -0.018 |
| CD48 | 962 | -0.023 |
| FYN | 2534 | -0.025 |
| ACPP | 55 | -0.033 |
| FLNA | 2316 | -0.037 |
| IRAK3 | 11213 | -0.045 |
| CAPN2 | 824 | -0.056 |
| LILRA2 | 11027 | -0.058 |
| CYTIP | 9595 | -0.067 |
| EMR3 | 84658 | -0.071 |
| CDC42EP3 | 10602 | -0.078 |

| | | |
|---|---|---|
| LILRA1 | 11024 | -0.084 |
| PLAC8 | 51316 | -0.088 |
| FAM65B | 9750 | -0.123 |
| CCDC88C | 440193 | -0.154 |
| S100A12 | 6283 | -0.157 |
| CD52 | 1043 | -0.228 |
| FGR | 2268 | -0.245 |
| CFP | 5199 | -0.282 |
| FCN1 | 2219 | -0.461 |

**Table S8: Upregulated ercDC_ccRCC DEGs.** Please find list in separate excel file

**Table S9: Downregulated ercDC_ccRCC DEGs.** Please find list in separate excel file

**Table S10: Top 20 overrepresented signaling pathways of upregulated ercDC_ccRCC DEGs in InnateDB analysis** (Breuer et al., 2013, Lynn et al., 2010, Lynn et al., 2008) (signaling pathways in orange are significantly overexpressed, adjusted p-value <0.05) or **Top 20 overrepresented signaling pathways of ercDC_ccRCC&infMP_ascOvCa in GSEA analysis**

| **InnateDB upregulated signaling pathway s of ercDC_RCC DEGs** | **Adjusted p-value** |
|---|---|
| Lysosome | $3.99 \times 10^{-7}$ |
| Complement and coagulation cascades | $1.57 \times 10^{-4}$ |
| Beta1 integrin cell surface interactions | $7.68 \times 10^{-4}$ |
| Collagen biosynthesis and modifying enzymes | 0.009 |
| Beta3 integrin cell surface interactions | 0.016 |
| HDL-mediated lipid transport | 0.020 |
| Phagosome | 0.041 |
| ECM-receptor interaction | 0.041 |
| Regulation of complement cascade | 0.068 |
| NOD-like receptor signaling pathway | 0.079 |
| Staphylococcus aureus infection | 0.098 |
| Validated transcriptional targets of AP1 family members Fra1 and Fra2 | 0.113 |
| Integrin cell surface interactions | 0.114 |
| Inhibition of matrix metalloproteinases | 0.148 |
| Toll-like receptor signaling pathway | 0.150 |
| Chemokine receptors bind chemokines | 0.158 |
| Degradation of collagen | 0.165 |
| Fibrinolysis pathway | 0.166 |
| Metal ion SLC transporters | 0.166 |
| Platelet amyloid precursor protein pathway | 0.166 |

| **GSEA overrepresented signaling pathways of ercDC_ccRCC&infMP_ascOvCa vs crtl group** | **Data base** | **NES [a]** | **Marker genes/related genes „Leading edge“ [b]** |
|---|---|---|---|
| Lysosome | KEGG | 1.94 | LGMN |
| Complement and coagulation cascades | KEGG | 1.86 | C2, SERPING1, C1QA, C1QB |
| Lipid digestion mobilization and transport | Reactome | 1.85 | PLTP, APOE |
| Activation of chaperone genes by XBP1S | Reactome | 1.76 | |
| Arginine and proline metabolism | KEGG | 1.69 | |
| Iron uptake and transport | Reactome | 1.68 | HMOX1 |
| Complement cascade | Reactome | 1.66 | C2, C1QA, C1QB |
| Extracellular matrix organization | Reactome | 1.66 | |
| MHC class II antigen presentation | Reactome | 1.64 | LGMN |
| Unfolded protein response | Reactome | 1.63 | |
| Diabetes pathways | Reactome | 1.61 | PLA2G7 |
| Peptide ligand binding receptors | Reactome | 1.60 | CXCL9, C3AR1 |
| Collagen formation | Reactome | 1.60 | |
| Response to elevated platelet cytosolic_CA2 | Reactome | 1.58 | SERPING1 |
| Toll like receptor signaling pathway | KEGG | 1.55 | CXCL9, CD14 |
| Axon guidance | Reactome | 1.55 | NRP1, NRP2, GFRA2 |
| Amino sugar and nucleotide sugar metabolism | KEGG | 1.55 | |
| Signal transduction by L1 | Reactome | 1.54 | NRP1 |
| Systemic lupus erythematosus | KEGG | 1.53 | C2, C1QA, C1QB |
| ECM receptor interaction | KEGG | 1.53 | SDC3 |

[a] NES was used for the arrangement of the top 20 signaling pathways with the strongest expression differences between ercDC_ccRCC&infMΦ_ascOvCa and control group because p-values and FDR values were very high. [b] ercDC_ccRCC marker genes and their related genes that belong to the so called “leading edge” genes of respective gene sets.

**Table S11: Modules (Xue et al., 2014) and their assigned correlating/associated stimuli. Related to figure 7.** We assigned a correlation/association of a stimulus with a module when the p-value was <0.05. If a correlation had a significance of $p \leq 10^{-6}$, solely this stimuli was listed as associated stimulus even if others had p-values <0.05. If none of the stimuli correlated with $p \leq 10^{-6}$, the two most significant stimuli ($p<0.05$) were used as module-associated stimuli. This approach is inspired by the connection of modules with M1- and M2-stimuli, respectively, described in Xue et al. The stimulus with the most significant correlation is listed first.

| **Module** | **Stimulus** | |
|---|---|---|
| | **positive correlation** | **negative correlation** |
| **1** | LA, OA | IL-13, IL-4 |
| **2** | n.d. | IL-13, IL-4 |
| **3** | sLPS, $M^b$ | IL-13, LiA |
| **4** | OA | IL-13, IL-4 |
| **5** | PA | IL-4, IL-13 |
| **6** | OA, LA | TPP, sLPS + IC |
| **7** | IFN-γ, PA | TPP, TNF+ $PGE_2$ |
| **8** | IFN-γ | TPP* |
| **9** | IFN-γ + TNF, IFN-γ | TPP, TPP + IFN-β |
| **10** | IL-4, TNF | PA, LiA |
| **11** | IL-4 | PA |
| **12** | IL-4, IFN-γ | TPP |
| **13** | IL-4, IL-13 | TPP |
| **14** | IL-4 | sLPS, upLPS + IC |
| **15** | IL-4 | TPP* |
| **16** | PA | IL-4, TPP + IFN-γ |
| **17** | PA | IL-4, IFN-γ + TNF |
| **18** | PA | sLPS, TPP |
| **19** | PA | IL-4, upLPS |
| **20** | PA | TPP |
| **21** | PA | TPP + IFN-β, TPP |
| **22** | OA | TPP + IFN-β + IFN-γ, TPP + IFN-β |
| **23** | OA, LiA | TPP, sLPS |
| **24** | OA | TPP, TPP + IFN-β + IFN-γ |
| **25** | PA | sLPS, TPP |
| **26** | LiA | sLPS, TPP |
| **27** | LA, HDL | sLPS |
| **28** | OA, IL-4 | sLPS |
| **29** | TNF+ $PGE_2$, TPP | $M^b$, IFN-γ |
| **30** | TPP | IL-4, $M^b$ |
| **31** | PA | IL-4, $M^b$ |
| **32** | TPP | IL-4, $M^b$ |
| **33** | TPP | PA, OA |
| **34** | TPP, sLPS + IC | OA, LA |
| **35** | IFN-γ + TNF, sLPS + IFN-γ | OA, LA |
| **36** | TPP, P3C + $PGE_2$ | OA, IFN-γ |
| **37** | IL-4, upLPS | PA |
| **38** | TPP, sLPS + IC | OA |
| **39** | TPP, $PGE_2$ | PA |
| **40** | TPP, upLPS | PA |
| **41** | GC, $PGE_2$ | PA, TNF |
| **42** | GC, $M^b$ | PA, TNF + $PGE_2$ |
| **43** | $M^b$, $PGE_2$ | sLPS, IFN-γ + TNF |
| **44** | OA, GC | IFN-γ, TPP + IFN-β + IFN-γ |
| **45** | IL-4 | PA, sLPS |
| **46** | IL-4 | PA |
| **47** | IL-4, GC | PA |
| **48** | IL-4 | PA, OA |
| **49** | n.d. | n.d. |

*Only one stimulus possessed a p-value<0.05, but $>10^{-6}$ for the module; n.d.: not detected; $M^b$: baseline macrophages

**Table S12: Patient characteristics of TCGA and Rostock cohort**

| **TCGA cohort** (n= 442) | | | |
|---|---|---|---|
| **Parameter** | | **N, value** | **%** |
| Sex | Male | 287 | 65 |
| | Female | 155 | 35 |
| Age (year) | Median (range) | 60 (29-90) | |
| Pathologic T | T1 | 224 | 51 |
| | T2 | 55 | 12 |
| | T3 | 158 | 36 |
| | T4 | 5 | 1 |
| Pathologic N | N0 | 204 | 46 |
| | N1 | 10 | 2 |
| | NX | 228 | 52 |
| Pathologic M | M0 | 353 | 80 |
| | M1 | 64 | 14 |
| | MX | 23 | 5 |
| G | G1 | 9 | 2 |
| | G2 | 188 | 43 |
| | G3 | 177 | 40 |
| | G4 | 65 | 15 |
| | GX | 1 | 0.0 |
| | NA | 2 | 0.0 |
| Follow-up time (year) | Median (range) | 3.6 (0.0 – 12.4) | |
| Cancer-specific survival | Cancer-related death | 87 | 20 |
| | Alive/non-cancer-related death | 355 | 80 |

| **Rostock cohort (Maruschke et al., 2014, Maruschke et al., 2011)** | | | | | | | | |
|---|---|---|---|---|---|---|---|---|
| **Patient** | **Tumor** | **Age/** | **Tumor** | **Tumor stage** | **Nuclear** | **Follow-** | **Outcome** | **Normal** |
| **Primary tumor G1** | | | | | | | | |
| 12 | ccRCC | 72/female | 3.0 | pT2N0 M0V0R0 | G1 | 148 | ANED* | X |
| 113 | ccRCC | 59/male | 4.0 | pT1 N0M0V0R0 | G1 | 133 | ANED | X |
| 158 | ccRCC | 72/male | 3.2 | pT2 N0M0V0R0 | G1 | 126 | AWD | X |
| 34 | ccRCC | 65/male | 5.2 | pT2N0M0V0R0 | G1 | 144 | ANED | X |
| 348 | ccRCC | 59/male | 8.0 | pT2N0M0V0R0 | G1 | 67 | ANED | X |
| 365 | ccRCC | 62/female | 5.5 | pT2N0M0V0R0 | G1 | 65 | ANED | X |
| 165 | ccRCC | 77/male | 4.5 | pT1N0M0V0R0 | G1 | 125 | ANED | X |
| 198 | ccRCC | 70/male | 3.8 | pT1N0M0V0R0 | G1 | 97 | DBOR | X |
| 262 | ccRCC | 71/female | 5.0 | pT1N0M0V0R0 | G1 | 96 | ANED | |
| 269 | ccRCC | 53/female | 8.0 | pT3aN0M0V0R0 | G1 | 94 | ANED | |
| 286 | ccRCC | 65/male | 5.5 | pT1N0M0V0R1 | G1 | 93 | ANED | |
| 288 | ccRCC | 70/female | 5.5 | pT1N0M0V0R0 | G1 | 90 | ANED | |
| 290 | ccRCC | 56/male | 3.5 | pT1N0M0V0R0 | G1 | 89 | ANED | |
| 294 | ccRCC | 62/female | 3.0 | pT1N0M0V0R0 | G1 | 76 | DBOR | |
| **Primary tumor G3** | | | | | | | | |
| 89 | ccRCC | 64/male | 11.0 | pT3bN2M1V1R0 | G3 | 2 | DOTD | |
| 188 | ccRCC | 59/male | 9.0 | pT2 N0M1V0R0 | G3 | 30 | DOTD | X |
| 197 | ccRCC | 63/male | 13.0 | pT3bN1M1V1R0 | G3 | 23 | DOTD | X |

| | | | | | | | | |
|---|---|---|---|---|---|---|---|---|
| 254 | ccRCC | 62/male | 12.0 | pT3bN0M1V1R0 | G3 | 29 | DOTD | |
| 382 | ccRCC | 56/male | 8.0 | pT3bN0M1V1R0 | G3 | 60 | AWD | X |
| 351 | ccRCC | 66/female | 10.5 | pT3bN1M1V1R0 | G3 | 11 | DOTD | X |
| 168 | ccRCC | 58/male | 12.0 | pT3aN0M0V1R1 | G3 | 46 | DOTD | |
| 190 | ccRCC | 56/male | 8.6 | pT3bN0M0V1R0 | G3 | 31 | DOTD | X |
| 230 | ccRCC | 55/female | 4.0 | pT3bN0M0V1R1 | G3 | 15 | DOTD | |
| 345 | ccRCC | 59/male | 13.0 | pT3aNXM1V0R0 | G3 | 60 | AWD | |
| 399 | ccRCC | 65/female | 7.5 | pT3aN0M1V0R0 | G3 | 15 | DOTD | |
| 510 | ccRCC | 79/female | 5.0 | pT1aN0M1V1R0 | G3 | 3 | AWD | |
| 283 | ccRCC | 48/male | 5.2 | pT1N0M1V1R0 | G3 | 94 | AWD | |
| 390 | ccRCC | 59/female | 16.0 | pT3bN0M1V1R1 | G3 | 2 | DOTD | X |

* ANED = alive with no evidence of disease, AWD = alive with disease, DOTD = death of tumor disease, DBOR = death of other reason

**Table S13: Antibodies.** Antibody concentrations for FACS sorting were adjusted to the total cell number of each sample. Isotypes were adjusted to the concentration of respective specific antibody.

| Antibody | Label | Species/Isotype | Clon | Company |
|---|---|---|---|---|
| B7-DC/PD-L2 | APC | Mouse IgG1, κ | MIH18 | BD |
| B7-H1/PD-L1 | FITC | Mouse IgG1, κ | M1H1 | BD |
| B7-H3 | APC | Mouse IgG2b, κ | FM276 | Miltenyi Biotec |
| CCR1 | PE | Mouse IgG2b | 53504.111 | R&D Systems |
| CCR2 | PE | Mouse IgG2b | #48607.211 | R&D Systems |
| CCR5 | FITC | Mouse IgG2a | 2D7/CCR5 | BD |
| CCR7 | APC-A750 | Rat IgG2a | 3D12 | eBioscience |
| CD1c | FITC | Mouse IgG2a | AD5-8E7 | Miltenyi Biotec |
| CD1c | PE-Cy7 | Mouse IgG1, κ | L161 | Biolegend |
| CD11c | APC | Mouse IgG1 | B-ly6 | BD |
| CD11c | A700 | Mouse IgG1 | B-ly6 | BD |
| CD11c | PE | Mouse IgG1, κ | B-ly6 | BD |
| CD11b | FITC | Mouse IgG1 | ICRF44 | eBioscience |
| CD13/ANPEP | APC | Mouse IgG1, κ | WM15 | BD |
| CD14 | PB | Maus IgG2a, κ | M5E2 | BD |
| CD14 | PerCP-Cy5.5 | Mouse IgG1, κ | 61D3 | eBioscience |
| CD19 | PB | Mouse IgG1, κ | HD37 | Dako |
| CD204/MSR1 | FITC | rec. human IgG1 | REA460 | Miltenyi Biotec |
| CD206/MRC1 | FITC | Mouse IgG1, κ | 15-2 | Biolgend |
| CD209 | APC | Mouse IgG2b, κ | DCN46 | BD |
| CD209 | PE | Mouse IgG2b, κ | DCN46 | BD |
| CD3 | PB | Mouse IgG1, κ | UCHT1 | BD |
| CD3 | V500 | Mouse IgG1 | UCHT1 | BD |
| CD32A | unlabeled | Goat IgG | Polyclonal | R&D Systems |
| CD36 | FITC | Mouse IgG2a | FA6-152 | Immunotech |
| CD40 | FITC | Mouse IgG1, κ | 5C3 | BD |
| CD45 | PE-Cy7 | Mouse IgG1, κ | HI30 | BD |
| CD45 | APC Cy7 | Mouse IgG1, κ | HI30 | BD |
| CD56 | V450 | Mouse IgG1 | B159 | BD |
| CD64 | FITC | Mouse IgG1, κ | 22 | Beckman Coulter |
| CD64 | PerCP Cy5.5 | Mouse IgG1 | 10.1 | BioLegend |
| CD68 | FITC | Mouse IgG | Y1/82A | BD |
| CD141 | APC | Mouse IgG1 | AD5-14H12 | Miltenyi Biotech |
| CD163 | PerCP Cy5.5 | Mouse IgG1, κ | GHI/61 | BioLegend |
| CD80 | A700 | Mouse IgG1 | L.307.4 | BD |
| CD83 | PE | Mouse IgG2b | HB15a | Immunotech |
| CD86 | PE | Mouse IgG1 | 2331(FUN-1) | BD |
| CD9 | FITC | Mouse IgG1 | MEM-61 | EuroBioSciences |
| CSF1R | A700 | Mouse IgG1 | 61708 | R&D Systems |
| $CX_3CR1$ | APC | Rat IgG2b, κ | 2A9-1 | eBioscience |
| FLT3 | APC | Mouse IgG1 | BV10A4H2 | eBioscience |

| | | | | |
|---|---|---|---|---|
| GPNMB | A700 | Mouse IgG2b | 303822 | Novus Biologicals |
| HLA-DR/ MHC-II | PE | Mouse IgG2a, κ | G46-6 | BD |
| Isotype | A700 | Mouse IgG1, κ | 11711 | R&D Systems |
| Isotype | APC | Mouse IgG1, κ | 11711 | R&D Systems |
| Isotype | APC | Mouse IgG2b, κ | 27-35 | BD |
| Isotype | FITC | Mouse IgG1, κ | MOPC21 | BD |
| Isotype | PE | Mouse IgG1 | MOPC21 | BD |
| Isotype | PE | Mouse IgG2a, κ | MOPC-173 | BioLegend |
| Isotype | PE | Mouse IgG2b, κ | MPC-11 | BioLegend |
| Isotype | PE-Cy7 | Mouse IgG1, κ | MOPC21 | BD |
| Isotype | FITC | Mouse IgM | IS5-20C4 | Miltenyi Biotec |
| MerTK | APC | Mouse IgG1, κ | 125518 | R&D Sytems |
| NRP1 | FITC | Mouse IgG1 | AD5-17F6 | Miltenyi Biotec |
| Slan/M-DC8 | FITC | Mouse IgM | DD-1 | Miltenyi Biotec |
| TIM-3 | FITC | Rat IgG2a | 344823 | R&D Systems |
| VSIG4 | unlabled | Rabbit | polyclonal | Abcam |
| Anti-goat | A488 | Donkey IgG | polyclonal | Thermo Fisher Scientific |
| Anti-mouse IgG | A647 | Donkey | polyclonal | Jackson ImmuneResearch |
| Anti-mouse IgG1 | A488 | Goat | polyclonal | Thermo Fisher Scientific |
| Anti-rabbit IgG | A488 | Goat | polyclonal | Thermo Fisher Scientific |

### Supplemental experimental procedures

#### Characteristics of human myeloid cell types from different non-lymphoid tissues used for comparison with the ercDC transcriptome

Renal DCs and macrophages are thought to play a role in protecting the kidney tubulointerstitium from immune attack-induced injury through T-cell tolerizing mechanisms (Kurts et al., 2007). To gain insight if ercDCs might exhibit tolerizing features associated with tissue/tumor immune protection, we thought to compare the ercDC transcriptome to that of other myeloid cell types originating from tissues with a known tolerogenic milieu (**table S2B**). Of particular interest were the CD11c$^{HI}$ and CD11c$^{LO}$ decidual macrophages (previously designated decidual DCs) (Houser et al., 2011), found in the tolerogenic placenta during early pregnancy. Like ercDCs, they are decribed to co-express CD209, CD14 as well as CD163, and seemed to be capable of antigen presentation, CD11c$^{HI}$ macrophages mainly expressed genes associated with lipid metabolism and inflammation, while CD11c$^{LO}$ macrophages exhibited strong expression of genes associated with tissue modulation, which are important for embryo implantation, but also for tumor growth and progression.

As an additional data set we selected transcriptomes of myeloid cells from gut and lung, considering that in these organs, the immune system is important to maintain the balance between immunity and peripheral tolerance. Three different DC subtypes are described in the lamina propria distinguished based on the expression of CD103 and Sirpα (Watchmaker et al., 2014)). CD103$^{+}$Sirpα$^{-}$ DCs are described to resemble CD141$^{+}$ DCs from blood and express marker associated with crosspresentation; CD103$^{+}$Sirpα$^{+}$ DCs have a described similarity with CD1c$^{+}$ DCs from blood. They are known to induce $T_H17$ cells and express CD209 and genes associated with tolerizing functions. The third population, CD103$^{-}$Sirpα$^{+}$ DCs, has strong similarity with monocyte-derived DCs (MoDCs) and is thought to effectively expand $T_H1$ cells. Concering myeloid cells from lung, two data sets were available (Shaykhiev et al., 2009)(Woodruff et al., 2005). They included macrophages isolated from non-smokers, smokers and COPD or asthma patients, respectively. Macrophages from smokers and COPD patients are described to express features associated with downregulating M1-associated proinflammatory genes and upregulating M2-associated regulatory genes compared to macrophages from non-smokers (Shaykhiev et al., 2009). They expressed high amounts of MMPs, a characteristic also described for macrophages in the tumors.

Further, we selected data sets of human tumor samples. These included, macrophages from gastrointestinal stromal tumors (GIST) (Cavnar et al., 2013) and inflammatory macrophages and DCs from ascites of ovarian cancer patients (Segura and Amigorena, 2013) Notably, the macrophages from GIST are described to have antitumoral, proinflammatory features resembling M1-macrophages. As representative for DCs with the capability of crosspresention and activation of CD8$^{+}$ T cells (Bachem et al., 2010, Mittag et al., 2011) we selected the data sets of CD141$^{+}$ DCs from Lindstedt et al. (Lindstedt et al., 2005).
