## Supplementary material for "A mosaic renal myeloid subtype with T-cell inhibitory and protumoral features is linked to immune escape and survival in clear cell renal cell cancer": Brech et al_supplementary Table S8

**Table S8:** Upregulated ercDC_ccRCC DEGs. Listed are genes significantly (adjusted p<0.05) upregulated between ercDC_ccRCC&infMΦ_ascOvCa and control group (431 of 788 ercDC_ccRCC DEGs). Genes are sorted acc. to increasing values. Genes that also belong to ercDC_ccRCC marker genes are in bold green. Genes related to ercDC_ccRCC marker ge Genes mentioned in main text are in grey.

| Gene symbol | Entrez ID | norm. expr.values (log2) ercDC_ccRCC&infMΦ_ascOvCa | norm. expr.values (log2) control group | logFC |
|---|---|---|---|---|
| **APOC1** | 341 | 11,82 | 10,04 | 1,78 |
| **FOLR2** | 2350 | 8,7 | 7,31 | 1,39 |
| **GAL3ST4** | 79690 | 8,01 | 6,59 | 1,42 |
| **SDC3** | 9672 | 7,65 | 6,42 | 1,23 |
| **ME1** | 4199 | 9,59 | 8,13 | 1,46 |
| **C2** | 717 | 8,52 | 7,07 | 1,46 |
| **CCL8** | 6355 | 8,53 | 6,57 | 1,96 |
| **CD209** | 30835 | 8,44 | 7,05 | 1,39 |
| **PLTP** | 5360 | 9,71 | 7,71 | 2 |
| SLC2A5 | 6518 | 6,85 | 5,53 | 1,32 |
| ITGA9 | 3680 | 6,19 | 5,28 | 0,9 |
| **SLCO2B1** | 11309 | 9,65 | 7,79 | 1,86 |
| **GPNMB** | 10457 | 11,22 | 9,09 | 2,13 |
| **ABCC5** | 10057 | 7,94 | 7,1 | 0,84 |
| AP2A2 | 161 | 8,97 | 8,28 | 0,69 |
| TNS3 | 64759 | 9,41 | 8,79 | 0,63 |
| **SEPP1** | 6414 | 9,11 | 6,69 | 2,42 |
| **NRP2** | 8828 | 8,12 | 6,62 | 1,5 |
| CXCL12 | 6387 | 6,52 | 5,73 | 0,79 |
| LHFPL2 | 10184 | 10,97 | 9,58 | 1,38 |
| **SLC38A6** | 145389 | 10,17 | 9,02 | 1,15 |
| CMKLR1 | 1240 | 8,61 | 7,16 | 1,46 |
| OLFML2B | 25903 | 7,65 | 6,34 | 1,31 |
| **NRP1** | 8829 | 9,98 | 8,25 | 1,73 |
| **SLC7A8** | 23428 | 7,9 | 6,27 | 1,63 |
| SLC38A7 | 55238 | 7,49 | 6,79 | 0,7 |
| LILRB5 | 10990 | 6,87 | 5,9 | 0,97 |
| **NUPR1** | 26471 | 9,85 | 8,51 | 1,34 |
| ACP2 | 53 | 9,95 | 9,05 | 0,91 |
| EPHB2 | 2048 | 6,71 | 5,8 | 0,91 |
| WBP5 | 51186 | 8,96 | 7,98 | 0,98 |
| CTSL | 1514 | 12,28 | 10,97 | 1,3 |
| SLC36A1 | 206358 | 8,1 | 7,41 | 0,69 |
| **FABP3** | 2170 | 8,4 | 6,86 | 1,54 |
| **DAB2** | 1601 | 9,9 | 8,39 | 1,51 |
| MMP14 | 4323 | 8,05 | 6,92 | 1,14 |
| KAL1 | 3730 | 8,95 | 7,78 | 1,16 |
| **APOE** | 348 | 11,05 | 9,23 | 1,82 |
| ABCA1 | 19 | 9,63 | 8,46 | 1,17 |

| | | | | |
|---|---|---|---|---|
| FRMD4A | 55691 | 7,92 | 6,81 | 1,11 |
| TEC | 7006 | 6,45 | 5,96 | 0,49 |
| **SIGLEC1** | 6614 | 9,68 | 8,23 | 1,45 |
| **CCL18** | 6362 | 11,42 | 9,62 | 1,8 |
| **ADAMDEC1** | 27299 | 9,06 | 7,33 | 1,73 |
| **C1QA** | 712 | 11,83 | 10,41 | 1,42 |
| DOCK4 | 9732 | 9,29 | 8,04 | 1,25 |
| **CRYAB** | 1410 | 6,75 | 5,68 | 1,08 |
| NR1H3 | 10062 | 9,01 | 7,95 | 1,06 |
| RAB3IL1 | 5866 | 7,3 | 6,76 | 0,54 |
| TREM2 | 54209 | 9,99 | 8,39 | 1,6 |
| PLOD2 | 5352 | 5,22 | 4,27 | 0,95 |
| SLC1A3 | 6507 | 8,06 | 6,67 | 1,39 |
| **RHOBTB3** | 22836 | 7,21 | 6,07 | 1,14 |
| DRAM1 | 55332 | 10,26 | 9,37 | 0,89 |
| HS3ST1 | 9957 | 6,9 | 5,74 | 1,16 |
| **MAF** | 4094 | 8,17 | 6,88 | 1,28 |
| **C1QB** | 713 | 12,33 | 10,92 | 1,42 |
| CD64 | 2209 | 6,96 | 5,79 | 1,18 |
| **IL2RA** | 3559 | 6,85 | 5,69 | 1,16 |
| **ARHGAP12** | 94134 | 7,61 | 6,79 | 0,82 |
| PSD3 | 23362 | 7,92 | 6,81 | 1,11 |
| TNFRSF11A | 8792 | 6,77 | 5,81 | 0,96 |
| CTSB | 1508 | 12,23 | 11,32 | 0,91 |
| ADAP2 | 55803 | 9,85 | 9,02 | 0,83 |
| CFH | 3075 | 5,72 | 5,09 | 0,63 |
| RNASE1 | 6035 | 10,53 | 8,67 | 1,86 |
| ITGAV | 3685 | 10,5 | 9,47 | 1,03 |
| **SCAMP5** | 192683 | 6,18 | 5,54 | 0,64 |
| TCN2 | 6948 | 8,26 | 7,45 | 0,81 |
| TDRKH | 11022 | 6,33 | 5,65 | 0,68 |
| **SERPING1** | 710 | 11,43 | 9,99 | 1,45 |
| IL10 | 3586 | 6,98 | 5,89 | 1,1 |
| MMP19 | 4327 | 9,5 | 8,23 | 1,27 |
| **MS4A4A** | 51338 | 10,52 | 9,3 | 1,21 |
| TMEM51 | 55092 | 8,87 | 7,95 | 0,92 |
| MYO7A | 4647 | 5,97 | 5,27 | 0,71 |
| ZFYVE26 | 23503 | 9,17 | 8,55 | 0,62 |
| ITSN1 | 6453 | 7,87 | 6,9 | 0,97 |
| SCARB2 | 950 | 9,58 | 8,67 | 0,91 |
| CFB | 629 | 8,06 | 6,82 | 1,25 |
| EDNRB | 1910 | 5,53 | 4,79 | 0,74 |
| CD163 | 9332 | 8,32 | 6,79 | 1,53 |
| PDCD1LG2 | 80380 | 7,64 | 6,48 | 1,16 |

| | | | | |
|---|---|---|---|---|
| STAB1 | 23166 | 8,69 | 7,56 | 1,13 |
| ADAM9 | 8754 | 10,03 | 9,1 | 0,93 |
| SNX24 | 28966 | 7,07 | 6,31 | 0,76 |
| CD81 | 975 | 12,34 | 11,6 | 0,75 |
| CD28 | 940 | 4,91 | 4,28 | 0,63 |
| FLCN | 201163 | 6,85 | 6,34 | 0,51 |
| C3 | 718 | 9,97 | 8,43 | 1,54 |
| MITF | 4286 | 8,4 | 7,7 | 0,7 |
| CRYBB1 | 1414 | 6,2 | 5,7 | 0,5 |
| EPAS1 | 2034 | 9,11 | 7,93 | 1,18 |
| GLUL | 2752 | 10,8 | 9,74 | 1,06 |
| MERTK | 10461 | 7,93 | 6,5 | 1,43 |
| ADORA3 | 140 | 7,24 | 6,02 | 1,22 |
| TRPV4 | 59341 | 6,56 | 5,98 | 0,57 |
| PTPRM | 5797 | 8,37 | 7,57 | 0,79 |
| BMP2K | 55589 | 8,97 | 8,16 | 0,81 |
| APPL2 | 55198 | 7,39 | 6,79 | 0,6 |
| KIAA0226L | 80183 | 8,69 | 7,6 | 1,09 |
| NPC1 | 4864 | 9,11 | 8,04 | 1,07 |
| LAIR1 | 3903 | 9,89 | 8,86 | 1,02 |
| VAT1 | 10493 | 10,31 | 9,55 | 0,76 |
| LGALS3BP | 3959 | 10,19 | 9,19 | 1 |
| GNPDA1 | 10007 | 9,75 | 9,19 | 0,56 |
| IDH1 | 3417 | 10,69 | 9,84 | 0,85 |
| HNMT | 3176 | 9,35 | 8,36 | 1 |
| **LGMN** | 5641 | 11,67 | 10,11 | 1,56 |
| CD84 | 8832 | 9,11 | 7,84 | 1,27 |
| SGMS1 | 259230 | 8,66 | 7,91 | 0,75 |
| CTSD | 1509 | 11,32 | 10,42 | 0,9 |
| BCL2L1 | 598 | 7,62 | 6,72 | 0,9 |
| STX4 | 6810 | 9,78 | 9,24 | 0,53 |
| SDS | 10993 | 7,87 | 6,8 | 1,07 |
| BNIP3 | 664 | 8,06 | 7,01 | 1,05 |
| SPP1 | 6696 | 11,15 | 8,64 | 2,51 |
| MPP1 | 4354 | 10,58 | 9,99 | 0,59 |
| | 3569 | 7,68 | 6,29 | 1,39 |
| NPL | 80896 | 9 | 7,94 | 1,06 |
| STARD13 | 90627 | 6,22 | 5,7 | 0,52 |
| FCGR1B | 2210 | 11,09 | 9,8 | 1,29 |
| MARCKS | 4082 | 10,25 | 8,94 | 1,31 |
| TCF12 | 6938 | 9,19 | 8,63 | 0,56 |
| PLA2G15 | 23659 | 8,76 | 8,07 | 0,68 |
| CCL4 | 6351 | 11,32 | 9,56 | 1,76 |
| VSIG4 | 11326 | 11,86 | 10,59 | 1,27 |

| | | | | |
|---|---|---|---|---|
| CCL2 | 6347 | 10,46 | 8,71 | 1,76 |
| MGAT4A | 11320 | 9,3 | 8,1 | 1,2 |
| RND3 | 390 | 8,82 | 7,69 | 1,13 |
| RGL1 | 23179 | 9,28 | 8,17 | 1,11 |
| LIPA | 3988 | 12,69 | 12 | 0,69 |
| COLGALT1 | 79709 | 8,91 | 8,32 | 0,59 |
| CREG1 | 8804 | 12,04 | 11,5 | 0,53 |
| LRP1 | 4035 | 8,64 | 7,99 | 0,65 |
| CYFIP1 | 23191 | 11,25 | 10,73 | 0,52 |
| CPM | 1368 | 9,7 | 8,68 | 1,02 |
| CALU | 813 | 8,49 | 7,88 | 0,61 |
| MYO5A | 4644 | 8,96 | 8,43 | 0,53 |
| **IGF1** | 3479 | 7,08 | 5,85 | 1,23 |
| MAPK13 | 5603 | 8,2 | 7,49 | 0,72 |
| GALC | 2581 | 9,19 | 8,47 | 0,72 |
| FZD5 | 7855 | 7,25 | 6,59 | 0,67 |
| SLC6A8 | 6535 | 6,72 | 6,19 | 0,53 |
| DYNLT3 | 6990 | 10,21 | 9,72 | 0,49 |
| PLIN2 | 123 | 10,84 | 9,93 | 0,92 |
| METTL1 | 4234 | 7,44 | 6,86 | 0,58 |
| EYA2 | 2139 | 5,29 | 4,87 | 0,42 |
| CXCL10 | 3627 | 10,24 | 8,15 | 2,08 |
| CXCL2 | 2920 | 10,79 | 9,38 | 1,41 |
| TFRC | 7037 | 10,35 | 9,33 | 1,03 |
| PLXNA3 | 55558 | 6,46 | 5,86 | 0,6 |
| ATP13A2 | 23400 | 7,37 | 6,72 | 0,65 |
| LYVE1 | 10894 | 6,39 | 5,4 | 1 |
| OLR1 | 4973 | 11 | 9,4 | 1,6 |
| RBM47 | 54502 | 9,35 | 8,69 | 0,65 |
| FN1 | 2335 | 10,06 | 8,12 | 1,93 |
| MARCO | 8685 | 10,44 | 9,42 | 1,03 |
| FAM13A | 10144 | 8,11 | 7,33 | 0,78 |
| AP1B1 | 162 | 8,95 | 8,46 | 0,49 |
| CAMSAP2 | 23271 | 7,83 | 7,03 | 0,8 |
| IGFBP4 | 3487 | 7,04 | 6,45 | 0,59 |
| P4HA2 | 8974 | 7,5 | 6,97 | 0,53 |
| ATP6V1C1 | 528 | 8,56 | 7,86 | 0,7 |
| ENOSF1 | 55556 | 8,16 | 7,49 | 0,67 |
| GPR65 | 8477 | 10,49 | 9,78 | 0,71 |
| OLFML3 | 56944 | 6,1 | 5,5 | 0,6 |
| SLC11A2 | 4891 | 8,37 | 7,79 | 0,57 |
| PLAT | 5327 | 5,3 | 4,77 | 0,52 |
| PLXND1 | 23129 | 9,37 | 8,85 | 0,52 |
| CTSZ | 1522 | 11,03 | 10,49 | 0,54 |

| | | | | |
|---|---|---|---|---|
| CXCL1 | 2919 | 8,55 | 7,33 | 1,22 |
| ETV5 | 2119 | 7,67 | 6,56 | 1,11 |
| DNASE2 | 1777 | 8,54 | 7,76 | 0,79 |
| CD204 | 4481 | 9,74 | 8,14 | 1,6 |
| IFI27 | 3429 | 9,65 | 8,19 | 1,47 |
| ASPH | 444 | 8,16 | 7,41 | 0,75 |
| PLAU | 5328 | 8,77 | 7,62 | 1,15 |
| HOMER3 | 9454 | 7,32 | 6,71 | 0,61 |
| ITPR2 | 3709 | 9,18 | 8,33 | 0,85 |
| FABP5 | 2171 | 11,79 | 10,75 | 1,03 |
| FNDC3B | 64778 | 8,84 | 8,13 | 0,71 |
| GPR137B | 7107 | 10,84 | 10,08 | 0,76 |
| ANGPTL4 | 51129 | 6,29 | 5,73 | 0,56 |
| TIMP2 | 7077 | 10,99 | 10,48 | 0,51 |
| GLA | 2717 | 10,87 | 10,29 | 0,58 |
| PI4K2A | 55361 | 8,75 | 8,17 | 0,58 |
| KCNMA1 | 3778 | 8,67 | 7,51 | 1,15 |
| ELL2 | 22936 | 8,79 | 7,84 | 0,96 |
| MMP2 | 4313 | 7,67 | 6,67 | 0,99 |
| DYRK4 | 8798 | 7,75 | 7,32 | 0,43 |
| CP | 1356 | 5,03 | 4,2 | 0,83 |
| A2M | 2 | 11,18 | 9,48 | 1,7 |
| FRMD4B | 23150 | 8,6 | 7,67 | 0,93 |
| PLEKHO2 | 80301 | 9,95 | 9,46 | 0,49 |
| HAMP | 57817 | 8,1 | 6,82 | 1,29 |
| ADAMTS2 | 9509 | 5,74 | 5,3 | 0,44 |
| SCIN | 85477 | 5,25 | 4,52 | 0,73 |
| IQCG | 84223 | 7,12 | 6,69 | 0,43 |
| LINC00597 | 81698 | 5,16 | 4,73 | 0,43 |
| LILRB4 | 11006 | 9,02 | 8,24 | 0,77 |
| PEAK1 | 79834 | 8,04 | 7,55 | 0,49 |
| SLC16A10 | 117247 | 7,59 | 6,48 | 1,11 |
| CTSA | 5476 | 11,29 | 10,74 | 0,55 |
| ACE | 1636 | 6,14 | 5,66 | 0,48 |
| ATG7 | 10533 | 7,31 | 6,79 | 0,52 |
| MKNK1 | 8569 | 9,51 | 8,86 | 0,65 |
| PLD3 | 23646 | 10,31 | 9,49 | 0,82 |
| CXCL11 | 6373 | 7,07 | 5,48 | 1,59 |
| SGPL1 | 8879 | 8,52 | 7,98 | 0,54 |
| FPR3 | 2359 | 10,57 | 9,15 | 1,42 |
| RCN3 | 57333 | 6,72 | 6,32 | 0,41 |
| PMP22 | 5376 | 10,3 | 9,2 | 1,1 |
| PROS1 | 5627 | 8,22 | 7,57 | 0,65 |
| RAB13 | 5872 | 10,31 | 9,57 | 0,74 |

| | | | | |
|---|---|---|---|---|
| CDR1 | 1038 | 4,36 | 3,84 | 0,51 |
| ECM1 | 1893 | 7,47 | 6,77 | 0,71 |
| WASF1 | 8936 | 4,81 | 4,31 | 0,5 |
| ATP2A2 | 488 | 8,51 | 8,07 | 0,44 |
| CADM1 | 23705 | 7,64 | 6,36 | 1,28 |
| FARP1 | 10160 | 7,68 | 7,15 | 0,53 |
| WDFY3 | 23001 | 7,36 | 6,77 | 0,59 |
| SLC37A4 | 2542 | 7,25 | 6,86 | 0,38 |
| RNASE2 | 6036 | 7,92 | 6,95 | 0,97 |
| ALG9 | 79796 | 7,64 | 7,2 | 0,44 |
| HSPB1 | 3315 | 10,96 | 10,13 | 0,83 |
| HSP90B1 | 7184 | 9,34 | 8,67 | 0,67 |
| ABCG1 | 9619 | 8,03 | 7,24 | 0,79 |
| DENND2D | 79961 | 9 | 8,54 | 0,46 |
| PDIA5 | 10954 | 7,16 | 6,67 | 0,49 |
| MMP9 | 4318 | 9,81 | 8,47 | 1,34 |
| SAMD4A | 23034 | 7,76 | 7,11 | 0,65 |
| PPARG | 5468 | 9,26 | 8,4 | 0,86 |
| TNS1 | 7145 | 8,89 | 7,65 | 1,23 |
| SCCPDH | 51097 | 8,5 | 7,94 | 0,56 |
| CH25H | 9023 | 7,89 | 6,63 | 1,26 |
| ADM | 133 | 9,08 | 8,12 | 0,96 |
| ST8SIA4 | 7903 | 7,17 | 6,48 | 0,69 |
| SCD | 6319 | 9,38 | 8,19 | 1,19 |
| ACP5 | 54 | 11,01 | 10,19 | 0,82 |
| PCOLCE2 | 26577 | 8,39 | 7,58 | 0,81 |
| Sep 08 | 23176 | 6,95 | 6,52 | 0,43 |
| PLEKHM2 | 23207 | 8,46 | 8,04 | 0,42 |
| **CXCL9** | 4283 | 9,73 | 7,88 | 1,85 |
| IL15RA | 3601 | 8,23 | 7,64 | 0,6 |
| APBB3 | 10307 | 7,35 | 6,91 | 0,45 |
| ZFP36L1 | 677 | 9,84 | 9,12 | 0,72 |
| AMPD3 | 272 | 8,78 | 8,03 | 0,74 |
| SLC4A7 | 9497 | 7,41 | 6,78 | 0,63 |
| TBC1D2 | 55357 | 8,5 | 7,89 | 0,6 |
| SLC2A8 | 29988 | 6,73 | 6,27 | 0,46 |
| EOGT | 285203 | 7,95 | 7,36 | 0,59 |
| NFE2L1 | 4779 | 9,28 | 8,84 | 0,44 |
| LPAR6 | 10161 | 10,06 | 9,23 | 0,83 |
| RENBP | 5973 | 7,58 | 7,07 | 0,51 |
| FUCA1 | 2517 | 11,39 | 10,57 | 0,82 |
| MAFB | 9935 | 11,67 | 10,74 | 0,93 |
| ZDHHC14 | 79683 | 7,2 | 6,72 | 0,48 |
| BAMBI | 25805 | 6 | 5,5 | 0,5 |

| | | | | |
|---|---|---|---|---|
| ABCC3 | 8714 | 8,43 | 7,69 | 0,73 |
| TPX2 | 22974 | 6,15 | 5,56 | 0,59 |
| GBAP1 | 2630 | 8,1 | 7,74 | 0,37 |
| GNPTAB | 79158 | 8,78 | 8,29 | 0,49 |
| CXCL3 | 2921 | 9,27 | 8,22 | 1,05 |
| CTSC | 1075 | 10,9 | 10,12 | 0,78 |
| CD82 | 3732 | 8,49 | 7,84 | 0,65 |
| RACGAP1 | 29127 | 7,72 | 7,16 | 0,55 |
| P2RX4 | 5025 | 9,7 | 9,03 | 0,67 |
| TGFBI | 7045 | 12,22 | 11,66 | 0,56 |
| LXN | 56925 | 7,38 | 6,86 | 0,52 |
| PPAP2B | 8613 | 9 | 8 | 1 |
| SPHK1 | 8877 | 7,41 | 6,77 | 0,65 |
| P4HB | 5034 | 9,86 | 9,14 | 0,72 |
| IER3 | 8870 | 11,54 | 10,32 | 1,21 |
| PLOD1 | 5351 | 8,6 | 8,22 | 0,38 |
| ITGB5 | 3693 | 6,83 | 6,05 | 0,77 |
| TMEM140 | 55281 | 8,75 | 8,22 | 0,53 |
| ATP6V0A1 | 535 | 8,86 | 8,32 | 0,54 |
| BNC2 | 54796 | 5,75 | 5,25 | 0,49 |
| NENF | 29937 | 8,57 | 8,18 | 0,39 |
| RRAGD | 58528 | 9,86 | 9,27 | 0,6 |
| CD59 | 966 | 8,6 | 7,77 | 0,83 |
| ACVRL1 | 94 | 7,62 | 7,1 | 0,52 |
| GAA | 2548 | 9,63 | 9,14 | 0,48 |
| CD80 | 941 | 8,19 | 7,1 | 1,09 |
| CTNS | 1497 | 8,43 | 7,83 | 0,6 |
| SLC31A1 | 1317 | 9,1 | 8,56 | 0,54 |
| NABP1 | 64859 | 9,35 | 8,72 | 0,63 |
| PDIA4 | 9601 | 9,21 | 8,68 | 0,54 |
| HSPA5 | 3309 | 10,33 | 9,79 | 0,54 |
| COLEC12 | 81035 | 8,99 | 8,14 | 0,85 |
| MDFIC | 29969 | 8,23 | 7,58 | 0,65 |
| BLNK | 29760 | 9,14 | 8,1 | 1,04 |
| MMP7 | 4316 | 6,57 | 5,61 | 0,96 |
| AAK1 | 22848 | 8,95 | 8,53 | 0,42 |
| CCL7 | 6354 | 6,43 | 5,57 | 0,86 |
| ARHGAP6 | 395 | 7,24 | 6,78 | 0,46 |
| FCGRT | 2217 | 10,71 | 10,24 | 0,48 |
| EPHX1 | 2052 | 8,02 | 7,43 | 0,58 |
| B2M | 567 | 9,52 | 9,13 | 0,39 |
| TRIP6 | 7205 | 8,46 | 8,06 | 0,4 |
| CD14 | 929 | 11,49 | 10,52 | 0,97 |
| ATXN1 | 6310 | 8,59 | 7,92 | 0,67 |

| | | | | |
|---|---|---|---|---|
| HTRA1 | 5654 | 7,6 | 6,95 | 0,65 |
| LAMP1 | 3916 | 9,72 | 8,81 | 0,91 |
| GGCX | 2677 | 7,5 | 7,12 | 0,37 |
| GATM | 2628 | 8,5 | 7,42 | 1,08 |
| PXDC1 | 221749 | 7,73 | 7,07 | 0,66 |
| SERPINE1 | 5054 | 7,06 | 6,24 | 0,83 |
| IKBKE | 9641 | 7,31 | 6,93 | 0,38 |
| DGKH | 160851 | 5,47 | 5,1 | 0,37 |
| PAX8 | 7849 | 6,53 | 5,95 | 0,58 |
| PRUNE2 | 158471 | 6,94 | 5,97 | 0,96 |
| ENG | 2022 | 9,2 | 8,5 | 0,7 |
| C1R | 715 | 6,65 | 6,08 | 0,57 |
| CD2AP | 23607 | 7,79 | 7,24 | 0,56 |
| TUBG1 | 7283 | 8,36 | 7,89 | 0,48 |
| KIAA1279 | 26128 | 7,78 | 7,38 | 0,4 |
| ERI2 | 112479 | 7,34 | 6,93 | 0,42 |
| AHI1 | 54806 | 7,36 | 6,7 | 0,65 |
| MSRB2 | 22921 | 8,5 | 8,06 | 0,44 |
| DHRS3 | 9249 | 9,01 | 8,16 | 0,85 |
| SERPINH1 | 871 | 7,86 | 7,23 | 0,63 |
| MR1 | 3140 | 7,46 | 6,96 | 0,5 |
| ME2 | 4200 | 9,09 | 8,6 | 0,5 |
| DSC2 | 1824 | 8,48 | 7,6 | 0,88 |
| LAMP2 | 3920 | 9,89 | 9,34 | 0,55 |
| MORF4L2 | 9643 | 6,95 | 6,43 | 0,52 |
| CA12 | 771 | 6,19 | 5,47 | 0,72 |
| HSD17B14 | 51171 | 7,13 | 6,61 | 0,51 |
| QKI | 9444 | 8,66 | 8,14 | 0,52 |
| RARRES1 | 5918 | 7,02 | 6,12 | 0,9 |
| CALR | 811 | 10,03 | 9,35 | 0,69 |
| LUM | 4060 | 4,74 | 4,2 | 0,54 |
| C10orf10 | 11067 | 5,87 | 5,43 | 0,45 |
| KIAA1199 | 57214 | 5,46 | 5,08 | 0,37 |
| COL1A1 | 1277 | 7,07 | 6,52 | 0,55 |
| PVRL2 | 5819 | 7,52 | 6,88 | 0,64 |
| SMURF2 | 64750 | 7,89 | 7,28 | 0,61 |
| CCL13 | 6357 | 7,34 | 6,43 | 0,91 |
| NTAN1 | 123803 | 9,16 | 8,68 | 0,49 |
| TFEC | 22797 | 9,53 | 8,68 | 0,85 |
| GCNT1 | 2650 | 7,69 | 7,15 | 0,54 |
| PLD1 | 5337 | 6,63 | 5,95 | 0,67 |
| CD9 | 928 | 9,62 | 8,55 | 1,07 |
| EPB41L2 | 2037 | 9,04 | 8,38 | 0,66 |
| EIF4A3 | 9775 | 10,54 | 10,09 | 0,44 |

| | | | | |
|---|---|---|---|---|
| CCRL2 | 9034 | 8,92 | 8,17 | 0,75 |
| ARHGAP10 | 79658 | 7,96 | 7,5 | 0,46 |
| PDGFRL | 5157 | 4,99 | 4,59 | 0,4 |
| EPB41L3 | 23136 | 10,43 | 9,8 | 0,64 |
| CD24 | 100133941 | 6,34 | 5,38 | 0,96 |
| FAM168A | 23201 | 7,61 | 7,21 | 0,4 |
| KLF6 | 1316 | 9,57 | 8,89 | 0,67 |
| DSE | 29940 | 11,12 | 10,48 | 0,65 |
| MELK | 9833 | 6,65 | 6,07 | 0,58 |
| CD68 | 968 | 10,56 | 10,16 | 0,4 |
| MDC1 | 9656 | 7,22 | 6,82 | 0,4 |
| CLN6 | 54982 | 7,98 | 7,48 | 0,5 |
| COL3A1 | 1281 | 5,74 | 5,32 | 0,42 |
| IBSP | 3381 | 5,37 | 4,95 | 0,41 |
| RYR1 | 6261 | 7,13 | 6,53 | 0,6 |
| TRPM2 | 7226 | 6,98 | 6,55 | 0,43 |
| ASIP | 434 | 5,23 | 4,84 | 0,39 |
| PDIA3 | 2923 | 10,24 | 9,89 | 0,35 |
| BHLHE41 | 79365 | 8,47 | 7,43 | 1,03 |
| CEP55 | 55165 | 6,05 | 5,34 | 0,72 |
| HIVEP3 | 59269 | 7,13 | 6,5 | 0,62 |
| PRDX4 | 10549 | 9,73 | 9,32 | 0,4 |
| STAT1 | 6772 | 10,14 | 9,5 | 0,64 |
| APOL1 | 8542 | 7,99 | 7,39 | 0,6 |
| ARHGEF11 | 9826 | 7,45 | 6,96 | 0,49 |
| TGM2 | 7052 | 9,77 | 8,9 | 0,87 |
| MT1E | 4493 | 9,64 | 8,97 | 0,67 |
| ARMCX1 | 51309 | 8,18 | 7,69 | 0,49 |
| LAMC1 | 3915 | 6,79 | 6,16 | 0,63 |
| CYB5A | 1528 | 9,61 | 9,19 | 0,42 |
| MGAT5 | 4249 | 9,45 | 9,06 | 0,39 |
| CDCP1 | 64866 | 8,31 | 7,59 | 0,72 |
| DENND4C | 55667 | 9,34 | 8,86 | 0,48 |
| TSPAN4 | 7106 | 8,42 | 7,8 | 0,62 |
| TLR7 | 51284 | 9,49 | 8,38 | 1,11 |
| MGLL | 11343 | 9,67 | 9,1 | 0,57 |
| GNG12 | 55970 | 6,91 | 6,35 | 0,56 |
| RPS27L | 51065 | 8,05 | 7,53 | 0,52 |
| RASGRP3 | 25780 | 7,95 | 7,05 | 0,9 |
| TNFAIP3 | 7128 | 10,77 | 9,97 | 0,8 |
| RBP4 | 5950 | 7,85 | 7,27 | 0,58 |
| COL1A2 | 1278 | 6,13 | 5,7 | 0,43 |
| SASH1 | 23328 | 9,01 | 8,35 | 0,66 |
| THBS3 | 7059 | 6,09 | 5,78 | 0,31 |

| | | | | |
|---|---|---|---|---|
| PLOD3 | 8985 | 9,36 | 9,03 | 0,33 |
| SGK1 | 6446 | 12,04 | 11,46 | 0,58 |
| SLC39A8 | 64116 | 8,06 | 7,01 | 1,05 |
| HMOX1 | 3162 | 11,21 | 10,54 | 0,68 |
| LIMK2 | 3985 | 7,4 | 6,87 | 0,53 |
| LY96 | 23643 | 10,77 | 10,36 | 0,4 |
| KIFC3 | 3801 | 6,08 | 5,71 | 0,36 |
| CD151 | 977 | 8,58 | 8,06 | 0,52 |
| RRBP1 | 6238 | 7,88 | 7,48 | 0,41 |
| MILR1 | 284021 | 8,3 | 7,76 | 0,54 |
| FER | 2241 | 6,19 | 5,8 | 0,39 |
| C3AR1 | 719 | 11,49 | 10,56 | 0,93 |
| B3GALNT1 | 8706 | 6,44 | 5,77 | 0,67 |
| LAP3 | 51056 | 11,47 | 11,01 | 0,46 |
| SPATA7 | 55812 | 6,01 | 5,62 | 0,39 |
| FAM114A1 | 92689 | 6,73 | 6,28 | 0,46 |
| ATRN | 8455 | 7,54 | 7,18 | 0,36 |
| CCL20 | 6364 | 9,16 | 7,78 | 1,38 |
| MT2A | 4502 | 12,07 | 11,5 | 0,58 |
| MUC1 | 4582 | 6,51 | 6,1 | 0,41 |
| GIMAP6 | 474344 | 8,04 | 7,24 | 0,8 |
| CCND1 | 595 | 6,73 | 6,03 | 0,7 |
| UAP1L1 | 91373 | 6,93 | 6,55 | 0,38 |
| GADD45B | 4616 | 9,53 | 8,87 | 0,66 |
| CBR1 | 873 | 8,19 | 7,83 | 0,36 |
| NUCB1 | 4924 | 8,87 | 8,33 | 0,55 |
| MT1G | 4495 | 10,37 | 9,63 | 0,75 |
| NLK | 51701 | 7,5 | 7,11 | 0,39 |
| IGFBP3 | 3486 | 6,52 | 5,93 | 0,59 |
| GADD45G | 10912 | 6,77 | 6,27 | 0,5 |
| VCAM1 | 7412 | 6,21 | 5,5 | 0,71 |
| PRDM1 | 639 | 8,73 | 7,91 | 0,83 |
| KCNJ5 | 3762 | 6,19 | 5,77 | 0,42 |
| ABL2 | 27 | 7,44 | 6,84 | 0,61 |
| STEAP3 | 55240 | 7,25 | 6,77 | 0,48 |
| CDH6 | 1004 | 5,62 | 5,24 | 0,38 |
| SREBF1 | 6720 | 7,44 | 7,11 | 0,33 |
| APOL2 | 23780 | 7,4 | 7,04 | 0,36 |
| FGFR1 | 2260 | 6,54 | 6,12 | 0,43 |
| TMEM180 | 79847 | 7,48 | 7,12 | 0,35 |

g adjusted p-
nes are green.

| adjusted p-value |
|---|
| 0,00003 |
| 0,00003 |
| 0,00006 |
| 0,00006 |
| 0,00008 |
| 0,00008 |
| 0,00012 |
| 0,00012 |
| 0,00019 |
| 0,00019 |
| 0,00019 |
| 0,00027 |
| 0,00031 |
| 0,00031 |
| 0,00031 |
| 0,00031 |
| 0,00034 |
| 0,00034 |
| 0,00034 |
| 0,00036 |
| 0,00036 |
| 0,0004 |
| 0,0004 |
| 0,00042 |
| 0,00042 |
| 0,00042 |
| 0,00044 |
| 0,00047 |
| 0,00047 |
| 0,0005 |
| 0,00051 |
| 0,00052 |
| 0,00057 |
| 0,00059 |
| 0,00059 |
| 0,00059 |
| 0,00065 |
| 0,00068 |
| 0,00068 |

| |
|---|
| 0,00068 |
| 0,00068 |
| 0,0007 |
| 0,00076 |
| 0,00077 |
| 0,00081 |
| 0,00081 |
| 0,00083 |
| 0,00083 |
| 0,00093 |
| 0,00099 |
| 0,00099 |
| 0,00109 |
| 0,00109 |
| 0,00109 |
| 0,00109 |
| 0,00111 |
| 0,00112 |
| 0,00112 |
| 0,00112 |
| 0,00112 |
| 0,00115 |
| 0,00119 |
| 0,00119 |
| 0,00119 |
| 0,00127 |
| 0,00135 |
| 0,00135 |
| 0,00149 |
| 0,00149 |
| 0,00149 |
| 0,0015 |
| 0,00165 |
| 0,00165 |
| 0,00193 |
| 0,00193 |
| 0,00193 |
| 0,00193 |
| 0,00199 |
| 0,00199 |
| 0,00206 |
| 0,00207 |
| 0,00207 |
| 0,00207 |

| |
|---|
| 0,00207 |
| 0,00207 |
| 0,00207 |
| 0,00207 |
| 0,00207 |
| 0,00207 |
| 0,00223 |
| 0,00228 |
| 0,00228 |
| 0,00244 |
| 0,00246 |
| 0,0025 |
| 0,0025 |
| 0,00257 |
| 0,0026 |
| 0,00262 |
| 0,00263 |
| 0,00267 |
| 0,00267 |
| 0,00267 |
| 0,00267 |
| 0,0028 |
| 0,00285 |
| 0,00288 |
| 0,00288 |
| 0,00288 |
| 0,0029 |
| 0,0033 |
| 0,00346 |
| 0,00348 |
| 0,00355 |
| 0,00369 |
| 0,00369 |
| 0,00375 |
| 0,00385 |
| 0,00389 |
| 0,00389 |
| 0,00395 |
| 0,004 |
| 0,00412 |
| 0,00412 |
| 0,00421 |
| 0,00432 |
| 0,00433 |

| |
|---|
| 0,00459 |
| 0,00459 |
| 0,00459 |
| 0,00459 |
| 0,00459 |
| 0,00459 |
| 0,00459 |
| 0,00463 |
| 0,00463 |
| 0,00465 |
| 0,00475 |
| 0,0049 |
| 0,00491 |
| 0,00491 |
| 0,00491 |
| 0,00504 |
| 0,00504 |
| 0,0054 |
| 0,00547 |
| 0,00561 |
| 0,00574 |
| 0,00575 |
| 0,00582 |
| 0,00582 |
| 0,00582 |
| 0,00605 |
| 0,0066 |
| 0,00663 |
| 0,00668 |
| 0,00675 |
| 0,00683 |
| 0,00683 |
| 0,00683 |
| 0,00687 |
| 0,00698 |
| 0,00711 |
| 0,00723 |
| 0,00754 |
| 0,0076 |
| 0,0076 |
| 0,0076 |
| 0,0076 |
| 0,0076 |
| 0,00797 |

| |
|---|
| 0,00807 |
| 0,00812 |
| 0,00812 |
| 0,00817 |
| 0,00818 |
| 0,00818 |
| 0,00827 |
| 0,00856 |
| 0,00876 |
| 0,00877 |
| 0,00969 |
| 0,00992 |
| 0,00992 |
| 0,00992 |
| 0,0101 |
| 0,01016 |
| 0,01024 |
| 0,01035 |
| 0,01036 |
| 0,01036 |
| 0,01039 |
| 0,0104 |
| 0,0104 |
| 0,01079 |
| 0,01099 |
| 0,01115 |
| 0,01139 |
| 0,01139 |
| 0,01139 |
| 0,01143 |
| 0,0118 |
| 0,01183 |
| 0,01183 |
| 0,01183 |
| 0,01194 |
| 0,012 |
| 0,01204 |
| 0,01226 |
| 0,0123 |
| 0,01241 |
| 0,01252 |
| 0,0127 |
| 0,01279 |
| 0,01288 |

| |
|---|
| 0,01288 |
| 0,01295 |
| 0,01313 |
| 0,01316 |
| 0,0132 |
| 0,0132 |
| 0,01333 |
| 0,01348 |
| 0,01351 |
| 0,01352 |
| 0,01353 |
| 0,01356 |
| 0,01363 |
| 0,01373 |
| 0,01383 |
| 0,01402 |
| 0,01404 |
| 0,01425 |
| 0,01452 |
| 0,01541 |
| 0,01549 |
| 0,01552 |
| 0,01595 |
| 0,01618 |
| 0,01647 |
| 0,01656 |
| 0,01656 |
| 0,01656 |
| 0,01704 |
| 0,01704 |
| 0,01737 |
| 0,01763 |
| 0,01765 |
| 0,01765 |
| 0,01765 |
| 0,01765 |
| 0,01779 |
| 0,01779 |
| 0,0179 |
| 0,0179 |
| 0,01799 |
| 0,01852 |
| 0,01852 |
| 0,01859 |

| |
|---|
| 0,01883 |
| 0,01883 |
| 0,01883 |
| 0,01894 |
| 0,01896 |
| 0,0196 |
| 0,0198 |
| 0,01998 |
| 0,02036 |
| 0,02045 |
| 0,02045 |
| 0,02048 |
| 0,02048 |
| 0,02081 |
| 0,02128 |
| 0,02128 |
| 0,02155 |
| 0,02162 |
| 0,02167 |
| 0,02167 |
| 0,02167 |
| 0,0217 |
| 0,02187 |
| 0,02206 |
| 0,02257 |
| 0,02272 |
| 0,02272 |
| 0,02272 |
| 0,02296 |
| 0,02296 |
| 0,02311 |
| 0,02311 |
| 0,02311 |
| 0,02312 |
| 0,02329 |
| 0,0235 |
| 0,02353 |
| 0,02353 |
| 0,02378 |
| 0,02387 |
| 0,02387 |
| 0,02393 |
| 0,02409 |
| 0,02411 |

| |
|---|
| 0,02411 |
| 0,02412 |
| 0,02418 |
| 0,02487 |
| 0,0252 |
| 0,02551 |
| 0,02551 |
| 0,02551 |
| 0,02585 |
| 0,02606 |
| 0,02606 |
| 0,02606 |
| 0,02606 |
| 0,02606 |
| 0,02606 |
| 0,02618 |
| 0,02626 |
| 0,0265 |
| 0,02654 |
| 0,02736 |
| 0,02736 |
| 0,02736 |
| 0,02772 |
| 0,02806 |
| 0,02842 |
| 0,02854 |
| 0,02854 |
| 0,02854 |
| 0,02866 |
| 0,02887 |
| 0,02912 |
| 0,02912 |
| 0,02912 |
| 0,02928 |
| 0,02929 |
| 0,02943 |
| 0,02989 |
| 0,02989 |
| 0,03004 |
| 0,03059 |
| 0,03059 |
| 0,03123 |
| 0,03123 |
| 0,03123 |

| |
|---|
| 0,03178 |
| 0,03203 |
| 0,033 |
| 0,03305 |
| 0,0332 |
| 0,03354 |
| 0,03429 |
| 0,03434 |
| 0,03434 |
| 0,03436 |
| 0,03443 |
| 0,03482 |
| 0,03482 |
| 0,03482 |
| 0,03497 |
| 0,03592 |
| 0,03598 |
| 0,03652 |
| 0,03661 |
| 0,03673 |
| 0,03716 |
| 0,03728 |
| 0,03733 |
| 0,03738 |
| 0,03756 |
| 0,03801 |
| 0,03801 |
| 0,03801 |
| 0,03811 |
| 0,03819 |
| 0,03838 |
| 0,03845 |
| 0,03845 |
| 0,0385 |
| 0,03866 |
| 0,03893 |
| 0,03893 |
| 0,03905 |
| 0,03919 |
| 0,03919 |
| 0,03919 |
| 0,03919 |
| 0,03928 |
| 0,03936 |

| |
|---|
| 0,03973 |
| 0,04045 |
| 0,04047 |
| 0,04047 |
| 0,04047 |
| 0,04047 |
| 0,04115 |
| 0,04142 |
| 0,04143 |
| 0,04155 |
| 0,04155 |
| 0,04161 |
| 0,04202 |
| 0,04242 |
| 0,04244 |
| 0,0434 |
| 0,0437 |
| 0,04372 |
| 0,04399 |
| 0,04402 |
| 0,04425 |
| 0,04425 |
| 0,04426 |
| 0,04447 |
| 0,04447 |
| 0,04452 |
| 0,04487 |
| 0,04579 |
| 0,04714 |
| 0,04714 |
| 0,04716 |
| 0,04729 |
| 0,04753 |
| 0,04865 |
| 0,04884 |
| 0,04884 |
| 0,04892 |
| 0,04933 |
| 0,04939 |
| 0,0495 |
