## Supplementary material for "A mosaic renal myeloid subtype with T-cell inhibitory and protumoral features is linked to immune escape and survival in clear cell renal cell cancer": Brech et al_supplementary Table S9

**Table S9**: Downregulated ercDC_ccRCC DEGs. Listed are genes significantly (adjusted p<0.05) downregulated betw ercDC_ccRCC&infMΦ_ascOvCa and control group (357 of 788 ercDC_ccRCC DEGs). Genes are sorted acc. to inc adjusted p-values. Genes that belong to ercDC_ccRCC marker genes are in bold green. Genes related to ercDC_ccRC genes are in green. Genes mentioned in main text are in grey

| **Gene symbol** | **Entrez ID** | **norm. expr. values (log) ercDC_RCC&infMΦ_ascOvCa** | **norm. expr.values (log) control group** | **logFC** |
|---|---|---|---|---|
| **FAM65B** | 9750 | 5,16 | 6,4 | -1,23 |
| RPS9 | 6203 | 11,06 | 11,73 | -0,67 |
| SLC25A6 | 293 | 10,86 | 11,57 | -0,71 |
| **FGR** | 2268 | 9,39 | 10,53 | -1,13 |
| NUP210 | 23225 | 6,66 | 7,48 | -0,81 |
| RHOF | 54509 | 5,82 | 6,94 | -1,12 |
| PBX2 | 5089 | 7,06 | 7,64 | -0,58 |
| CCND3 | 896 | 8,81 | 9,53 | -0,71 |
| **CFP** | 5199 | 6,44 | 7,77 | -1,34 |
| ICAM3 | 3385 | 6,56 | 7,76 | -1,2 |
| EIF4EBP2 | 1979 | 7,83 | 8,42 | -0,59 |
| **FYN** | 2534 | 6,14 | 7,22 | -1,07 |
| CD244 | 51744 | 5,14 | 6,09 | -0,95 |
| RAB11FIP4 | 84440 | 5,73 | 6,37 | -0,65 |
| SMAD3 | 4088 | 5,91 | 6,56 | -0,65 |
| **CYTIP** | 9595 | 9,11 | 9,94 | -0,83 |
| EIF3E | 3646 | 8,19 | 8,71 | -0,51 |
| RPL15 | 6138 | 8,28 | 8,79 | -0,5 |
| LIMD2 | 80774 | 5,99 | 6,5 | -0,51 |
| ZBTB18 | 10472 | 6,57 | 7,21 | -0,65 |
| **EMR3** | 84658 | 5,51 | 6,98 | -1,47 |
| CSNK1G2 | 1455 | 7,07 | 7,59 | -0,52 |
| GSTP1 | 2950 | 9,74 | 10,3 | -0,56 |
| BCL11A | 53335 | 5,33 | 6,54 | -1,21 |
| FCHO1 | 23149 | 6,1 | 6,62 | -0,52 |
| ATP8A1 | 10396 | 5,96 | 6,76 | -0,8 |
| RPL31 | 6160 | 8 | 8,68 | -0,68 |
| UPK3A | 7380 | 5,03 | 5,59 | -0,56 |
| ITGA4 | 3676 | 5,84 | 6,78 | -0,94 |
| GLTSCR2 | 29997 | 8,11 | 8,66 | -0,55 |
| LSP1 | 4046 | 7,96 | 8,87 | -0,91 |
| SEZ6L | 23544 | 4,82 | 5,4 | -0,58 |
| TSPAN32 | 10077 | 5,87 | 6,66 | -0,78 |
| EIF3G | 8666 | 9,65 | 10,18 | -0,53 |
| KIAA0922 | 23240 | 7,48 | 8,29 | -0,81 |
| JAK1 | 3716 | 8,75 | 9,34 | -0,59 |
| **NEDD9** | 4739 | 6,77 | 7,65 | -0,88 |
| ACAA1 | 30 | 8,38 | 8,86 | -0,48 |

| | | | | |
|---|---|---|---|---|
| P2RY10 | 27334 | 4,45 | 5,22 | -0,77 |
| SIGIRR | 59307 | 6,23 | 6,73 | -0,51 |
| **TES** | 26136 | 8 | 8,75 | -0,75 |
| RPS23 | 6228 | 7,21 | 7,77 | -0,56 |
| RPS16 | 6217 | 9,18 | 9,64 | -0,46 |
| MED28 | 80306 | 7,76 | 8,26 | -0,49 |
| **CD48** | 962 | 8,97 | 9,82 | -0,85 |
| RAC2 | 5880 | 8,37 | 9,26 | -0,88 |
| CD101 | 9398 | 6,47 | 7,63 | -1,16 |
| SF3A1 | 10291 | 8,16 | 8,68 | -0,53 |
| USP3 | 9960 | 9,49 | 10,02 | -0,53 |
| SPN | 6693 | 6,63 | 7,43 | -0,81 |
| S1PR4 | 8698 | 6,86 | 7,41 | -0,55 |
| **LYST** | 1130 | 7,27 | 8,34 | -1,07 |
| RAB11FIP1 | 80223 | 7,96 | 8,58 | -0,62 |
| CNN2 | 1265 | 8,35 | 8,93 | -0,58 |
| POLR1D | 51082 | 9,04 | 9,51 | -0,47 |
| C15orf39 | 56905 | 6,58 | 7,21 | -0,63 |
| TRAF3IP3 | 80342 | 5,66 | 6,48 | -0,83 |
| MST4 | 51765 | 6,54 | 7,31 | -0,77 |
| RNF41 | 10193 | 6,25 | 6,73 | -0,48 |
| FCER1A | 2205 | 4,97 | 6,36 | -1,39 |
| RNF24 | 11237 | 6,52 | 7,18 | -0,66 |
| WAC | 51322 | 8,02 | 8,49 | -0,47 |
| BID | 637 | 8,81 | 9,36 | -0,56 |
| FLT3 | 2322 | 5,43 | 6,36 | -0,93 |
| MTMR14 | 64419 | 7,39 | 7,85 | -0,46 |
| PRKCE | 5581 | 5,48 | 5,98 | -0,49 |
| ZNF652 | 22834 | 5,99 | 6,85 | -0,85 |
| CNOT8 | 9337 | 8,02 | 8,62 | -0,6 |
| PTP4A2 | 8073 | 9,97 | 10,51 | -0,54 |
| FLOT2 | 2319 | 7,64 | 8,29 | -0,65 |
| CD11C | 3687 | 9,01 | 9,9 | -0,89 |
| NACA | 4666 | 8,7 | 9,18 | -0,48 |
| APOBR | 55911 | 7,27 | 7,91 | -0,63 |
| CA5B | 11238 | 5,6 | 6,09 | -0,49 |
| **CD52** | 1043 | 10,72 | 11,48 | -0,76 |
| KLF12 | 11278 | 5,48 | 6,26 | -0,78 |
| FAM117A | 81558 | 6,35 | 6,91 | -0,56 |
| INSR | 3643 | 6,55 | 7,2 | -0,65 |
| NLRP1 | 22861 | 5,88 | 6,62 | -0,74 |
| CS | 1431 | 9,84 | 10,19 | -0,35 |
| PELI2 | 57161 | 6,19 | 6,87 | -0,68 |
| APAF1 | 317 | 7,06 | 7,65 | -0,6 |

| | | | | |
|---|---|---|---|---|
| MAP4K1 | 11184 | 6,29 | 7,04 | -0,76 |
| GALNT3 | 2591 | 4,82 | 5,39 | -0,57 |
| RASSF2 | 9770 | 7,71 | 8,46 | -0,75 |
| EIF4B | 1975 | 7,63 | 8,4 | -0,77 |
| CPPED1 | 55313 | 7,23 | 8,05 | -0,82 |
| **CCDC88C** | 440193 | 4,64 | 5,48 | -0,84 |
| CD1C | 911 | 6,2 | 7,52 | -1,31 |
| PLP2 | 5355 | 9,74 | 10,44 | -0,7 |
| PPM1F | 9647 | 7,17 | 7,63 | -0,46 |
| VPS51 | 738 | 8,27 | 8,71 | -0,44 |
| RBL2 | 5934 | 6,79 | 7,34 | -0,55 |
| IL16 | 3603 | 6,91 | 7,34 | -0,44 |
| EIF3M | 10480 | 9,05 | 9,61 | -0,56 |
| ITPK1 | 3705 | 8,55 | 8,98 | -0,43 |
| EIF3H | 8667 | 7,3 | 7,86 | -0,55 |
| MEFV | 4210 | 4,72 | 5,35 | -0,63 |
| PRKCB | 5579 | 6,61 | 7,54 | -0,93 |
| ZBTB11 | 27107 | 7,77 | 8,24 | -0,47 |
| UNC119 | 9094 | 7,54 | 7,98 | -0,44 |
| PGLS | 25796 | 8,03 | 8,59 | -0,55 |
| STK17B | 9262 | 7,68 | 8,45 | -0,78 |
| ZDHHC18 | 84243 | 7,58 | 8,08 | -0,5 |
| OXA1L | 5018 | 9,68 | 10,07 | -0,39 |
| GSE1 | 23199 | 7,04 | 7,61 | -0,57 |
| DNAJC4 | 3338 | 6,99 | 7,51 | -0,53 |
| TKT | 7086 | 9,12 | 9,79 | -0,67 |
| GMFG | 9535 | 10,23 | 10,68 | -0,46 |
| RPL36 | 25873 | 9,89 | 10,32 | -0,43 |
| TOB1 | 10140 | 8,36 | 9,23 | -0,87 |
| RPS27 | 6232 | 8,08 | 8,48 | -0,41 |
| NIN | 51199 | 7,34 | 7,81 | -0,46 |
| IMPA2 | 3613 | 6,92 | 7,57 | -0,64 |
| **CDC42EP3** | 10602 | 7,41 | 8,31 | -0,9 |
| CDKN1B | 1027 | 9,32 | 9,83 | -0,51 |
| CAT | 847 | 8,61 | 9,31 | -0,7 |
| ESYT1 | 23344 | 9,09 | 9,58 | -0,48 |
| FDFT1 | 2222 | 8,65 | 9,2 | -0,55 |
| DGKE | 8526 | 4,68 | 5,18 | -0,5 |
| FBL | 2091 | 9,28 | 9,73 | -0,45 |
| AGTPBP1 | 23287 | 7,8 | 8,33 | -0,53 |
| ATP2A3 | 489 | 5,71 | 6,32 | -0,61 |
| CORO1A | 11151 | 8,09 | 8,87 | -0,78 |
| Sep 09 | 10801 | 6 | 6,57 | -0,57 |
| SIK3 | 23387 | 6,15 | 6,75 | -0,59 |

| | | | | |
|---|---|---|---|---|
| KAT6A | 7994 | 6,87 | 7,28 | -0,41 |
| RSL1D1 | 26156 | 7,61 | 8,09 | -0,48 |
| C1RL | 51279 | 6,47 | 7,17 | -0,69 |
| GABBR1 | 2550 | 5,84 | 6,38 | -0,54 |
| **CAPN2** | 824 | 9,86 | 10,6 | -0,74 |
| XYLT1 | 64131 | 6,58 | 7,39 | -0,82 |
| NDUFA10 | 4705 | 8,51 | 8,94 | -0,43 |
| MPHOSPH9 | 10198 | 6,25 | 6,74 | -0,5 |
| NDST1 | 3340 | 6,13 | 6,6 | -0,47 |
| C20orf27 | 54976 | 6,14 | 6,75 | -0,61 |
| PTPN6 | 5777 | 9,75 | 10,27 | -0,52 |
| NAP1L1 | 4673 | 9,09 | 9,66 | -0,57 |
| PSIP1 | 11168 | 6,76 | 7,39 | -0,63 |
| MAN2A2 | 4122 | 6,98 | 7,41 | -0,43 |
| ACAP1 | 9744 | 5,72 | 6,32 | -0,6 |
| ZNF467 | 168544 | 5,78 | 6,22 | -0,43 |
| SLC9A3R1 | 9368 | 7,68 | 8,15 | -0,46 |
| CERK | 64781 | 7,72 | 8,18 | -0,46 |
| PDE4A | 5141 | 6,02 | 6,65 | -0,63 |
| RBM3 | 5935 | 6,88 | 7,39 | -0,51 |
| RFX7 | 64864 | 7,23 | 7,67 | -0,44 |
| ELF4 | 2000 | 8,01 | 8,46 | -0,45 |
| CDKN2D | 1032 | 5,96 | 6,42 | -0,46 |
| ROGDI | 79641 | 7 | 7,55 | -0,55 |
| **FLNA** | 2316 | 8,08 | 8,82 | -0,74 |
| WDR48 | 57599 | 7,3 | 7,76 | -0,47 |
| FAM60A | 58516 | 6,69 | 7,26 | -0,56 |
| CASP1 | 834 | 8,6 | 9,42 | -0,83 |
| AIP | 9049 | 7,26 | 7,69 | -0,43 |
| TESC | 54997 | 5,29 | 6,02 | -0,73 |
| LCP1 | 3936 | 11,4 | 11,85 | -0,45 |
| CYLD | 1540 | 8,07 | 8,54 | -0,47 |
| RARA | 5914 | 6,19 | 6,9 | -0,71 |
| NEK9 | 91754 | 7,55 | 7,92 | -0,38 |
| VCL | 7414 | 8,14 | 8,85 | -0,71 |
| ABCA7 | 10347 | 6,27 | 6,69 | -0,42 |
| IL18R1 | 8809 | 4,28 | 4,92 | -0,64 |
| ADRBK2 | 157 | 8,19 | 8,64 | -0,46 |
| RUNX3 | 864 | 7,46 | 8,11 | -0,65 |
| CCNG1 | 900 | 9 | 9,48 | -0,47 |
| SCRN1 | 9805 | 7,57 | 8,2 | -0,62 |
| AHCYL2 | 23382 | 6,29 | 6,7 | -0,41 |
| PHF17 | 79960 | 5,83 | 6,32 | -0,48 |
| SETD2 | 29072 | 6,14 | 6,55 | -0,41 |

| | | | | |
|---|---|---|---|---|
| PCYOX1L | 78991 | 7,21 | 7,7 | -0,48 |
| AES | 166 | 7,76 | 8,22 | -0,46 |
| **PLAC8** | 51316 | 7,56 | 8,84 | -1,28 |
| SULT1B1 | 27284 | 5,2 | 5,7 | -0,5 |
| RIN3 | 79890 | 7,1 | 7,8 | -0,7 |
| RPS21 | 6227 | 8,66 | 9,04 | -0,37 |
| RAP1GAP2 | 23108 | 6,44 | 7,05 | -0,61 |
| DHPS | 1725 | 7,38 | 7,8 | -0,42 |
| PITPNM1 | 9600 | 6,47 | 6,91 | -0,44 |
| VRK1 | 7443 | 7,18 | 7,63 | -0,45 |
| ADAP1 | 11033 | 7,48 | 7,96 | -0,48 |
| ATP11B | 23200 | 6,42 | 6,91 | -0,49 |
| TUBA4A | 7277 | 7,17 | 7,76 | -0,58 |
| WDR82 | 80335 | 9,5 | 9,87 | -0,37 |
| FRAT2 | 23401 | 7,63 | 8,08 | -0,45 |
| ZFAND1 | 79752 | 7,27 | 7,75 | -0,48 |
| LRMP | 4033 | 7,08 | 7,76 | -0,68 |
| CPSF6 | 11052 | 6,52 | 6,97 | -0,45 |
| CD207 | 50489 | 4,71 | 5,56 | -0,85 |
| C11orf21 | 29125 | 5,58 | 6,4 | -0,82 |
| SRRM1 | 10250 | 8,61 | 8,93 | -0,32 |
| BRD3 | 8019 | 7,75 | 8,11 | -0,36 |
| SERTAD2 | 9792 | 9,21 | 9,6 | -0,38 |
| STAG3L4 | 64940 | 6,24 | 6,69 | -0,44 |
| DHTKD1 | 55526 | 6,23 | 6,7 | -0,47 |
| NONO | 4841 | 8,29 | 8,69 | -0,4 |
| TBL1X | 6907 | 6,31 | 7,09 | -0,78 |
| EIF2S3 | 1968 | 7,6 | 8,33 | -0,72 |
| LRBA | 987 | 6,61 | 7,11 | -0,5 |
| VAMP2 | 6844 | 6,94 | 7,38 | -0,43 |
| MKNK2 | 2872 | 8,6 | 9,08 | -0,48 |
| STAG2 | 10735 | 7,95 | 8,44 | -0,48 |
| FMNL1 | 752 | 7,5 | 8,02 | -0,53 |
| ELF2 | 1998 | 6,53 | 6,98 | -0,44 |
| SETBP1 | 26040 | 5,3 | 5,74 | -0,45 |
| MYO1F | 4542 | 9,27 | 9,75 | -0,48 |
| SPATA6 | 54558 | 5,31 | 5,84 | -0,54 |
| DAPP1 | 27071 | 7,76 | 8,38 | -0,63 |
| IDH3A | 3419 | 8,05 | 8,51 | -0,46 |
| TMEM104 | 54868 | 6,98 | 7,35 | -0,37 |
| MBNL3 | 55796 | 7,27 | 7,75 | -0,48 |
| CCNI | 10983 | 8,1 | 8,64 | -0,54 |
| ZNF318 | 24149 | 6,89 | 7,28 | -0,39 |
| RASGRP2 | 10235 | 5,83 | 6,4 | -0,57 |

| | | | | |
|---|---|---|---|---|
| PYCARD | 29108 | 9,79 | 10,25 | -0,46 |
| BACH2 | 60468 | 4,11 | 4,46 | -0,35 |
| STAT5B | 6777 | 6,81 | 7,19 | -0,38 |
| C6orf48 | 50854 | 8,36 | 8,91 | -0,55 |
| ARF5 | 381 | 7,99 | 8,36 | -0,37 |
| RPS27A | 6233 | 8,58 | 8,97 | -0,39 |
| MEF2D | 4209 | 6,9 | 7,3 | -0,4 |
| MEX3C | 51320 | 6,84 | 7,35 | -0,52 |
| CELF2 | 10659 | 8,11 | 8,69 | -0,57 |
| GDI2 | 2665 | 9,76 | 10,23 | -0,47 |
| OSBPL8 | 114882 | 7,37 | 8,09 | -0,72 |
| GID8 | 54994 | 8,29 | 8,65 | -0,36 |
| MAP2K3 | 5606 | 7,4 | 7,97 | -0,57 |
| IPCEF1 | 26034 | 5,85 | 6,52 | -0,66 |
| ARAF | 369 | 7,87 | 8,23 | -0,36 |
| CCDC69 | 26112 | 6,21 | 6,83 | -0,61 |
| FRY | 10129 | 6,65 | 7,45 | -0,81 |
| SNRK | 54861 | 8,11 | 8,56 | -0,46 |
| SVIL | 6840 | 6,72 | 7,43 | -0,71 |
| SPINT1 | 6692 | 7,23 | 7,74 | -0,51 |
| AFF3 | 3899 | 4,84 | 5,46 | -0,62 |
| ARHGEF6 | 9459 | 9,37 | 9,79 | -0,42 |
| CREBBP | 1387 | 7,33 | 7,67 | -0,34 |
| VEZF1 | 7716 | 8,31 | 8,75 | -0,44 |
| DIAPH1 | 1729 | 8,1 | 8,56 | -0,46 |
| CHST2 | 9435 | 6,02 | 6,57 | -0,55 |
| CDC40 | 51362 | 6,68 | 7,1 | -0,42 |
| RPL14 | 9045 | 8,91 | 9,39 | -0,48 |
| RMND5A | 64795 | 6,93 | 7,34 | -0,41 |
| CIDEB | 27141 | 6,96 | 7,42 | -0,46 |
| LIMD1 | 8994 | 6,89 | 7,33 | -0,44 |
| CHAF1A | 10036 | 5,81 | 6,14 | -0,33 |
| TP53 | 7157 | 6,67 | 7,15 | -0,48 |
| BAG1 | 573 | 7,88 | 8,28 | -0,4 |
| INPP5F | 22876 | 5,84 | 6,35 | -0,51 |
| TRAPPC6A | 79090 | 7,42 | 7,81 | -0,38 |
| ANP32A | 8125 | 8,95 | 9,35 | -0,4 |
| SLC12A6 | 9990 | 6,86 | 7,43 | -0,57 |
| PMS2P1 | 5379 | 6,89 | 7,3 | -0,41 |
| RNF126 | 55658 | 6,92 | 7,3 | -0,37 |
| BPTF | 2186 | 5,81 | 6,14 | -0,34 |
| SELL | 6402 | 6,73 | 7,49 | -0,76 |
| PNRC2 | 55629 | 8,69 | 9,05 | -0,36 |
| VIPR1 | 7433 | 5,58 | 6,08 | -0,5 |

| | | | | |
|---|---|---|---|---|
| TMEM8B | 51754 | 5,77 | 6,19 | -0,42 |
| SLC25A40 | 55972 | 8,48 | 8,96 | -0,48 |
| NADSYN1 | 55191 | 7,96 | 8,31 | -0,35 |
| PPP3CA | 5530 | 8,44 | 8,9 | -0,47 |
| NCF2 | 4688 | 10,88 | 11,37 | -0,48 |
| ZNF592 | 9640 | 7,31 | 7,65 | -0,34 |
| TMEM66 | 51669 | 7,54 | 7,91 | -0,37 |
| IRF4 | 3662 | 5,92 | 6,57 | -0,65 |
| LSM6 | 11157 | 8,82 | 9,2 | -0,38 |
| POLR1E | 64425 | 5,27 | 5,6 | -0,34 |
| URI1 | 8725 | 7,23 | 7,63 | -0,4 |
| TNKS2 | 80351 | 7,94 | 8,39 | -0,46 |
| STK24 | 8428 | 8,35 | 8,69 | -0,34 |
| LIG3 | 3980 | 5,93 | 6,26 | -0,33 |
| HIVEP1 | 3096 | 6,93 | 7,49 | -0,56 |
| FAM168B | 130074 | 7,59 | 8 | -0,41 |
| SLC38A1 | 81539 | 5,58 | 6,49 | -0,92 |
| SPIB | 6689 | 5,16 | 5,66 | -0,5 |
| SLCO3A1 | 28232 | 7,28 | 7,88 | -0,6 |
| CYFIP2 | 26999 | 6,5 | 7,21 | -0,7 |
| AMPD2 | 271 | 7,62 | 8,22 | -0,6 |
| DCLRE1C | 64421 | 6,68 | 7,08 | -0,4 |
| NFE2 | 4778 | 4,43 | 4,95 | -0,52 |
| CARS2 | 79587 | 8,43 | 8,78 | -0,35 |
| G3BP2 | 9908 | 8,25 | 8,62 | -0,37 |
| PPP6C | 5537 | 8,9 | 9,26 | -0,36 |
| SLC25A28 | 81894 | 7,5 | 7,91 | -0,41 |
| RSL24D1 | 51187 | 9,06 | 9,46 | -0,41 |
| MTMR1 | 8776 | 7,46 | 8 | -0,54 |
| RPS6KA3 | 6197 | 9 | 9,41 | -0,41 |
| NCOA1 | 8648 | 8,07 | 8,44 | -0,37 |
| EPB41 | 2035 | 6,1 | 6,49 | -0,4 |
| RPL23A | 6147 | 9,34 | 9,79 | -0,45 |
| Sep 06 | 23157 | 7,3 | 7,79 | -0,48 |
| LST1 | 7940 | 9,87 | 10,42 | -0,55 |
| PACS1 | 55690 | 6,94 | 7,27 | -0,33 |
| LNPEP | 4012 | 7,13 | 7,65 | -0,53 |
| RCOR1 | 23186 | 7,83 | 8,17 | -0,34 |
| OBFC1 | 79991 | 6,69 | 7,02 | -0,34 |
| CASP2 | 835 | 5,73 | 6,14 | -0,41 |
| USP48 | 84196 | 6,51 | 6,85 | -0,34 |
| IFT20 | 90410 | 8,71 | 9,13 | -0,42 |
| SCAF4 | 57466 | 6,47 | 6,9 | -0,43 |
| CBX7 | 23492 | 6,92 | 7,28 | -0,36 |

| | | | | |
|---|---|---|---|---|
| RGS14 | 10636 | 5,77 | 6,15 | -0,38 |
| NAAA | 27163 | 7,98 | 8,49 | -0,52 |
| CHP1 | 11261 | 7,62 | 8,04 | -0,42 |
| SLC25A38 | 54977 | 7,26 | 7,59 | -0,33 |
| MKL1 | 57591 | 7,13 | 7,5 | -0,37 |
| LSM7 | 51690 | 8,42 | 8,79 | -0,38 |
| EFHD2 | 79180 | 9,56 | 10,01 | -0,46 |
| ZBTB33 | 10009 | 8,84 | 9,18 | -0,34 |
| SH2D3C | 10044 | 5,5 | 6,01 | -0,51 |
| CD1E | 913 | 4,58 | 5,72 | -1,14 |
| C2orf49 | 79074 | 5,79 | 6,1 | -0,31 |
| PDE6G | 5148 | 6,41 | 6,77 | -0,36 |
| CRBN | 51185 | 8,36 | 8,75 | -0,4 |
| CLEC4A | 50856 | 7,78 | 8,56 | -0,77 |
| **IRAK3** | 11213 | 7,18 | 8,02 | -0,84 |
| NOTCH2 | 4853 | 8,19 | 8,73 | -0,54 |
| KCTD15 | 79047 | 4,8 | 5,28 | -0,47 |
| SMARCC1 | 6599 | 7,39 | 7,75 | -0,35 |
| BCL2L13 | 23786 | 6,48 | 6,82 | -0,35 |
| KDM4B | 23030 | 7,02 | 7,38 | -0,36 |
| SLC25A12 | 8604 | 6,28 | 6,62 | -0,34 |
| ERLIN2 | 11160 | 6,42 | 6,85 | -0,44 |
| KLF13 | 51621 | 7,21 | 7,74 | -0,54 |
| CD37 | 951 | 9,15 | 9,74 | -0,59 |
| U2SURP | 23350 | 6,31 | 6,79 | -0,48 |
| MAML3 | 55534 | 5,91 | 6,43 | -0,52 |
| ZNF281 | 23528 | 7,48 | 7,86 | -0,39 |
| RFTN1 | 23180 | 8,65 | 9,08 | -0,43 |
| C21orf91 | 54149 | 6,51 | 6,98 | -0,47 |
| ANAPC15 | 25906 | 7,64 | 7,99 | -0,35 |
| UQCRB | 7381 | 6,38 | 6,75 | -0,37 |
| PLCB2 | 5330 | 8,01 | 8,41 | -0,4 |
| AKIRIN1 | 79647 | 7,91 | 8,35 | -0,44 |
| STK4 | 6789 | 7,23 | 7,79 | -0,56 |
| METTL9 | 51108 | 8,88 | 9,4 | -0,52 |
| PHF20 | 51230 | 7,39 | 7,84 | -0,45 |
| IPO5 | 3843 | 7,6 | 8,06 | -0,46 |
| CBX4 | 8535 | 7,15 | 7,56 | -0,41 |
| RSBN1 | 54665 | 7,42 | 7,85 | -0,43 |
| SLC1A5 | 6510 | 7,39 | 7,8 | -0,4 |
| LANCL1 | 10314 | 7,85 | 8,21 | -0,36 |
| ATPAF2 | 91647 | 6,47 | 6,81 | -0,33 |
| HNRNPUL1 | 11100 | 7,95 | 8,31 | -0,36 |
| ZMIZ1 | 57178 | 8,79 | 9,11 | -0,32 |

| | | | | |
|---|---|---|---|---|
| SCN9A | 6335 | 3,97 | 4,4 | -0,44 |
| OGT | 8473 | 6,57 | 7,18 | -0,61 |
| KLHDC2 | 23588 | 8,3 | 8,68 | -0,39 |
| TMPO | 7112 | 7,13 | 7,65 | -0,52 |
| EVI2B | 2124 | 11,04 | 11,39 | -0,36 |
| KDM5A | 5927 | 6,64 | 7,03 | -0,39 |
| UQCRC2 | 7385 | 6,85 | 7,21 | -0,36 |
| FXYD5 | 53827 | 9,88 | 10,31 | -0,43 |
| CBX6 | 23466 | 7,35 | 7,72 | -0,38 |
| DDX46 | 9879 | 7,73 | 8,06 | -0,33 |
| ADRBK1 | 156 | 7,28 | 7,7 | -0,42 |

ween
reasing
[illegible]C marker

| adjusted p-value |
|---|
| 0,00034 |
| 0,0004 |
| 0,00043 |
| 0,00059 |
| 0,00068 |
| 0,00068 |
| 0,0007 |
| 0,0007 |
| 0,00081 |
| 0,00093 |
| 0,00104 |
| 0,00109 |
| 0,00112 |
| 0,00119 |
| 0,00119 |
| 0,00126 |
| 0,00149 |
| 0,00166 |
| 0,00166 |
| 0,0017 |
| 0,0017 |
| 0,00183 |
| 0,00196 |
| 0,00197 |
| 0,00199 |
| 0,00199 |
| 0,00207 |
| 0,00223 |
| 0,00223 |
| 0,00224 |
| 0,00228 |
| 0,0025 |
| 0,00254 |
| 0,00267 |
| 0,00288 |
| 0,00289 |
| 0,00298 |
| 0,00316 |

| |
|---|
| 0,00319 |
| 0,00327 |
| 0,0033 |
| 0,00366 |
| 0,00375 |
| 0,00375 |
| 0,00375 |
| 0,00375 |
| 0,00375 |
| 0,00392 |
| 0,00421 |
| 0,00424 |
| 0,00434 |
| 0,00459 |
| 0,00463 |
| 0,0047 |
| 0,00475 |
| 0,00475 |
| 0,00504 |
| 0,00545 |
| 0,00547 |
| 0,00575 |
| 0,00582 |
| 0,0059 |
| 0,006 |
| 0,00616 |
| 0,00625 |
| 0,00625 |
| 0,00625 |
| 0,00637 |
| 0,0066 |
| 0,00663 |
| 0,00663 |
| 0,00663 |
| 0,00681 |
| 0,00683 |
| 0,00683 |
| 0,00683 |
| 0,00687 |
| 0,00687 |
| 0,00693 |
| 0,00711 |
| 0,00739 |
| 0,00741 |

| |
|---|
| 0,00741 |
| 0,00741 |
| 0,0076 |
| 0,0076 |
| 0,0076 |
| 0,0076 |
| 0,0076 |
| 0,00777 |
| 0,00781 |
| 0,00818 |
| 0,00818 |
| 0,00912 |
| 0,00925 |
| 0,00926 |
| 0,00926 |
| 0,00931 |
| 0,00949 |
| 0,00966 |
| 0,00975 |
| 0,00975 |
| 0,00975 |
| 0,00991 |
| 0,00992 |
| 0,01 |
| 0,01007 |
| 0,01007 |
| 0,01017 |
| 0,01024 |
| 0,01035 |
| 0,0104 |
| 0,0104 |
| 0,0104 |
| 0,0104 |
| 0,01092 |
| 0,01099 |
| 0,01101 |
| 0,01123 |
| 0,01138 |
| 0,01139 |
| 0,01162 |
| 0,0118 |
| 0,01185 |
| 0,01194 |
| 0,01194 |

| |
|---|
| 0,01196 |
| 0,012 |
| 0,012 |
| 0,012 |
| 0,01213 |
| 0,01219 |
| 0,01226 |
| 0,01241 |
| 0,0126 |
| 0,0132 |
| 0,01332 |
| 0,01333 |
| 0,01348 |
| 0,01348 |
| 0,01348 |
| 0,01351 |
| 0,01354 |
| 0,01361 |
| 0,01363 |
| 0,01376 |
| 0,01383 |
| 0,01404 |
| 0,01414 |
| 0,01425 |
| 0,01475 |
| 0,01514 |
| 0,01514 |
| 0,01539 |
| 0,01541 |
| 0,01545 |
| 0,01549 |
| 0,01549 |
| 0,01549 |
| 0,01551 |
| 0,01552 |
| 0,0158 |
| 0,01618 |
| 0,01621 |
| 0,01656 |
| 0,01661 |
| 0,01661 |
| 0,0167 |
| 0,01672 |
| 0,01732 |

| |
|---|
| 0,01765 |
| 0,01788 |
| 0,01792 |
| 0,01799 |
| 0,01799 |
| 0,01818 |
| 0,01826 |
| 0,01832 |
| 0,01837 |
| 0,0186 |
| 0,0187 |
| 0,01883 |
| 0,01891 |
| 0,01893 |
| 0,01898 |
| 0,01902 |
| 0,01971 |
| 0,01987 |
| 0,01987 |
| 0,01998 |
| 0,02036 |
| 0,02036 |
| 0,02036 |
| 0,02036 |
| 0,02036 |
| 0,02066 |
| 0,02072 |
| 0,02081 |
| 0,02107 |
| 0,02115 |
| 0,02128 |
| 0,0214 |
| 0,0214 |
| 0,02167 |
| 0,02167 |
| 0,02167 |
| 0,02225 |
| 0,02236 |
| 0,02244 |
| 0,02255 |
| 0,02255 |
| 0,02255 |
| 0,02257 |
| 0,02257 |

| |
|---|
| 0,02272 |
| 0,02296 |
| 0,02348 |
| 0,0235 |
| 0,02353 |
| 0,02372 |
| 0,02378 |
| 0,02378 |
| 0,02387 |
| 0,02409 |
| 0,02484 |
| 0,02508 |
| 0,02535 |
| 0,02576 |
| 0,02606 |
| 0,02606 |
| 0,02606 |
| 0,02618 |
| 0,02618 |
| 0,02656 |
| 0,02656 |
| 0,02686 |
| 0,02705 |
| 0,02736 |
| 0,02736 |
| 0,02736 |
| 0,02849 |
| 0,02868 |
| 0,02929 |
| 0,02941 |
| 0,03059 |
| 0,03059 |
| 0,03059 |
| 0,03104 |
| 0,03104 |
| 0,03112 |
| 0,03123 |
| 0,03215 |
| 0,03272 |
| 0,03289 |
| 0,03306 |
| 0,03327 |
| 0,03354 |
| 0,03366 |

| |
|---|
| 0,03369 |
| 0,03369 |
| 0,03371 |
| 0,03383 |
| 0,03383 |
| 0,03388 |
| 0,0344 |
| 0,03444 |
| 0,03475 |
| 0,03482 |
| 0,03482 |
| 0,03482 |
| 0,03482 |
| 0,03497 |
| 0,03511 |
| 0,03547 |
| 0,03584 |
| 0,03589 |
| 0,03598 |
| 0,03598 |
| 0,0363 |
| 0,03652 |
| 0,03652 |
| 0,03687 |
| 0,03738 |
| 0,03765 |
| 0,03765 |
| 0,03801 |
| 0,03801 |
| 0,03837 |
| 0,03845 |
| 0,03845 |
| 0,03845 |
| 0,03845 |
| 0,03845 |
| 0,0385 |
| 0,03871 |
| 0,03919 |
| 0,03919 |
| 0,03919 |
| 0,03928 |
| 0,03928 |
| 0,03936 |
| 0,03955 |

| |
|---|
| 0,03997 |
| 0,04045 |
| 0,04045 |
| 0,04047 |
| 0,04047 |
| 0,04047 |
| 0,04062 |
| 0,04082 |
| 0,04133 |
| 0,04144 |
| 0,04147 |
| 0,04155 |
| 0,04155 |
| 0,04155 |
| 0,04155 |
| 0,04161 |
| 0,04203 |
| 0,04211 |
| 0,04214 |
| 0,04241 |
| 0,04242 |
| 0,04242 |
| 0,04242 |
| 0,04242 |
| 0,04251 |
| 0,04256 |
| 0,04402 |
| 0,04425 |
| 0,04425 |
| 0,04443 |
| 0,04443 |
| 0,04443 |
| 0,04443 |
| 0,04443 |
| 0,04447 |
| 0,04452 |
| 0,04487 |
| 0,04535 |
| 0,04573 |
| 0,04585 |
| 0,04716 |
| 0,04729 |
| 0,04729 |
| 0,04753 |

| |
|---|
| 0,04753 |
| 0,04768 |
| 0,04794 |
| 0,04844 |
| 0,04884 |
| 0,04884 |
| 0,04892 |
| 0,04892 |
| 0,04939 |
| 0,0495 |
| 0,04983 |
